# A comprehensive overview of the Chloroflexota community in wastewater treatment plants worldwide

**DOI:** 10.1101/2023.06.26.546502

**Authors:** Francesca Petriglieri, Zivile Kondrotaite, Caitlin Singleton, Marta Nierychlo, Morten K. D. Dueholm, Per H. Nielsen

## Abstract

Filamentous Chloroflexota are abundant in activated sludge wastewater treatment plants (WWTPs) worldwide and are occasionally associated with poor solid-liquid separation or foaming, but most of the abundant lineages remain undescribed. Here, we present a comprehensive overview of Chloroflexota abundant in WWTPs worldwide, using high-quality metagenome-assembled genomes (MAGs) and 16S rRNA amplicon data from 740 Danish and global WWTPs. Many novel taxa were described, encompassing 4 families, 13 genera and 29 novel species. These were widely distributed across most continents, influenced by factors such as climate zone and WWTP process design. Visualization by fluorescence *in situ* hybridization (FISH) confirmed their high abundances in many WWTPs based on the amplicon data and showed a filamentous morphology for nearly all species. Most formed thin and short trichomes integrated into the floc structure, unlikely to form the typical inter-floc bridging that hinders activated sludge floc settling. Metabolic reconstruction of 53 high-quality MAGs, representing most of the novel genera, offered further insights into their versatile metabolisms and suggested a primary role in carbon removal and involvement in nitrogen and sulfur cycling. The presence of glycogen reserves, detected by FISH-Raman microspectroscopy, seemed widespread across the phylum demonstrating that these bacteria likely utilize glycogen as an energy storage to survive periods with limited resources. This study gives a broad overview of the Chloroflexota community in global activated sludge WWTPs and improves our understanding of their roles in these engineered ecosystems.

**Importance:** Chloroflexota are often abundant members of the biomass in wastewater treatment plants (WWTPs) worldwide, typically with a filamentous morphology, forming the backbones of the activated sludge (AS) floc. However, their overgrowth can often cause operational issues connected to poor settling or foaming, impairing effluent quality and increases operational costs. Despite the importance, few Chloroflexota genera have been characterized so far. Here, we present a comprehensive overview of Chloroflexota abundant in WWTPs worldwide and an in-depth characterization of their morphology, phylogeny, and ecophysiology, obtaining a broad understanding of their ecological role in activated sludge.

## Introduction

Microorganisms belonging to the phylum Chloroflexota are frequently observed in the filamentous biomass in activated sludge (AS) wastewater treatment plants (WWTPs). They promote floc-formation by creating the backbone upon which other bacteria can attach (1–3). However, the uncontrolled overgrowth of specific genera, such as *Candidatus* Amarolinea, can also cause operational issues (4). According to the current taxonomic classification, based on 16S rRNA gene phylogeny (2), Chloroflexota found in WWTPs belong mainly to the classes Anaerolineae and Chloroflexia, with only a few cultured representatives mostly in the family *Anaerolineaceae* (2, 5–10).

Chloroflexota have a versatile facultative anaerobic metabolism, likely providing them with a competitive advantage in treatment plants with nitrogen removal and/or enhanced biological phosphorus removal (EBPR) systems characterized by alternating oxic and anoxic stages (2). The preferred substrates of known Chloroflexota filamentous species are carbohydrates, complex polymers such as cellulose, or amino acids (2). All described genera have the potential for fermentation of different products and some have the genes to carry out dissimilatory nitrate reduction or partial denitrification (11, 12), indicating a potential role in nitrogen removal from wastewater. The isolation of a nitrite-oxidizing bacterium, *Nitrolancea hollandica*, belonging to this phylum further supports their possible involvement in several steps in the nitrogen cycle (13).

Historically, filamentous bacteria in AS were identified using morphological features defined by specific staining methods and light microscopy (14–16), which often resulted in imprecise classification with little phylogenetic resolution (4). The introduction of high-throughput DNA sequencing and bioinformatics tools offered a breakthrough for the profiling of microbial communities. However, incomplete universal reference databases along with a lack of taxonomy for most uncultured lineages, including abundant Chloroflexota in WWTPs has hampered our ability to study these key organisms at lower taxonomic ranks (17). To improve the taxonomic resolution in microbial profiling studies, we introduced the Microbial Database for Activated Sludge (MiDAS), which includes a global, ecosystem-specific 16S rRNA gene reference database for wastewater treatment systems (MiDAS 4) (18). It also serves as a powerful tool for the design of genus- or species-specific fluorescence *in situ* hybridization (FISH) probes, which can subsequently be applied in combination with other techniques (e.g., microautoradiography or Raman microspectroscopy) for physiological characterization (1, 19). Furthermore, the recent retrieval of thousands of high quality (HQ) metagenome-assembled genomes (MAGs) (20, 21), together with the retrieval of several MAGs from Chloroflexota abundant in AS (12, 22–24), will allow us to obtain an improved overview of the phylogeny and role of Chloroflexota in the AS system.

Here, we present a comprehensive overview of the Chloroflexota abundant in WWTPs worldwide, using HQ MAGs and amplicon data from Danish and global WWTPs, in combination with the MiDAS 4 database. In total, we described 4 families, 13 genera and 29 novel species, which appeared to be widely distributed across most continents, and influenced by factors such as climate zone and WWTP process design. The design of specific genus-level FISH probes enabled investigation of their morphology, abundance, and spatial arrangement. Most of the novel Chloroflexota presented a typical filamentous morphology and demonstrated the presence of glycogen reserves, detected by FISH-Raman. Moreover, the annotation of 53 HQ MAGs, recently retrieved from Danish WWTPs (21), provided further insights into Chloroflexota functional potential and their involvement in nutrient cycling. This study represents a first fundamental milestone in the understanding of the ecological role of these microorganisms in the activated sludge microbial community.

## Materials and Methods

### Sampling and fixation

Sampling of AS was carried out within the Danish MiDAS survey (25) and the global MiDAS project (18). In short, fresh biomass samples from full-scale AS WWTPs were collected and either sent to Aalborg University (Danish MiDAS) or preserved in RNAlater and shipped to Aalborg University with cooling elements (Global MiDAS). Upon arrival, samples were stored at −20°C for sequencing workflows and fixed for FISH with 50% ethanol (final volume) or 4% PFA (final volume), as previously described (26).

### Community profiling using 16S rRNA gene amplicon sequencing

DNA extraction, sample preparation, and amplicon sequencing were performed as previously described (18, 25). Briefly, DNA was extracted using a custom plate-based extraction protocol based on the FastDNA spin kit for soil (MP Biomedicals). The protocol is available at www.midasfieldguide.org (aau_wwtp_dna_v.8.0). For Global MiDAS samples, RNAlater was removed by centrifugation and resuspension of the sample in 320 μL PBS. For Danish MiDAS samples, 160 µl of sample was mixed with 160 µl of PBS. All samples were transferred to Lysing Matrix E barcoded tubes and bead beating was performed in a FastPrep-96 bead beater (MP Biomedicals) (3 × 120 s, 1800 rpm, 2 min incubation on ice between beating). Community profiling was performed using 16S rRNA amplicon sequencing. V1-V3 16S rRNA gene regions were amplified using the 27F (AGAGTTTGATCCTGGCTCAG) (27) and 534R (ATTACCGCGGCTGCTGG) (28) primers, and the resulting amplicons were used in all the analyses. The V4 16S rRNA gene region was amplified using the 515F (50-GTGYCAGCMGCCGCGGTAA-30) (29) and 806R (50-GACTACNVGGGTWTCTAAT-30) (30) primers for comparison with the previous dataset. Data was analyzed using R (version 3.5.2) (31), RStudio software (32) and visualized using ampvis2 (version 2.7.5) (33) and ggplot2 (34). The Köppen-Geiger climate zone classification (35) was utilized to categorize the countries participating in the global MiDAS project (18). Details about the climate zones classification and the countries belonging to it can be found in **Table S1**.

### Phylogenetic analysis based on the 16S rRNA gene, FISH probe design and evaluation

Phylogenetic analysis of 16S rRNA gene sequences and design of FISH probes for the novel Chloroflexota were performed using the ARB software v.6.0.6 (36). A phylogenetic tree was calculated based on comparative analysis of aligned 16S rRNA gene sequences, retrieved from the MiDAS 4 database (18), using the maximum likelihood method and a 1000 – replicates bootstrap analysis. Coverage and specificity were evaluated and validated in silico with the MathFISH web tool for hybridization efficiencies of target and potentially weak non-target matches (37). When needed, unlabelled competitors and helper probes were designed. All probes were purchased from Biomers (Ulm, Germany), labelled with 6-carboxyfluorescein (6-Fam), indocarbocyanine (Cy3), or indodicarbocyanine (Cy5) fluorochromes.

### Fluorescence in situ hybridization (FISH), quantitative FISH (qFISH) and Raman microspectroscopy

FISH was performed as described by Daims et al. (2005) (38). Optimal formamide concentration for each novel FISH probe was determined after performing hybridization at different formamide concentrations in the range 0-70% (with 5% increments). The intensity of at least 50 cells was measured using ImageJ (39) software. Optimal hybridization conditions are described in **Table S2**. EUBmix (40, 41) was used to target all bacteria and NON-EUB (42) was used as a negative control for sequence independent probe binding. Quantitative FISH (qFISH) biovolume fractions of individual genera were calculated as a percentage area of the total biovolume, hybridizing with both EUBmix probes and specific probe. In case of specific probe not overlapping with EUBmix, a mix of EUBmix and CFXmix (43, 44), both labelled in Cy5, were used as a universal probe for total biomass coverage. qFISH analyses, performed using the Daime image analysis software (45), were based on 30 fields of view taken at 630× magnification. Microscopic analysis was performed with Axioskop epifluorescence microscope (Carl Zeiss, Germany) equipped with LEICA DFC7000 T CCD camera or a white light laser confocal microscope (Leica TCS SP8 X). Raman microspectroscopy was applied in combination with FISH to look for the storage polymers polyphosphate (poly-P), glycogen, and polyhydroxyalkanoates (PHA) as previously described (19).

### Genome phylogeny, annotation and metabolic reconstruction

A set of 1083 MAGs (NCBI BioProject PRJNA629478, Singleton et al., 2021), meeting the MIMAG HQ draft standards of full-length rRNA genes, completeness >90% and contamination <5% (46), was searched for Chloroflexota members using GTDB-Tk v2.1.0 (RefSeq release 207) ‘de_novo_wf’ pipeline (47). Species representatives, based on 95% average nucleotide identity clustering of the MAGs, and completeness and contamination estimates were determined from Singleton et al., (2021) (**SData 1**). A total of 53 Chloroflexota MAGs were identified. The phylogenetic maximum likelihood tree was created using the concatenated, trimmed alignment of the 120 single copy gene proteins from the GTDB-Tk de novo workflow, which included our MAGs and representative RefSeq genomes, as well as *Candidatus* (*Ca.*) Amarolinea aalborgensis. Three Cyanobacterota genomes (NCBI accession numbers: GCA_000317655, GCA_002813895, GCA_003566215) were used as an outgroup to root the tree. The ∼5000 amino acid alignment was used as input for IQ-TREE v2.1.2 (48), which was run using the WAG+G model and 1000x bootstrap iterations using UFBoot ultrafast bootstrap approximation. The tree was visualized in ARB v6.0.3 (36) to set the root using the outgroup Cyanobacterota, and exported for visualization and final aesthetic adjustments in iTOL v6.1.1 (49) and Inkscape v0.92. Pyani v0.2.11 (50) was used to determine average nucleotide identity (ANI).

Genomes were annotated as previously described (51). Briefly, the EnrichM v5.0 ‘annotate’ pipeline (github.com/geronimp/enrichM) was used to annotate the protein sequences of the genomes against the EnrichM v10 database, which included the KEGG (52) orthology (KO) number annotated Uniref100 database. Enrichm ‘classify’ --cutoff 1 was used to determine the presence of 100% complete KEGG modules, such as for transporters and glycolysis (SData2 and 3). Additionally, the MAGs were uploaded to the ‘MicroScope Microbial Genome Annotation & Analysis Platform’ (MAGE) (53) for manual inspection and cross-validation of KO annotations found using EnrichM.

## Results and discussion

### Phylogenetic evaluation of the Chloroflexota members abundant in WWTPs

The phylogenetic diversity of novel and well-known Chloroflexota abundant in global WWTPs was evaluated using a comparison of genome-based and 16S rRNA gene-based phylogenies to obtain a robust taxonomic assignment and to resolve potential discrepancies between 16S rRNA gene- and genome-based classification methods. The phylogenomic analysis (**Figure 1, Additional File 1**) revealed clustering into different novel families and genera, largely supported by 16S rRNA gene-based classification using the MiDAS 4 reference database (**Figure 2, Additional File 1**).

**Figure 1.**
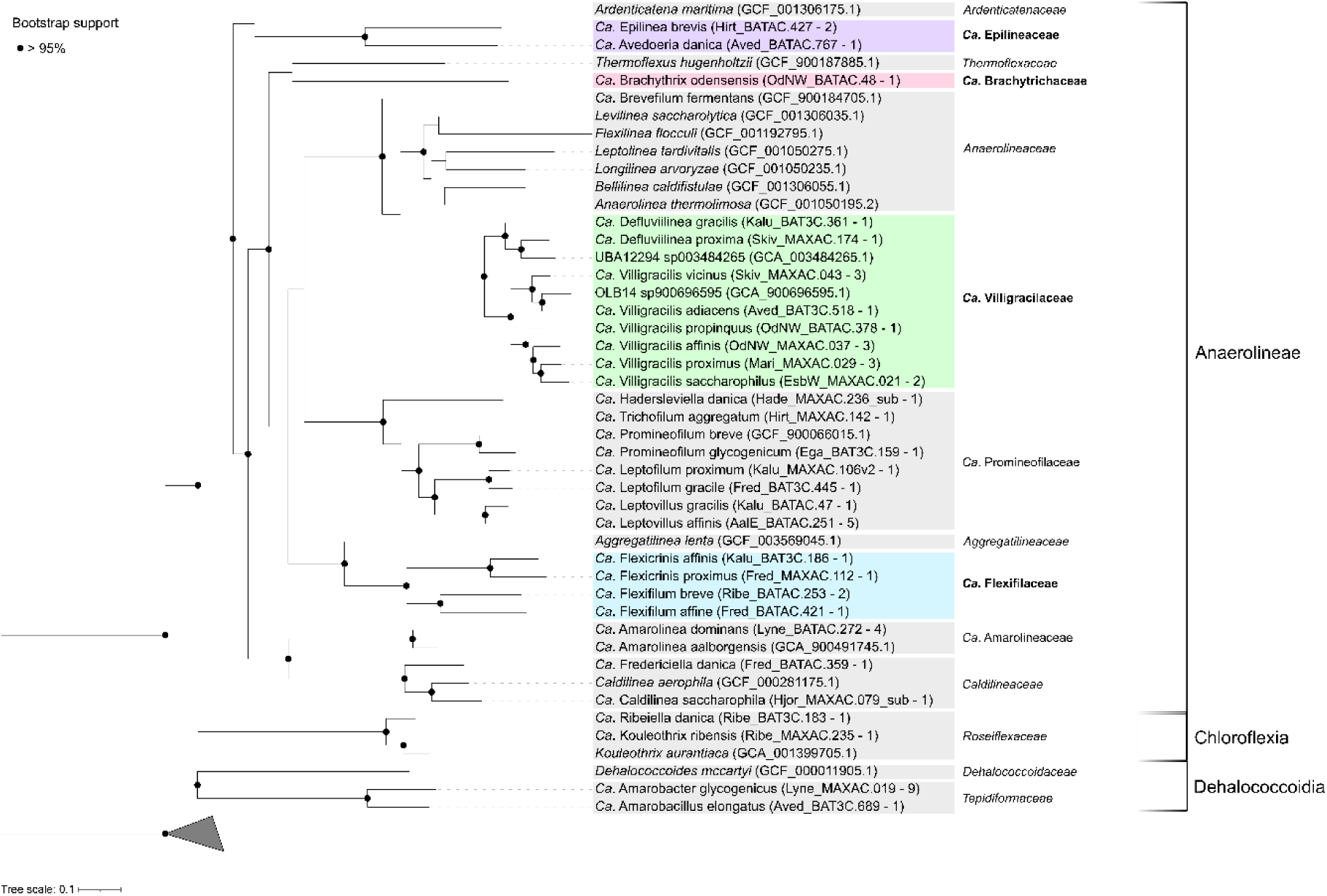
Phylogenetic genome tree of abundant Chloroflexota representatives. The maximum likelihood genome tree was created from the concatenated alignment of 120 single copy marker gene proteins trimmed to 5000 amino acids using the WAG+G model and 1000x UFBoot bootstrap iterations. Bootstrap support > 95% is shown by the solid black circles. Three Cyanobacterota genomes (NCBI accession numbers: GCA_000317655, GCA_002813895, GCA_003566215) were used as an outgroup to root the tree. For NCBI GenBank genome accession numbers see SData File 1. MAGs belonging to novel families are marked with colored boxes, while MAGs clustering with validly published families are marked with grey boxes. MAGs from species representatives are used to construct the tree and indicated between brackets, as well as the number of available MAGs for each lineage. The scale bar represents substitutions per amino acid base.

**Figure 2.**
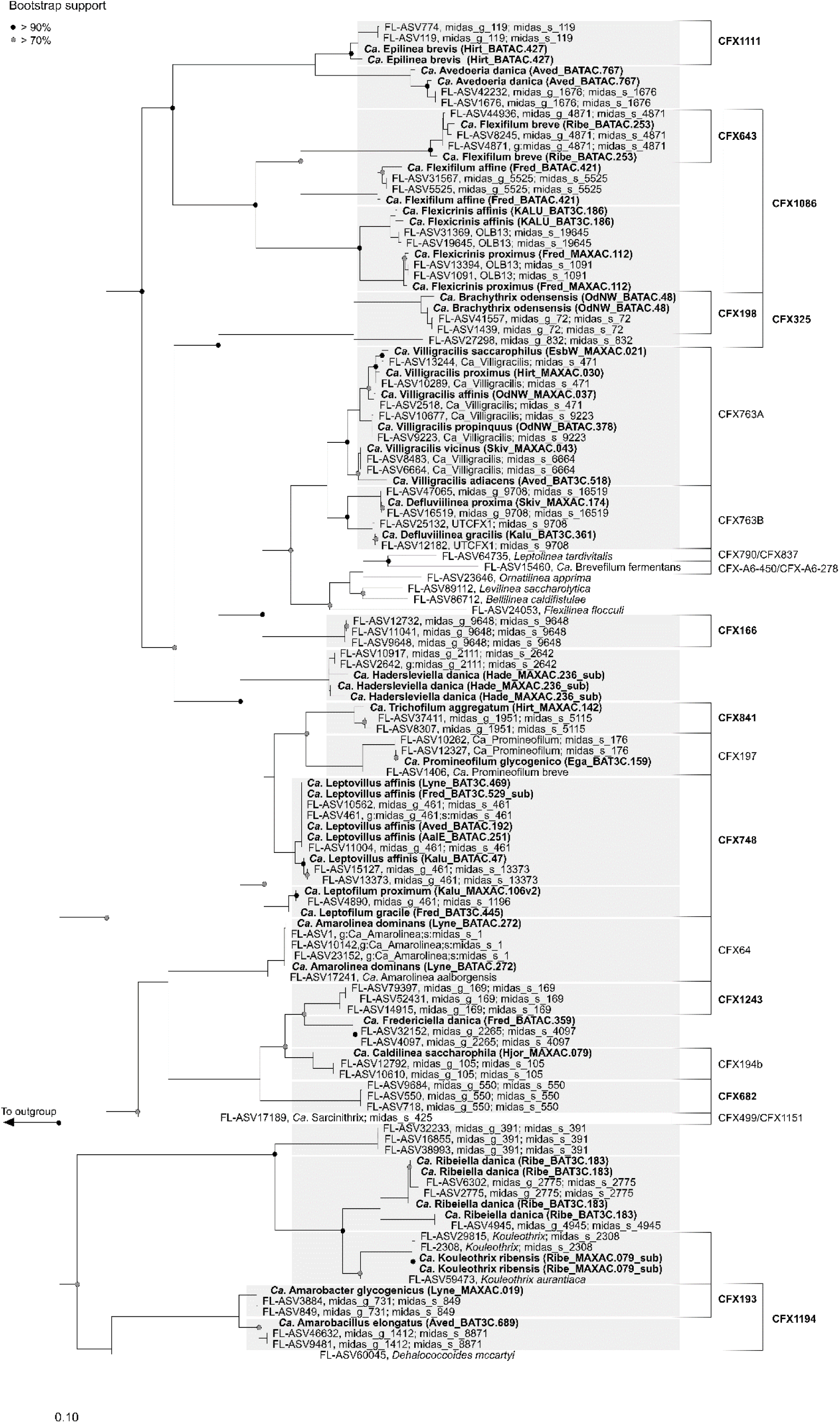
Maximum-likelihood (PhyML) 16S rRNA gene phylogenetic tree of Chloroflexota genera abundant in WWTPs. 16S rRNA gene sequences were retrieved from the global MiDAS 4 database or from the MAGs (bold). 16S rRNA gene sequences belonging to novel species representatives MAGs are indicated in bold blue. Grey boxes are used to indicate the taxonomy of novel species. The alignment used for the tree applied a 20% conservational filter to remove hypervariable positions, giving 1159 aligned positions. Coverage of the existing and designed in this study FISH probes is indicated with black brackets and is based on the MiDAS 4 database (18). Bootstrap values from 1000 re-samplings are indicated for branches with >70% (gray dot) and >90% (black) support. Species of the phylum Cyanobacteria were used as the outgroup. The scale bar represents substitutions per nucleotide base.

Of the 53 MAGs analyzed, several belonged to well-known families, such as *Ca*. Promineofilaceae (11 MAGs), *Ca*. Amarolineaceae (4 MAGs), *Caldilineaceae* (2 MAGs), *Roseiflexaceae* (2 MAGs), and *Tepidiformaceae* (10 MAGs), while the remaining represented the proposed new *Candidatus* families Epilineaceae (3 MAGs), Brachytrichaceae (1 MAG), Villigracilaceae (15 MAGs) and Flexifilaceae (5 MAGs). In many cases, the MAGs represented novel genera, as with *Ca*. Epilinea brevis (2 MAGs), *Ca*. Avedoeria danica (1 MAG), *Ca*. Brachythrix odensensis (1 MAG), *Ca.* Defluviilinea gracilis (1 MAG) and proxima (1 MAG), *Ca*. Hadersleviella danica (1 MAG), *Ca*. Trichofilum aggregatum (1 MAG), *Ca*. Leptofilum proximum (1 MAG) and gracile (1 MAG), *Ca*. Leptovillus gracilis (1 MAG) and affinis (5 MAGs), *Ca*. Flexicrinis affinis (1 MAG) and proximus (1 MAG), *Ca*. Flexifilum breve (1 MAG) and affine (1 MAG), *Ca*. Fredericiella danica (1 MAG), *Ca*. Ribeiella danica (1 MAG), *Ca*. Amarobacter glycogenicus (9 MAGs) and *Ca*. Amarobacillus elongatus (1 MAG). A few MAGs clustered together with genomes from known genera and represented new species, such as *Ca*. Promineofilum glycogenicum (1 MAG), *Ca*. Amarolinea dominans (4 MAGs), *Ca*. Caldilinea saccharophila (1 MAG, former *Ca*. Amarithrix (2)) and *Ca*. Kouleothrix ribensis (1 MAG). Interestingly, 13 MAGs corresponded to the well-known genus *Ca*. Villigracilis by comparison with the original 16S rRNA gene sequence used to define the genus (1), including the novel species *Ca*. Villigracilis vicinus (3 MAGs), adiacens (1 MAG), propinquus (1 MAG), affinis (3 MAGs), proximus (3 MAGs) and saccharophilus (2 MAGs). An in-depth analysis of the recently published *Ca*. Villigracilis nielsenii MAG (54) revealed its clustering within the *Ca*. Villigracilaceae family but outside of the *Ca*. Villigracilis genus (**Additional File 1, S Figure 3**), and we therefore propose to rename it *Candidatus* Manresella nielsenii, from the origin of the sludge (**Additional File 2**). A detailed summary of the phylogeny and supporting information for the phylogenetic analysis can be found in **Additional File 1**.

**Figure 3.**
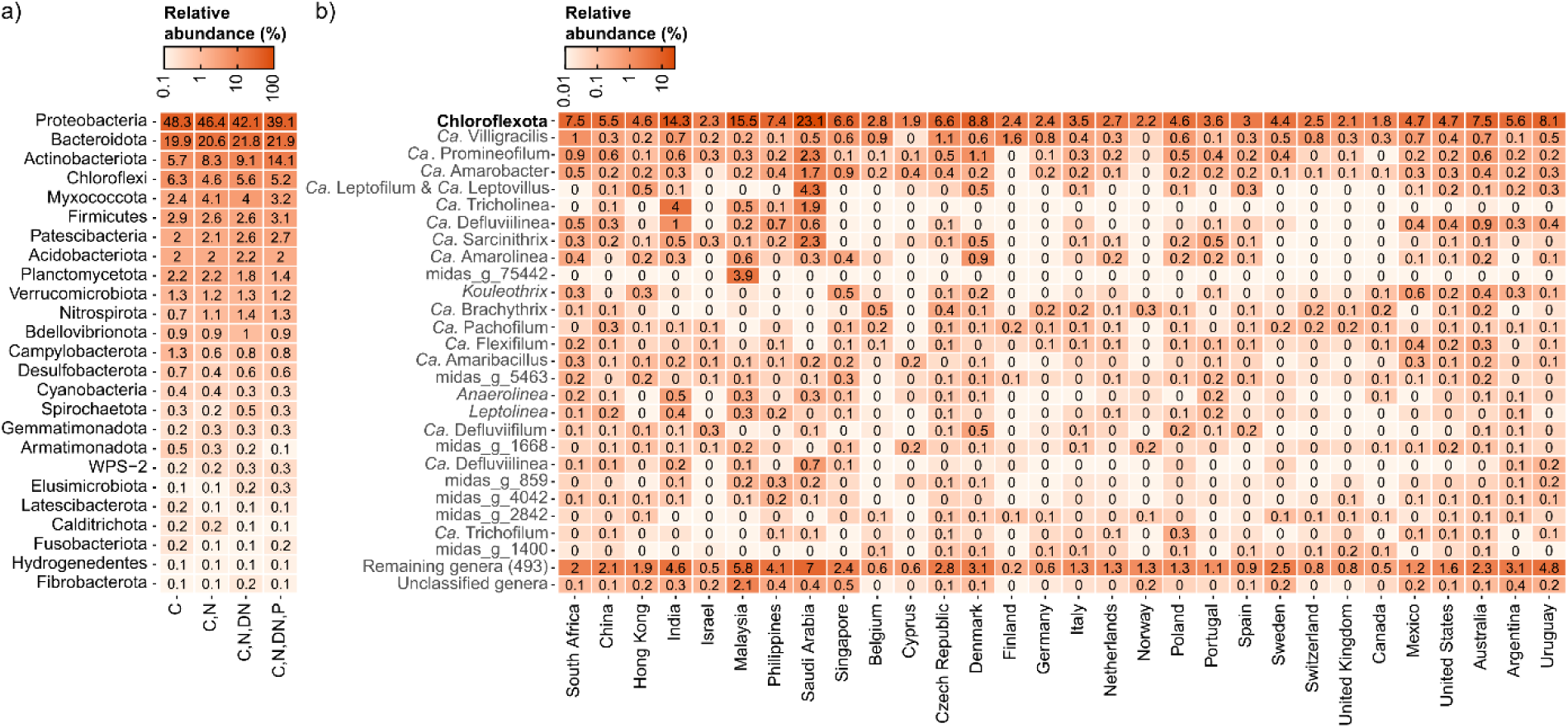
Global average relative read abundance of a) abundant phyla in WWTPs with different process designs and b) abundant Chloroflexota genera in different countries. Results are based on 929 activated sludge samples from 740 WWTPs. C, carbon removal; N, nitrification; DN, denitrification; P, biological P removal.

Despite absence of genomic information, 16S rRNA gene-based phylogeny (**Figure 2**), FISH probe design, and experimental analysis (see below) were possible for additional genera abundant in both global and Danish activated sludge samples. Therefore, we propose to rename the genera with placeholder names midas_g_391, midas_g_550, midas_g_9648 as *Ca*. Amarofilum, *Ca*. Pachofilum, and *Ca*. Tricholinea, respectively. 16S rRNA gene-based analysis using the MiDAS 4 database showed that midas_g_169 corresponded to *Ca*. Defluviifilum, as defined by Speirs et al., (2) and we suggest to adopt this name in future studies.

### Geographical distribution of Chloroflexota in global full-scale WWTPs

We analyzed the occurrence and diversity of both novel and well-known Chloroflexota genera abundant in global AS ecosystems, using data from the global MiDAS survey (18). On a global scale, the phylum Chloroflexota was the fourth most abundant phylum, making up 6.3% of the total reads (**Figure 3a**), similar to previous observations in Denmark (1), Spain (55), and Australia (56). Among the most abundant genera worldwide (**Figure 3b**), several well-known microorganisms appeared to be widespread, such as *Ca.* Villigracilis (1), *Ca.* Promineofilum (11), *Ca.* Sarcinithrix (1), *Ca*. Amarolinea (12), and *Kouleothrix* (57). All other abundant Chloroflexota were mainly undescribed, but potentially important for the process, such as the genus *Ca*. Defluviilinea (former UTCFX1), observed as part of the heterotrophic bacteria in anammox bioreactors (58, 59), or *Ca*. Flexifilum, belonging to the former family A4b, first identified in nitrifying-denitrifying industrial WWTPs (60). Interestingly, few novel genera were only found locally in individual countries, such as the genus midas_g_75442, present only in Malaysia (**Figure 3b)**.

*Ca.* Defluviifilum and *Ca*. Caldilinea saccharophila (former *Ca*. Amarithrix), together with the novel *Ca*. Amarofilum and *Ca*. Pachofilum, were commonly found in high abundance in Danish WWTPs, ranging from 0.1 to 1.1% but reaching up to 8% in some samples (**Figure S1**).

Examining global WWTP community composition enabled a deeper insight into factors affecting the occurrence of the different genera. We expected that the previously demonstrated slow-growth and facultative anaerobic metabolism of Chloroflexota species would favor their prevalence in long-sludge age WWTPs and WWTPs with biological N and P removal (2). *Ca*. Villigracilis, *Ca*. Promineofilum, and *Ca*. Amarobacter occurred in higher abundance in WWTPs with a complex process design involving both N and P removal, indicating a potential role in these processes (**Figure 4a**). The same genera also appeared to be influenced by the fraction of industrial wastewater (shown as the COD fraction in the influent), preferring low to medium content (<50%) of industrial wastewater (**Figure 4b**). This observation confirms previous findings, where filamentous Chloroflexota were detected in low abundance by FISH in industrial sludge (61). The differences in the Chloroflexota communities were more accentuated when considering the different climate zones (**Figure 4c**), with highest abundances of all genera observed in dry and temperate climates. Interestingly, some genera seemed to be specific to areas with hot temperatures, such as *Ca*. Defluviilinea and *Ca*. Tricholinea, dominant in the dry and arid climates, or *Ca*. Brachythrix, which seemed predominant in countries with polar climate (**Figure 4c-d**).

**Figure 4.**
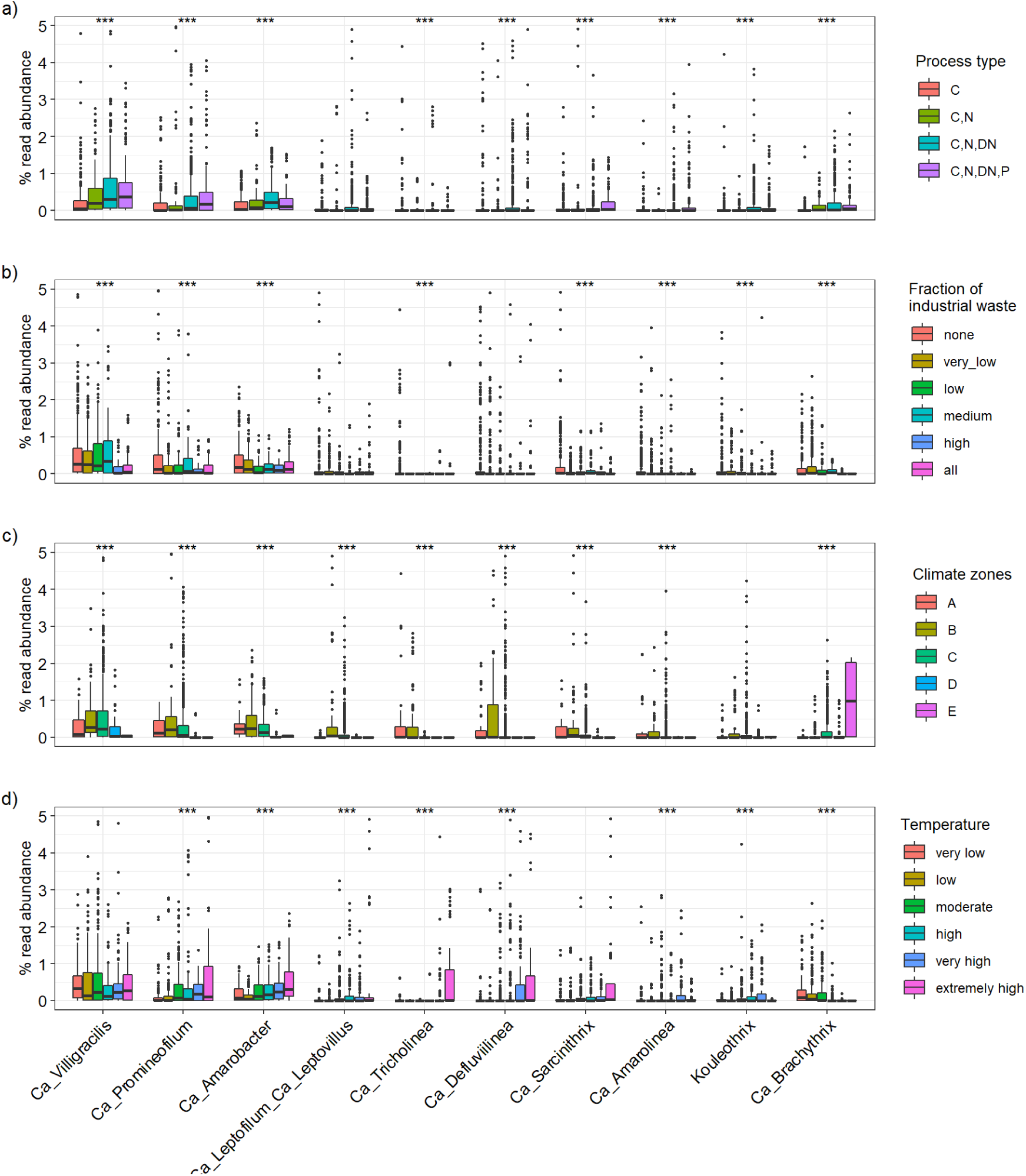
Distribution of selected abundant Chloroflexota genera across the world based on A) process type (C, 113 plants; C, N, 48 plants; C, N, DN, 208 plants, C, N, DN, P, 111 plants), (C, carbon removal; N, nitrification; DN, denitrification; P, biological P removal); B) fractions of industrial wastewater (0% - 169 plants; 0-10% - 105 plants; 11-30% - 67 plants; 31-50% - 41 plants; 51-99% - 20 plants; 100% - 40 plants); C) climate zones (A: tropical/megathermal climates, 29 plants; B: dry (desert and semi-arid) climates, 48 plants; C: temperate/mesothermal climates, 368 plants; D: continental/microthermal climates, 24 plants; E: polar climates, 2 plants), and D) different temperature ranges analyzed in process tanks (1-10.0°C, 43 plants; 10.1-15.0°C, 96 plants; 15.1-20.0°C, 112 plants; 20.1-25°C, 73 plants; 25.1-30.0°C, 48 plants; 30.1-38.0°C, 32 plants). Detailed information about the different climate zones and the countries belonging to each of them is in Table S1. Significant differences within individual groups are indicated by *** (Kruskal-Wallis test, p<0.001). For visualization purposes, samples with abundances higher than 5% are not shown in this figure.

### In situ characterization of Chloroflexota abundant in Danish and global WWTPs

We designed new FISH probes to target and characterize the abundant novel Chloroflexota genera *in situ* and re-evaluated the coverage and specificity of existing FISH probes (**Figure 2, Table 1**). *In silico* evaluation of the widely-applied CFXmix (43, 44) using the MiDAS 4 database showed good coverage of the phylum in the AS ecosystem, and it is recommended to be used in combination with EUB mix (40, 41) for better coverage of the Chloroflexota (**Table 1**). All but two of the genera investigated, *Ca.* Epilinea and *Ca.* Promineofilum, did hybridize with the EUB mix. *In silico* evaluation of previously published genus-level FISH targeting *Ca*. Amarolinea, *Ca*. Sarcinithrix, *Ca*. Promineofilum, and *Ca*. Villigracilis probes showed high specificity and hybridized with filaments of variable length and thickness (1, 3, 11, 61, 62) (**Figure 2, Table 1**). The majority of the novel Chloroflexota appeared to have the same conventional morphology (**Figure 5**), with filaments of different length and thickness often found in bundles inside the flocs, or sometimes creating inter-floc bridges (**Figure 5**; **Table 1**). Epiphytic bacteria were found on protruding filaments belonging to *Ca*. Flexifilum and other bacteria of the *Flexifilaceae* family, *Ca*. Trichofilum, *Ca.* Defluviifilum, *Ca*. Amarofilum and *Ca*. Pachofilum (**Figure 5**). Interestingly, the short filaments of *Ca*. Epilinea were found sometimes to be themselves attached to other Chloroflexota filaments (**Figure 5**). The surface adhesion mechanism of these microorganisms is still unclear, but the presence of pili has been previously reported for several isolates (63–65) and these appendages could mediate the adhesion process (66). The two genera from the order Dehalococcoidia, *Ca.* Amarobacter and *Ca.* Amarobacillus, were small rods.

**Figure 5.**
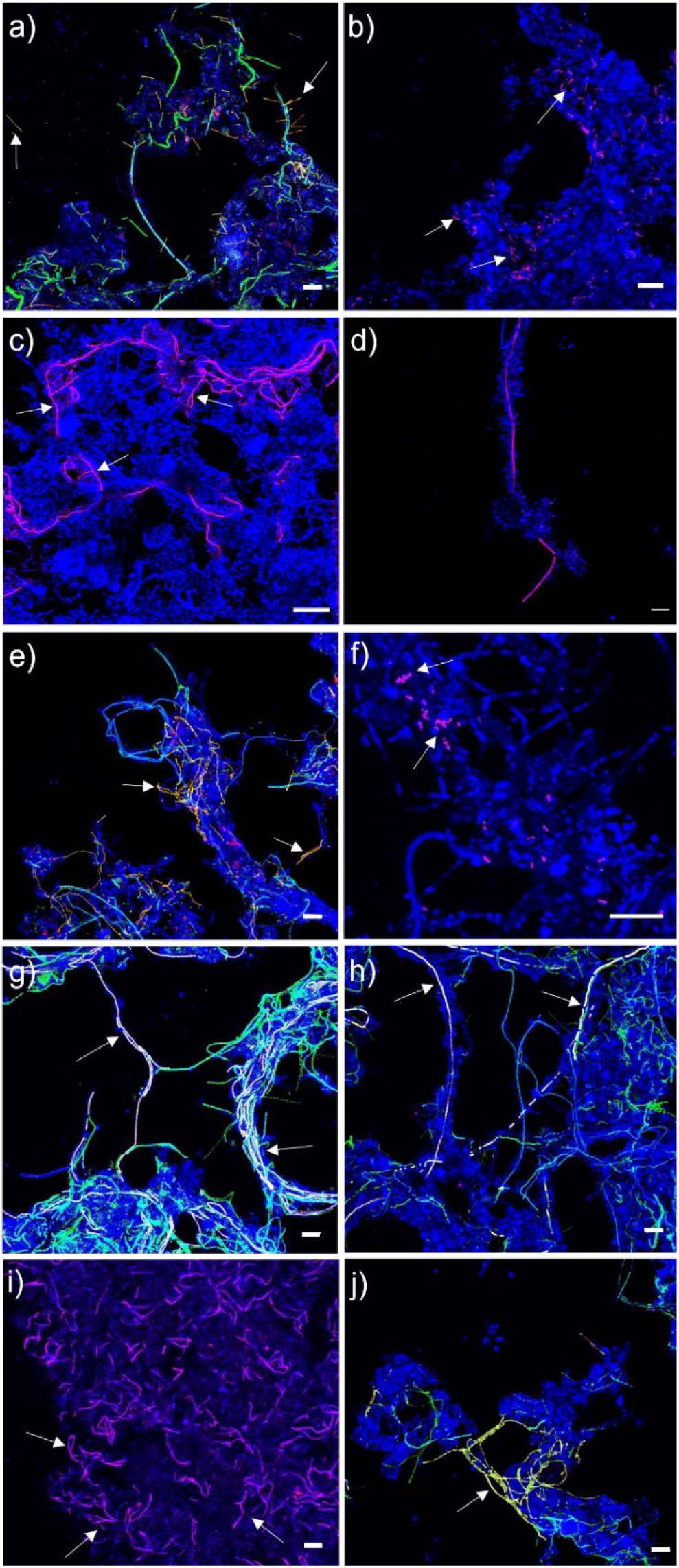
FISH micrographs of novel Chloroflexota genera in full-scale activated sludge. All bacteria were targeted with EUBmix (blue), in some micrographs CFXmix (green) was also applied to target most bacteria belonging to the Chloroflexota phylum. Specific probe targets: a) *Ca*. Epilinea (yellow); b) *Ca*. Brachythrix (magenta); c) *Ca*. Trichofilum (magenta); d) *Ca*. Flexifilum (magenta); e) - *Ca*. Leptofilum and *Ca*. Leptovillus (yellow); f) *Ca*. Amarobacter (magenta); g) *Ca*. Amarofilum (white); h) *Ca*. Pachofilum (white); i) *Ca*. Tricholinea (magenta); j) *Ca*. Defluviifilum (yellow). Scale bar is 20 µm.

**Table 1.**
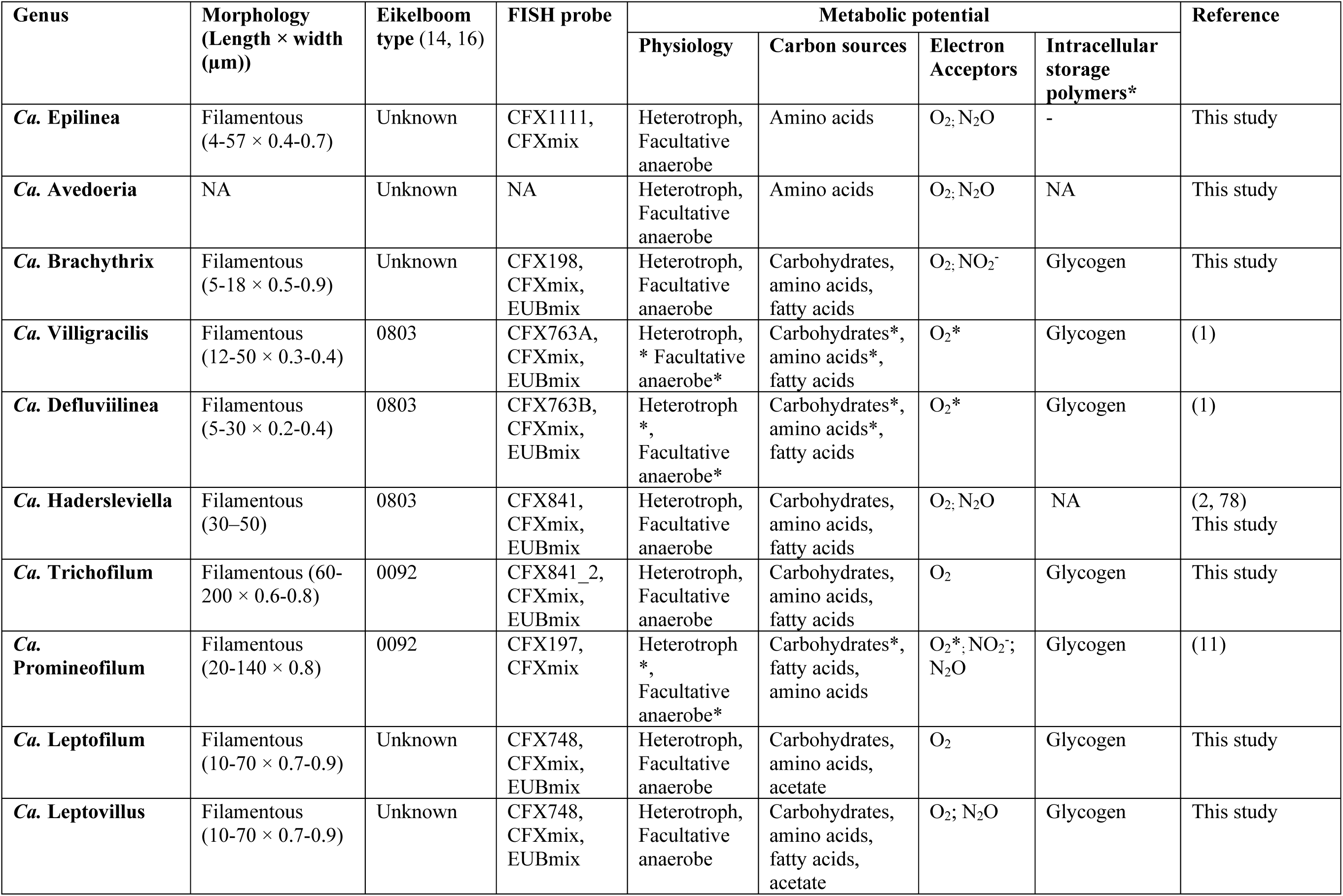
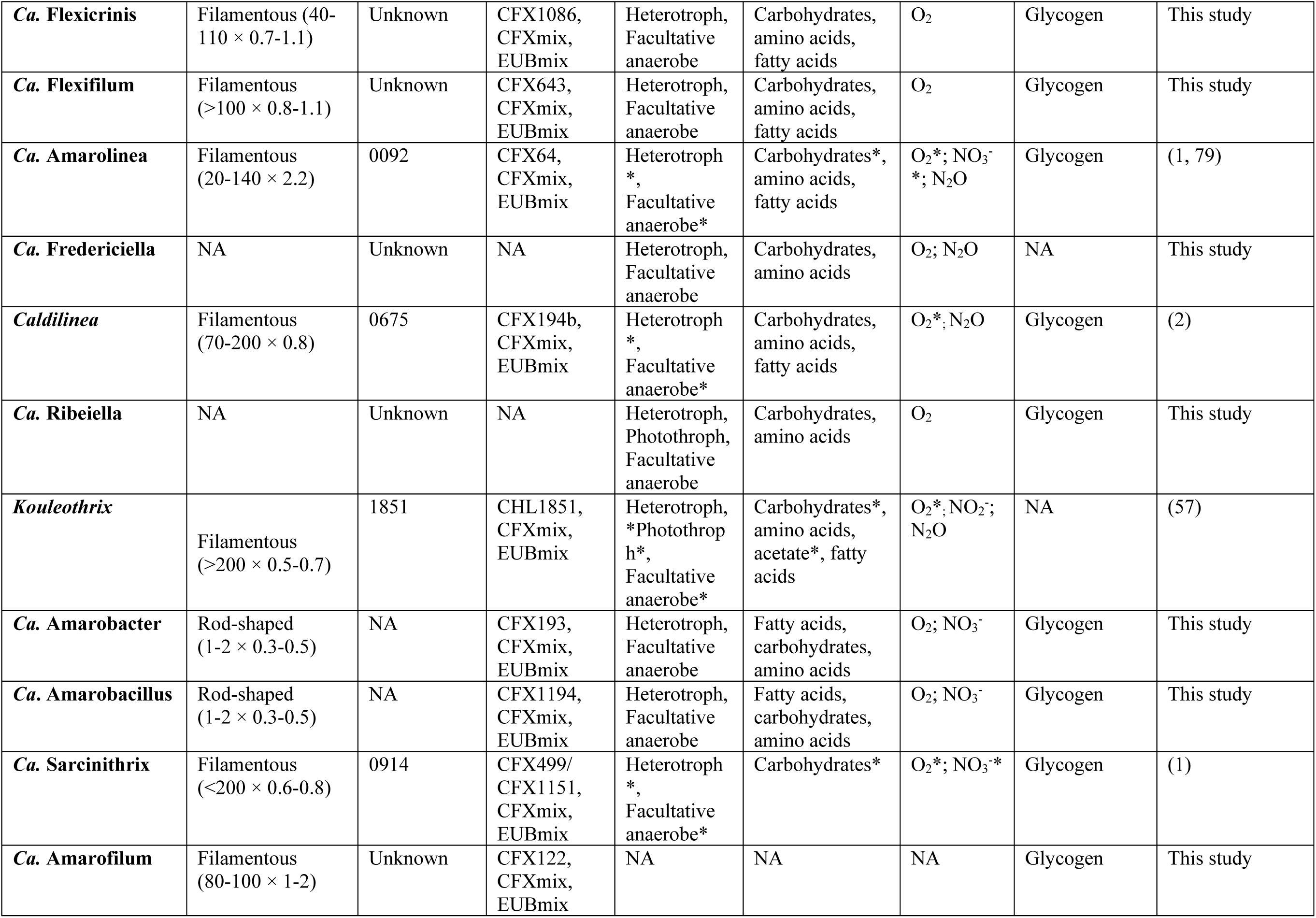
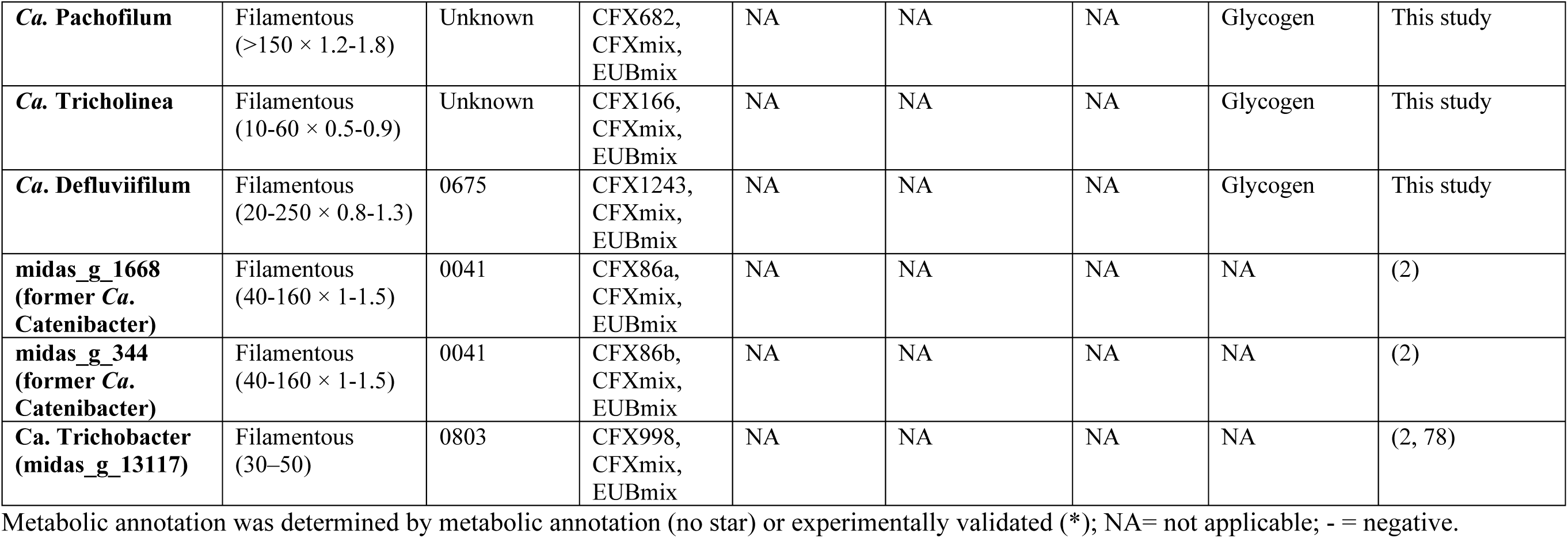
Summary table of morphology and ecophysiology of known Chloroflexota genera.

The FISH probes designed for the novel Chloroflexota genera were applied to Danish and, when possible, global activated sludge samples for FISH-based quantification (**Table S3**). Amplicon sequencing relative read abundances were in most cases very similar, while a few genera (*Ca*. Defluviifilum and *Ca*. Leptofilum) had slightly lower and some slightly higher (*Ca*. Amaribacillus, *Ca*. Amaribacter and *Ca*. Epilinea) read abundances compared to qFISH results. These differences are likely due to variations in cell size, extraction efficiency, and 16S rRNA gene copy number variation (67). The 16S rRNA gene copy numbers in the MAGs analyzed in this study varied from 1 to 5 (**SData 1**), which could lead to overestimation of some genera when using amplicon-based quantification (67). Primers are also known to introduce bias in amplicon sequencing (67). Therefore, the composition of the Chloroflexota community was compared using two commonly applied primer sets (the V1-V3 and V4 regions of the 16S rRNA gene sequences) (**Figure S2**). The overall relative abundance of the phylum Chloroflexota was comparable, but significant differences appeared in the relative abundances of specific genera **(Figure S2**). The V4-dataset showed lower relative abundances of some genera such as *Ca.* Promineofilum and *Ca.* Leptofilum, and almost complete disappearance of *Ca*. Villigracilis and *Ca*. Defluviilinea, which were likely not targeted by the V4 primer-set. These findings and the similarity of the abundances calculated by qFISH and V1-V3 amplicon sequencing confirmed that the V1-V3 primer set is more suited to encompass the diversity of the phylum Chloroflexota in AS systems. This is of particular importance if the potential effect on settling properties of these filamentous bacteria is evaluated using amplicon sequencing data.

To investigate the *in situ* physiology of the novel Chloroflexota genera, we performed Raman microspectroscopy in combination with the new FISH probes. This approach allows the detection of general cellular components, such as nucleic acids, membrane lipids, or proteins (68), as well as storage polymers important for the physiology of microorganisms involved in nitrogen or phosphorus removal in activated sludge (19, 51). Most of the Chloroflexota genera showed the presence of peaks characterizing common biological components, such as phenylalanine, nucleic acids, lipids, as well as a peak for glycogen, which most likely serves as a storage compound to survive periods with low energy sources (**Figure S3**). The presence of glycogen, which appears to be a conserved feature of the phylum, confirms the potential for glycogen accumulation proposed by our metabolic predictions (see below) and studies of other known Chloroflexota (11, 12, 63). Although granules of PHAs and poly-P have been identified previously in Chloroflexota isolates (63), these storage polymers were not detected *in situ*.

### Metabolic potential of the abundant Chloroflexota genera

Functional analysis of the 53 Chloroflexota MAGs revealed similar metabolisms to previously published models (11, 12). They exhibited a very versatile metabolism, revealing heterotrophic lifestyles with possible involvement in degradation of complex organic compounds and utilization of a wide selection of different sugars and amino acids as carbon sources (**Figure 6, SData 2 and 3**). Manual inspection of the MAGs confirmed that all the MAGs encoded pathways for full central carbon processing, through glycolysis, pentose phosphate pathway, and TCA cycle (**Figure 6, SData 2 and 3**).

**Figure 6.**
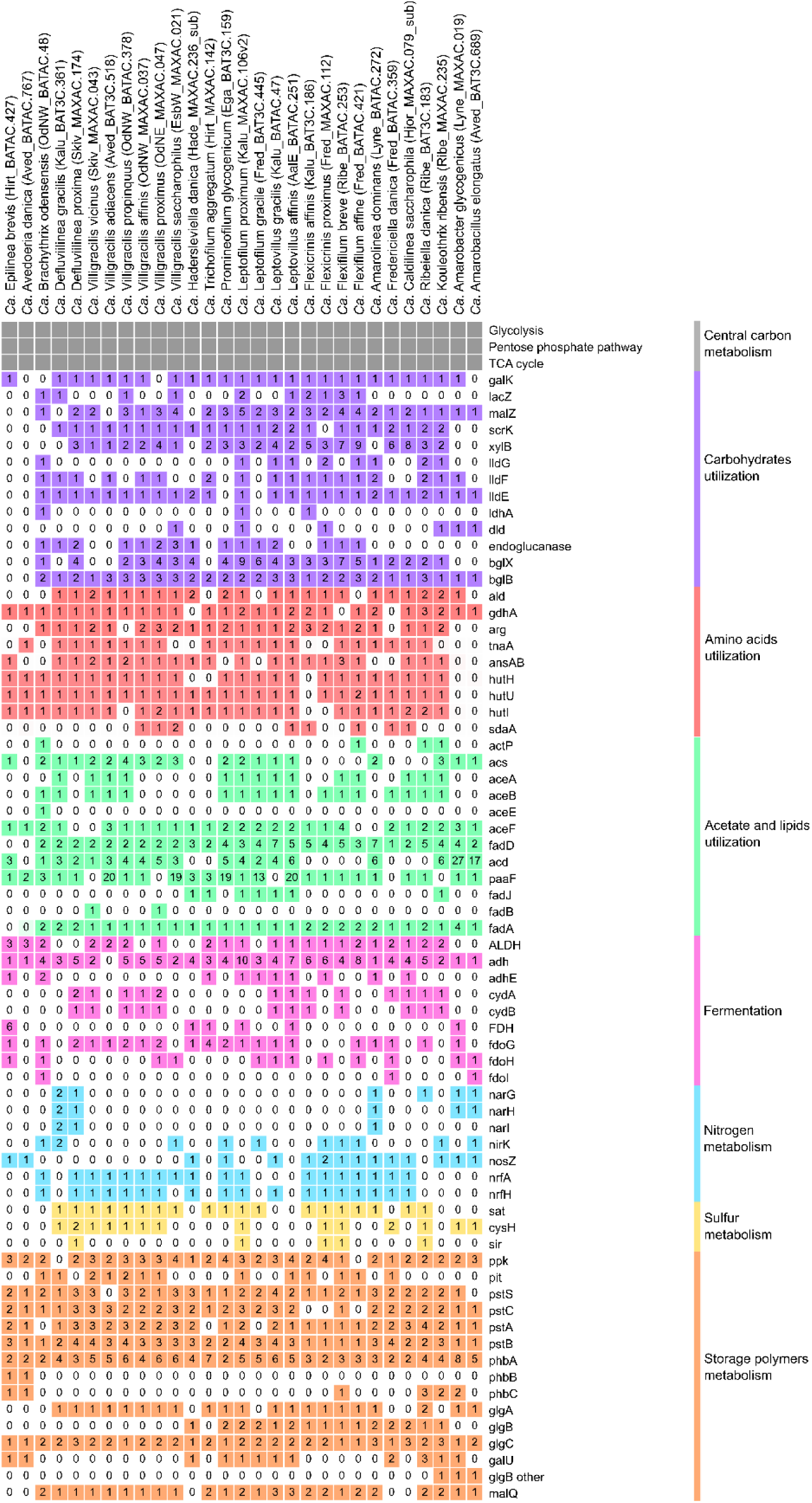
Basic functional potential of the Chloroflexota MAGs. For the full list of gene names and associated KO numbers see Data S2-3. The MAGs and genomes are ordered as in the genome tree in Figure 1. For simplicity, only MAGs from species representatives are showed. Numbers indicate gene copy number.

Aerobic uptake of different substrates is a shared trait exhibited by members of this phylum (1, 61). This widespread trait was confirmed in our MAGs, as ABC transporters for glucose/mannose (*gts*), fructose (*frc*), ribose (*rbs*), xylose (*xyl*), and glycerol 3-phosphate (*ugp* and *malK*) were predicted across all the MAGs (**SData 2 and 3**). Additionally, genes encoding a putative simple sugar ABC transporter and a multiple sugar transport system (*ggu*) were widely distributed across the MAGs (**SData 2 and 3**). Genes encoding for degradation of different sugars, such as galactose (*gal*), lactose (*lacZ*), sucrose (*malZ*), fructose (*scrK* or *fruK*) or xylose (*xyl*) were also present across all MAGs (**Figure 6, SData 2 and 3**). Additionally, the use of lactate as a carbon source in some of the MAGs was indicated by the presence of genes encoding for lactate permease (*lctP*) and lactate utilization (*lldG*, *lldF*, *lldE*, *ldhA*, *dld*) (**Figure 6, SData 2 and 3**). Several MAGs (17/53) also indicated the potential degradation of cellulose to glucose, with genes encoding endoglucanase and beta-glucosidase (*blgX* and/or *blgB*) **(Figure 6, SData 2 and 3**).

The presence of genes encoding for the transport of peptides (*dpp*), oligopeptides (*opp*), branched-chain amino acids (*liv*), and a putative polar amino acids transport system indicated that amino acids could be another important energy source (**SData 2 and 3**). Nearly all the MAGs encoded the alanine dehydrogenase (*ald*) for the oxidation of alanine to pyruvate under anoxic conditions. Furthermore, genes encoding the degradation of different amino acids were widespread in all the MAGs and included glutamate dehydrogenase (*gdhA*), arginase (*arg*), tryptophan *(tnaA*), asparagine (*ansA*), aromatic amino acids (*paaABCDE* and *paaK*), histidine (*hutHUI*, *ftcD,* and *fold*), and L-serine (*sdaA*). However, previous experimental studies did not show amino acid uptake under oxic or anoxic conditions in *Ca*. Villigracilis, *Ca*. Amarolinea, or *Ca*. Promineofilum (1, 11, 61), therefore further experimental validation is needed.

The acetate transporter gene (*actP*) was encoded only in two MAGs, belonging to the genera *Kouleothrix* and *Ca*. Brachythrix. The uptake of acetate and short-chain fatty acids has also not been experimentally confirmed in some of the abundant genera (1, 61, 69). Nonetheless, the presence of the gene *acs* was widespread in the MAGs, suggesting the potential use of acetate or fatty acids as carbon sources by these organisms, maybe deriving from internal pools (**Figure 6, SData 2 and 3**). The glyoxylate cycle was complete in part of the genomes (15/53), but only one MAG, belonging to *Ca*. Brachythrix, encoded the full potential for aerobic pyruvate oxidation to acetyl-CoA (*aceE* and *aceF*) (**Figure 6, SData 2 and 3**). Additionally, genes encoding for biosynthesis (*fadD*) or beta-oxidation of fatty acids (*acd*, *fadJ,* and *fadA*) were also widespread in the genera and identified in nearly all MAGs (**Figure 6, SData 2 and 3**). Interestingly, oxidation of long-chain fatty acids appeared to be one of the preferred energy sources for the bacteria belonging to the two non-filamentous genera in the order Dehalococcoidia, *Ca*. Amarobacter and *Ca*. Amarobacillus. They had a high copy number of genes involved in this pathway (*fadD*, *acd*, *paaF*, *fadJ, fadA*), comparable to those found in the well-known lipid user in AS systems *Ca*. Microthrix (70), indicating their potential specialization as lipid consumers (**Figure 6, SData 2 and 3**). Acetate use as carbon source is common in the Dehalococcoidia isolates (71, 72) and beta-oxidation has also been proposed as a potential metabolic route to obtain carbon and reducing equivalents by Dehaloccoidia found in marine sediments (73). Further analysis using transcriptomics and/or proteomics, could help to clarify the activity and expression level of these enzymes.

Anaerobic sugar uptake has previously been demonstrated *in situ* in several Chloroflexota genera and the potential for fermentation of substrates to acetoin or lactate was indicated in the genomes of *Ca*. Amarolinea and *Ca*. Promineofilum (1, 11, 12, 61). The potential for fermentation was encoded by most of the MAGs (*ALDH*, *adh* and *adhE*), with ethanol as a possible by-product (**Figure 6, SData 2 and 3**). Part of the MAGs (15/53) encoded genes for formate dehydrogenase (*fdh* and/or *fdoGHI*), potentially used to reduce formate produced during anaerobic fermentation, as suggested for *Ca*. Promineofilum breve (11).

Alternative electron acceptors under anaerobic conditions included nitrate, with potential dissimilatory nitrate reduction to nitrite (*narGHI*) identified in the four MAGs associated with the genus *Ca*. Amarolinea, as previously reported for *Ca*. Amarolinea aalborgensis (12) and in the two MAGs representing the new genus *Ca*. Defluviilinea (**Figure 6, SData 2 and 3**). Genes for nitrite reduction to nitric oxide (*nirK*) were present in 10/53 MAGs, while no MAGs encoded potential for nitric oxide reduction to nitrous oxide (*norBC*). 25 MAGs encoded genes (*nosZ*) for potential reduction of the latter to gaseous nitrogen (**Figure 6, SData 2 and 3**). The potential for dissimilatory nitrite reduction to ammonia (*nrfAH*) was widespread across the MAGs, similar to previous findings for *Ca*. Amarolinea aalborgensis (12) or *Ca*. Promineofilum breve (11), while no potential for nitrification was detected (**Figure 6, SData 2 and 3**). The potential for sulfate reduction to H_2_S through the assimilatory pathway (*sat*, *cysH* and *sir*) was also widespread across the MAGs (**Figure 6, SData 2 and 3**). Some Chloroflexota bacteria are described to be involved in the biogeochemical sulfur cycling (74–76).

Many bacteria that live under alternating oxic-anoxic conditions produce storage compounds, such as poly-P, glycogen, and PHA, which can be used in dynamic systems when environmental carbon or energy reserves are scarce. Examples are polyphosphate-accumulating organisms, involved in biological P removal in activated sludge (77). Genes indicating the potential for polyphosphate accumulation, such as the phosphate transporters (*pit*, *pstSCAB*) and the polyphosphate kinase (*ppk*) were widespread across the MAGs (**Figure 6, SData 2 and 3**). However, this storage compound was not detected in any Chloroflexota genus *in situ*. Only MAGs belonging to *Ca*. Epilinea and *Ca*. Avedoeria encoded the full potential for PHA accumulation (*phaABC*) (**Figure 6, SData 2 and 3**), but this intracellular polymer was not detected experimentally. Previous genome studies indicated the presence of glycogen as a storage compound in *Ca*. Amarolinea aalborgensis and *Ca*. Promineofilum breve (11, 12). The potential for glycogen biosynthesis was confirmed by the identification of the genes involved in the pathway (*glgABC*, *galU*, *glgP,* and *glgY*), as well as for its degradation (*malQ*) (**Figure 6, SData 2 and 3**), and experimentally validated in most of the genera.

The potential for carbon fixation through the enzyme RuBisCO and Calvin-Benson-Bassham cycle was also present in several MAGs (9/53), as previously observed for *Kouleothrix* and other Chloroflexota genera (2) (**SData 2 and 3**). Interestingly, the potential for bacteriochlorophyll biosynthesis and photorespiration were encoded by *Ca*. Ribeiella danica and *Ca*. Kouleothrix ribensis, similar to the isolates of the class Chloroflexia *Roseiflexus castenholzii* and *Chloroflexus aurantiacus* (**SData 2 and 3**) (23). As for the latter, *Ca*. Ribeiella and *Kouleothrix* present a fused form of the genes encoding for type II photosystem reaction centers (*pufLM*), recognized by manual inspection of the genomes (23). However, it is unclear if these pathways have a role in the activated sludge environment, where light is not easily accessible and the Calvin–Benson–Bassham pathway would be an energetically expensive alternative to the available organic carbon. Further analysis of the activity and expression level of these enzymes could help to clarify their role *in situ*.

### Ecological significance of Chloroflexota in activated sludge

This study provides a broad overview of the Chloroflexota abundant in AS WWTPs and contributes to recognizing their role in these systems. The phylum Chloroflexota encompasses a variety of different metabolisms, ranging from haloalkane-reducers (73, 80, 81), to anoxygenic photosynthetic microorganisms (23, 64, 82, 83), sponge holobionts (84–86), and extremophiles (87), all with roles in carbon, sulfur, and nitrogen cycling. It is therefore unsurprising that members of the Chloroflexota are present in the activated sludge ecosystem worldwide, although most lineages remain undescribed.

Our comprehensive approach, which includes the utilization of genome- and 16S rRNA gene-resolved phylogeny, allowed the identification of four novel families, 14 novel genera and 29 species, most of which are widely distributed across the continents and are seemingly influenced by factors such as climate zones, temperature, and WWTP process design. However, the variation in presence and abundance between countries and even between plants in the same country may also be largely influenced by immigration of microbial populations with the influent and seasonality (88, 89).

Generally, filamentous bacteria act as the backbone of activated sludge flocs, to which floc-forming bacteria attach and grow, typically as microcolonies. However, this beneficial role can change if the filaments are proliferating when the environmental conditions promote their growth, resulting in bulking and poor sludge-water separation (90). The Chloroflexota has often been associated with bulking episodes, consequently the morphological characterization of the novel genera is essential (2, 4). Most of the novel genera are, however, characterized by thin and short trichomes, some were short and attached to other filaments, so they did not form filamentous bridging which prevents the flocs from clustering as is known from *Ca.* Amarolinea and a few others, stressing that most are likely good for the floc formation. All filamentous members of Chloroflexota belonged to the orders Anaerolineae and Chloroflexia while the members from Dehalococcoidia, *Ca*. Amarobacter and *Ca*. Amarobacillus were rod shaped, showing that not all AS Chloroflexota are filamentous.

Confirming previous findings, Chloroflexota abundant in AS likely have a heterotrophic and facultative anaerobic lifestyle, which may explain their higher abundance in plants with more complex designs with oxic and anoxic conditions (1, 2, 11, 91). The different genera have versatile metabolisms, but with a seemingly important role in carbon cycling, with the potential ability to use various sugars and amino acids, but in some cases also lipids and acetate. They are assumed to have high hydrolytic activity, and the production of exo-enzymes for polysaccharide degradation has been previously shown *in situ* and suggests their importance in degrading the exopolymeric matrix, rather than soluble substrates (91, 92). Fermentation seems to be widespread across all the genera and likely sustains these organisms during anoxic conditions, as shown for some of the related isolates (7, 8, 10, 93). Despite being fermentative and facultative anaerobic microorganisms, Chloroflexota abundant in AS systems generally appear to die-off when the biomass is fed to anaerobic digesters, likely due to the substantial differences in environmental conditions, such as higher temperatures or salinity, while different Chloroflexota filaments seem to be predominant in the anaerobic digester environment (94–96). Their involvement in nitrogen, sulfur, and phosphorus cycling seems to be more limited. However, these organisms still have an important role in nutrient cycling if considered as active members of the bacterial consortia typical of the activated sludge floc, producing essential substrates that can be used by other microorganisms. Our study provides the foundation for future in-depth characterization of their physiology and relationships with other microorganisms. Following this, we believe gene expression and regulation studies are the next important steps forward in understanding the ecology of these process-critical bacteria.

### Formal taxonomic proposal

Etymologies and protologues for the novel proposed species are provided in Additional File1.

## Supporting information

Additional File 1 - Supplementary phylogenetic analysis

Additional File 2 - Taxonomic proposal and protologue tables

Supplementary Material - images and tables

## Acknowledgements

We would like to thank Prof. Aharon Oren for his assistance with the naming etymology. We also thank Claudia Etchebehere for the proposal for renaming *Candidatus* Manresella nielsenii. The project was funded by the Villum Foundation (Dark Matter grant13351) and the Poul Due Jensen Foundation (Microflora Danica).

## References

1. Nierychlo M, Milobedzka A, Petriglieri F, Mcilroy B, Nielsen PH, Mcilroy SJ. 2019. The morphology and metabolic potential of the Chloroflexi in full-scale activated sludge wastewater treatment plants. FEMS Microbiol Ecol 95:1–11.

2. Speirs LBM, Rice DTF, Petrovski S, Seviour RJ, Mcilroy SJ. 2019. The phylogeny, biodiversity, and ecology of the Chloroflexi in activated sludge. Front Microbiol 10:10:2015.

3. Speirs LBM, Dyson ZA, Tucci J, Seviour RJ. 2017. Eikelboom filamentous morphotypes 0675 and 0041 embrace members of the Chloroflexi: resolving their phylogeny, and design of fluorescence *in situ* hybridisation probes for their identification. FEMS Microbiol Ecol 93:1–13.

4. Nierychlo M, McIlroy SJ, Kucheryavskiy S, Jiang C, Ziegler AS, Kondrotaite Z, Stokholm-Bjerregaard M, Nielsen PH. 2020. *Candidatus* Amarolinea and *Candidatus* Microthrix are mainly responsible for filamentous bulking in Danish municipal wastewater treatment plants. Front Microbiol 11:1214.

5. Kale V, Björnsdóttir SH, Fridjónsson ÓH, Pétursdóttir SK, Ómarsdóttir S, Hreggvidsson GÓ. 2013. *Litorilinea aerophila* gen. nov., sp. nov., an aerobic member of the class Caldilineae, phylum Chloroflexi, isolated from an intertidal hot spring. Int J Syst Evol Microbiol 63:1149–1154.

6. Kawaichi S, Ito N, Kamikawa R, Sugawara T, Yoshida T, Sako Y. 2013. *Ardenticatena maritima* gen. nov., sp. nov., a ferric iron- and nitrate-reducing bacterium of the phylum “Chloroflexi” isolated from an iron-rich coastal hydrothermal field, and description of Ardenticatenia classis nov. Int J Syst Evol Microbiol 63:2992–3002.

7. Podosokorskaya OA, Bonch-Osmolovskaya EA, Novikov AA, Kolganova T V., Kublanov I V. 2013. *Ornatilinea apprima* gen. nov., sp. nov., a cellulolytic representative of the class Anaerolineae. Int J Syst Evol Microbiol 63:86–92.

8. Sekiguchi Y, Yamada T, Hanada S, Ohashi A, Harada H, Kamagata Y. 2003. *Anaerolinea thermophila* gen. nov., sp. nov. and *Caldilinea aerophila* gen. nov., sp. nov., novel filamentous thermophiles that represent a previously uncultured lineage of the domain bacteria at the subphylum level. Int J Syst Evol Microbiol 53:1843–1851.

9. Sun L, Toyonaga M, Ohashi A, Matsuura N, Tourlousse DM, Meng XY, Tamaki H, Hanada S, Cruz R, Yamaguchi T, Sekiguchi Y. 2016. Isolation and characterization of *Flexilinea flocculi* gen. Nov., sp. nov., a filamentous, anaerobic bacterium belonging to the class anaerolineae in the phylum Chloroflexi. Int J Syst Evol Microbiol 66:988–996.

10. Yamada T, Imachi H, Ohashi A, Harada H, Hanada S, Kamagata Y, Sekiguchi Y. 2007. *Bellilinea caldifistulae* gen. nov., sp. nov and *Longilinea arvoryzae* gen. nov., sp. nov., strictly anaerobic, filamentous bacteria of the phylum Chloroflexi isolated from methanogenic propionate-degrading consortia. Int J Syst Evol Microbiol 57:2299–2306.

11. McIlroy SJ, Karst SM, Nierychlo M, Dueholm MS, Albertsen M, Kirkegaard RH, Seviour RJ, Nielsen PH. 2016. Genomic and in situ investigations of the novel uncultured Chloroflexi associated with 0092 morphotype filamentous bulking in activated sludge. ISME J 10:2223–2234.

12. Andersen MH, Mcilroy SJ, Nierychlo M, Nielsen PH, Albertsen M. 2019. Genomic insights into *Candidatus* Amarolinea aalborgensis gen . nov ., sp . nov ., associated with settleability problems in wastewater treatment plants. Syst Appl Microbiol 42:77–84.

13. Sorokin DY, Vejmelkova D, Lücker S, Streshinskaya GM, Rijpstra WIC, Sinninghe Damsté JS, Kleerbezem R, van Loosdrecht M, Muyzer G, Daims H. 2014. *Nitrolancea hollandica* gen. nov., sp. nov., a chemolithoautotrophic nitrite-oxidizing bacterium isolated from a bioreactor belonging to the phylum Chloroflexi. Int J Syst Evol Microbiol 64:1859–1865.

14. Eikelboom DH, Geurkink B. 2002. Filamentous micro-organisms observed in industrial activated sludge plants. Water Sci Technol 46:535–542.

15. Eikelboom HD. 2000. Process control of activated sludge plants by microscopic investigation. IWA Publishing.

16. Jenkins D, Richard MG, Daigger GT. 1993. Manual on the causes and control of activated sludge bulking and foaming, 2nd Edn. Washington D.C: Lewis Publishers.

17. McIlroy SJ, Kirkegaard RH, McIlroy B, Nierychlo M, Kristensen JM, Karst SMSM, Albertsen M, Nielsen PHH. 2017. MiDAS 2.0: An ecosystem-specific taxonomy and online database for the organisms of wastewater treatment systems expanded for anaerobic digester groups. Database 2017:1–9.

18. Dueholm MKD, Nierychlo M, Andersen KS, Rudkjøbing V, Knutsson S, the Global MiDAS Consortium, Albertsen M, Nielsen PH. 2022. MiDAS 4: A global catalogue of full-length 16S rRNA gene sequences and taxonomy for studies of bacterial communities in wastewater treatment plants. Nat Commun 13:1908.

19. Fernando EY, McIlroy SJ, Nierychlo M, Herbst F-A, Petriglieri F, Schmid MC, Wagner M, Nielsen JL, Nielsen PH. 2019. Resolving the individual contribution of key microbial populations to enhanced biological phosphorus removal with Raman–FISH. ISME J 13:1933–1946.

20. Parks DH, Rinke C, Chuvochina M, Chaumeil P, Woodcroft BJ, Evans PN, Hugenholtz P, Tyson GW. 2017. Recovery of nearly 8,000 metagenome-assembled genomes substantially expands the tree of life. Nat Microbiol 2:1533–1542.

21. Singleton CM, Petriglieri F, Kristensen JM, Kirkegaard RH, Michaelsen TY, Andersen MH, Kondrotaite Z, Karst SM, Dueholm MS, Nielsen PH, Albertsen M. 2021. Connecting structure to function with the recovery of over 1000 high-quality metagenome-assembled genomes from activated sludge using long-read sequencing. Nat Commun 12:2009.

22. Kirkegaard RH, Dueholm MS, McIlroy SJ, Nierychlo M, Karst SM, Albertsen M, Nielsen PH. 2016. Genomic insights into members of the candidate phylum Hyd24-12 common in mesophilic anaerobic digesters. ISME J 10:2352–2364.

23. Ward LM, Hemp J, Shih PM, Mcglynn SE, Fischer WW. 2018. Evolution of phototrophy in the Chloroflexi phylum driven by horizontal gene transfer. Front Microbiol 9:1–16.

24. Dam HT, Vollmers J, Sobol MS, Cabezas A, Kaster AK. 2020. Targeted cell sorting combined with single cell genomics captures low abundant microbial dark matter with higher sensitivity than metagenomics. Front Microbiol 11:2020.

25. Nierychlo M, Andersen KS, Xu Y, Green N, Jiang C, Albertsen M, Dueholm MS, Nielsen PH. 2020. MiDAS 3: An ecosystem-specific reference database, taxonomy and knowledge platform for activated sludge and anaerobic digesters reveals species-level microbiome composition of activated sludge. Water Res 182:115955.

26. Nielsen JL. 2009. Protocol for fluorescence *in situ* hybridization (FISH) with rRNA-targeted oligonucleotides, p. 73-84. In FISH Handbook for biological wastewater treatment.

27. Lane DJ. 1991. 16S/23S rRNA sequencing., p. 115–175. In Stackebrandt, E. and Goodfellow, M., Eds., Nucleic acid techniques in bacterial ystematic. John Wiley and Sons.

28. Muyzer G, de Waal EC, Uitterlinden AG. 1993. Profiling of complex microbial populations by denaturing gradient gel electrophoresis analysis of polymerase chain reaction-amplified genes coding for 16S rRNA. Appl Environ Microbiol 59:695–700.

29. Parada AE, Needham DM, Fuhrman JA. 2016. Every base matters: Assessing small subunit rRNA primers for marine microbiomes with mock communities, time series and global field samples. Environ Microbiol 18:1403–1414.

30. Apprill A, Mcnally S, Parsons R, Weber L. 2015. Minor revision to V4 region SSU rRNA 806R gene primer greatly increases detection of SAR11 bacterioplankton. Aquat Microb Ecol 75:129–137.

31. R Core Team. 2020. R: A language and environment for statistical computing. R Foundation for Statistical Computing, Vienna, Austria.

32. RStudio Team. 2015. RStudio: Integrated Development Environment for R. Boston, MA.

33. Andersen KSS, Kirkegaard RH, Karst SM, Albertsen M. 2018. ampvis2: an R package to analyse and visualise 16S rRNA amplicon data. bioRxiv 299537.

34. Wickham H. 2009. ggplot2 - Elegant Graphics for Data AnalysisSpringer. Springer Science & Business Media.

35. Peel MC, Finlayson BL, McMahon TA. 2007. Updated world map of the Köoppen-Geiger climate classification. Hydrol Earth Syst Sci 11:1633–1644.

36. Ludwig W, Strunk O, Westram R, Richter L, Meier H, Yadhukumar A, Buchner A, Lai T, Steppi S, Jacob G, Förster W, Brettske I, Gerber S, Ginhart AW, Gross O, Grumann S, Hermann S, Jost R, König A, Liss T, Lüßbmann R, May M, Nonhoff B, Reichel B, Strehlow R, Stamatakis A, Stuckmann N, Vilbig A, Lenke M, Ludwig T, Bode A, Schleifer KH. 2004. ARB: A software environment for sequence data. Nucleic Acids Res 32:1363–1371.

37. Yilmaz LS, Parnerkar S, Noguera DR. 2011. MathFISH, a web tool that uses thermodynamics-based mathematical models for in silico evaluation of oligonucleotide probes for fluorescence *in situ* hybridization. Appl Environ Microbiol 77:1118–1122.

38. Daims H, Stoecker K, Wagner M. 2005. Fluorescence in situ hybridization for the detection of prokaryotes, p. 213–239. In Osborn, AM, Smith, CJ (eds.), Molecular Microbial Ecology. Taylor & Francis, New York.

39. Schneider CA, Rasband WS, Eliceiri KW. 2012. NIH Image to ImageJ: 25 years of image analysis. Nat Methods 9:671–675.

40. Amann RI, Binder BJ, Olson RJ, Chisolm SW, Devereux R, Stahl DA. 1990. Combination of 16S rRNA-targeted oligonucleotide probes with flow cytometry for analyzing mixed microbial populations. Appl Env Microbiol 56:1919–1925.

41. Daims H, Brühl A, Amann R, Schleifer KH, Wagner M. 1999. The domain-specific probe EUB338 is insufficient for the detection of all Bacteria: development and evaluation of a more comprehensive probe set. Syst Appl Microbiol 22:434–444.

42. Wallner G, Amann R, Beisker W. 1993. Optimizing fluorescent *in situ* hybridization with rRNA-targeted oligonucleotide probes for flow cytometric identification of microorganisms. Cytometry 14:136–143.

43. Björnsson L, Hugenholtz P, Tyson GW, Blackall LL. 2002. Filamentous Chloroflexi (green non-sulfur bacteria) are abundant in wastewater treatment processes with biological nutrient removal. Microbiology 148:2309–2318.

44. Gich F, Garcia-Gil J, Overmann J. 2002. Previously unknown and phylogenetically diverse members of the green nonsulfur bacteria are indigenous to freshwater lakes. Arch Microbiol 177:1–10.

45. Daims H, Lücker S, Wagner M. 2006. Daime, a novel image analysis program for microbial ecology and biofilm research. Environ Microbiol 8:200–213.

46. Bowers RM, Kyrpides NC, Stepanauskas R, Harmon-Smith M, Doud D, Reddy TBK, Schulz F, Jarett J, Rivers AR, Eloe-Fadrosh EA, Tringe SG, Ivanova NN, Copeland A, Clum A, Becraft ED, Malmstrom RR, Birren B, Podar M, Bork P, Weinstock GM, Garrity GM, Dodsworth JA, Yooseph S, Sutton G, Glöckner FO, Gilbert JA, Nelson WC, Hallam SJ, Jungbluth SP, Ettema TJG, Tighe S, Konstantinidis KT, Liu W-T, Baker BJ, Rattei T, Eisen JA, Hedlund B, McMahon KD, Fierer N, Knight R, Finn R, Cochrane G, Karsch-Mizrachi I, Tyson GW, Rinke C, Kyrpides NC, Schriml L, Garrity GM, Hugenholtz P, Sutton G, Yilmaz P, Meyer F, Glöckner FO, Gilbert JA, Knight R, Finn R, Cochrane G, Karsch-Mizrachi I, Lapidus A, Meyer F, Yilmaz P, Parks DH, Murat Eren A, Schriml L, Banfield JF, Hugenholtz P, Woyke T, Consortium TGS. 2017. Minimum information about a single amplified genome (MISAG) and a metagenome-assembled genome (MIMAG) of bacteria and archaea. Nat Biotechnol 35:725–731.

47. Chaumeil P-A, Mussig AJ, Hugenholtz P, Parks DH. 2020. GTDB-Tk: a toolkit to classify genomes with the Genome Taxonomy Database. Bioinformatics 36:1925–1927.

48. Minh BQ, Schmidt HA, Chernomor O, Schrempf D, Woodhams MD, von Haeseler A, Lanfear R. 2020. IQ-TREE 2: new models and efficient methods for phylogenetic inference in the genomic era. Mol Biol Evol 37:1530–1534.

49. Letunic I, Bork P. 2021. Interactive Tree Of Life (iTOL) v5: an online tool for phylogenetic tree display and annotation. Nucleic Acids Res https://doi.org/10.1093/nar/gkab301.

50. Pritchard L, Glover RH, Humphris S, Elphinstone JG, Toth IK. 2016. Genomics and taxonomy in diagnostics for food security: soft-rotting enterobacterial plant pathogens. Anal methods 8:12–24.

51. Petriglieri F, Singleton C, Peces M, Petersen JF, Nierychlo M, Nielsen PH. 2021. “*Candidatus* Dechloromonas phosphoritropha” and “*Ca*. D. phosphorivorans”, novel polyphosphate accumulating organisms abundant in wastewater treatment systems. ISME J 15:3605–3614.

52. Kanehisa M, Goto S. 2000. KEGG: Kyoto encyclopedia of genes and genomes. Nucleic Acids Res 28:27–30.

53. Vallenet D, Calteau A, Dubois M, Amours P, Bazin A, Beuvin M, Burlot L, Bussell X, Fouteau S, Gautreau G, Lajus A, Langlois J, Planel R, Roche D, Rollin J, Rouy Z, Sabatet V, Médigue C. 2020. MicroScope: an integrated platform for the annotation and exploration of microbial gene functions through genomic, pangenomic and metabolic comparative analysis. Nucleic Acids Res 48:D579– D589.

54. Bovio-Winkler P, Guerrero LD, Erijman L, Oyarzúa P, Suárez-Ojeda ME, Cabezas A, Etchebehere C. 2023. Genome-centric metagenomic insights into the role of Chloroflexi in anammox, activated sludge and methanogenic reactors. BMC Microbiol 23:45.

55. Celis M De, Belda I, Ortiz-álvarez R, Arregui L, Marquina D, Serrano S, Santos A. 2020. Tuning up microbiome analysis to monitor WWTPs ’ biological reactors functioning. Sci Rep 10:4079.

56. Nguyen LN, S CA, Hasan A, Bustamante H, Aurisch R, Lowrie R, Nghiem LD. 2019. Application of a novel molecular technique to characterise the effect of settling on microbial community composition of activated sludge. J Environ Manag 251:109594.

57. Beer M, Seviour EM, Kong Y, Cunningham M, Blackall LL, Y RJS. 2002. Phylogeny of the filamentous bacterium Eikelboom Type 1851, and design and application of a 16S rRNA targeted oligonucleotide probe for its fluorescence *in situ* identification in activated sludge. FEMS Microbiol Lett 207:179–183.

58. Lawson CE, Wu S, Bhattacharjee AS, Hamilton JJ, McMahon KD, Goel R, Noguera DR. 2017. Metabolic network analysis reveals microbial community interactions in anammox granules. Nat Commun 8:1–12.

59. Wang Y, Niu Q, Zhang X, Liu L, Wang Y, Chen Y, Negi M, Figeys D. 2019. Exploring the effects of operational mode and microbial interactions on bacterial community assembly in a one-stage partial-nitritation anammox reactor using integrated multi-omics. Microbiome 7:122.

60. Juretschko S, Loy A, Lehner A, Wagner M. 2002. The microbial community composition of a nitrifying-denitrifying activated sludge from an industrial sewage treatment plant analyzed by the full-cycle rRNA approach. Syst Appl Microbiol 25:84–99.

61. Kragelund C, Levantesi C, Borger A, Thelen K, Eikelboom D, Tandoi V, Kong Y, Van Der Waarde J, Krooneman J, Rossetti S, Thomsen TR, Nielsen PH. 2007. Identity, abundance and ecophysiology of filamentous Chloroflexi species present in activated sludge treatment plants. FEMS Microbiol Ecol 59:671–682.

62. Speirs L, Nittami T, McIlroy S, Schroeder S, Seviour RJ. 2009. Filamentous bacterium Eikelboom Type 0092 in activated sludge plants in Australia is a member of the phylum chloroflexi. Appl Environ Microbiol 75:2446–2452.

63. Gaisin VA, Kooger R, Grouzdev DS, Gorlenko VM, Pilhofer M. 2020. Cryo-Electron tomography reveals the complex ultrastructural organization of multicellular filamentous Chloroflexota (Chloroflexi) bacteria. Front Microbiol 11:1–15.

64. Gaisin VA, Kalashnikov AM, Sukhacheva M V., Namsaraev ZB, Barhutova DD, Gorlenko VM, Kuznetsov BB. 2015. Filamentous anoxygenic phototrophic bacteria from cyanobacterial mats of Alla hot springs (Barguzin Valley, Russia). Extremophiles 19:1067–1076.

65. Fukushima SI, Morohoshi S, Hanada S, Matsuura K, Haruta S. 2016. Gliding motility driven by individual cell-surface movements in a multicellular filamentous bacterium *Chloroflexus aggregans*. FEMS Microbiol Lett 363:1–5.

66. Tomich M, Planet PJ, Figurski DH. 2007. The tad locus: postcards from the widespread colonization island. Nat Rev Microbiol 5:363–375.

67. Albertsen M, Karst SM, Ziegler AS, Kirkegaard RH, Nielsen PH. 2015. Back to basics - The influence of DNA extraction and primer choice on phylogenetic analysis of activated sludge communities. PLoS One 10:1–15.

68. Hao L, McIlroy SJ, Kirkegaard RH, Karst SM, Fernando WEY, Aslan H, Meyer RL, Albertsen M, Nielsen PH, Dueholm MS. 2018. Novel prosthecate bacteria from the candidate phylum Acetothermia. ISME J 12:2225–2237.

69. Kindaichi T, Nierychlo M, Kragelund C, Nielsen JL, Nielsen PH. 2013. High and stable substrate specificities of microorganisms in enhanced biological phosphorus removal plants. Environ Microbiol 15:1821–1831.

70. Nierychlo M, Singleton CM, Petriglieri F, Thomsen L, Petersen JF, Peces M, Kondrotaite Z, Dueholm MS, Nielsen PH. 2021. Low global diversity of *Candidatus Microthrix*, a troublesome filamentous organism in full-scale WWTPs. Front Microbiol 12:690251.

71. Löffler FE, Yan J, Ritalahti KM, Adrian L, Edwards EA, Konstantinidis KT, Müller JA, Fullerton H, Zinder SH, Spormann AM. 2013. *Dehalococcoides mccartyi* gen. nov., sp. nov., obligately organohalide-respiring anaerobic bacteria relevant to halogen cycling and bioremediation, belong to a novel bacterial class, Dehalococcoidia classis nov., order Dehalococcoidales ord. nov. and famil. Int J Syst Evol Microbiol 63:625–635.

72. Kochetkova T V., Zayulina KS, Zhigarkov VS, Minaev N V., Chichkov BN, Novikov AA, Toshchakov S V., Elcheninov AG, Kublanov I V. 2020. *Tepidiforma bonchosmolovskayae* gen. nov., sp. nov., a moderately thermophilic Chloroflexi bacterium from a chukotka hot spring (arctic, Russia), representing a novel class, Tepidiformia, which includes the previously uncultivated lineageOLB14. Int J Syst Evol Microbiol 70:1192–1202.

73. Wasmund K, Schreiber L, Lloyd KG, Petersen DG, Schramm A, Stepanauskas R, Jørgensen BB, Adrian L. 2014. Genome sequencing of a single cell of the widely distributed marine subsurface Dehalococcoidia, phylum Chloroflexi. ISME J 8:383–397.

74. Vigneron A, Cruaud P, Culley AI, Couture RM, Lovejoy C, Vincent WF. 2021. Genomic evidence for sulfur intermediates as new biogeochemical hubs in a model aquatic microbial ecosystem. Microbiome 9:1–14.

75. Wasmund K, Mußmann M, Loy A. 2017. The life sulfuric: microbial ecology of sulfur cycling in marine sediments. Environ Microbiol Rep 9:323–344.

76. Mehrshad M, Rodriguez-Valera F, Amoozegar MA, López-García P, Ghai R. 2018. The enigmatic SAR202 cluster up close: shedding light on a globally distributed dark ocean lineage involved in sulfur cycling. ISME J 12:655–668.

77. Nielsen PH, Mcilroy SJ, Albertsen M. 2019. Re-evaluating the microbiology of the enhanced biological phosphorus removal process. Curr Opin Biotechnol 57:111–118.

78. Speirs LBM, Tucci J, Seviour RJ. 2015. The activated sludge bulking filament Eikelboom morphotype 0803 embraces more than one member of the Chloroflexi. FEMS Microbiol Ecol 91:fiv100.

79. Andersen MH, Mcilroy SJ, Nierychlo M, Nielsen PH, Albertsen M. 2020. Genomic insights into *Candidatus* Amarolinea aalborgensis gen. nov., sp. nov., associated with settleability problems in wastewater treatment plants. Syst Appl Microbiol 42:77–84.

80. Yan J, Rash BA, Rainey FA, Moe WM. 2009. Isolation of novel bacteria within the Chloroflexi capable of reductive dechlorination of 1,2,3-trichloropropane. Environ Microbiol 11:833–843.

81. Hug LA, Castelle CJ, Wrighton KC, Thomas BC, Sharon I, Frischkorn KR, Williams KH, Tringe SG, Banfield JF. 2013. Community genomic analyses constrain the distribution of metabolic traits across the Chloroflexi phylum and indicate roles in sediment carbon cycling. Microbiome 1:22.

82. Ward LM, Bertran E, Johnston DT. 2021. Expanded genomic sampling refines current understanding of the distribution and evolution of sulfur metabolisms in the Desulfobulbales. Front Microbiol 12:2021.

83. Ward LM, Li-Hau F, Kakegawa T, McGlynn SE. 2021. Complex history of aerobic respiration and phototrophy in the Chloroflexota class Anaerolineae revealed by high-quality draft genome of *Ca*. Roseilinea mizusawaensis AA3_104. Microbes Environ 36:ME21020.

84. Robbins SJ, Song W, Engelberts JP, Glasl B, Slaby BM, Boyd J, Marangon E, Botté ES, Laffy P, Thomas T, Webster NS. 2021. A genomic view of the microbiome of coral reef demosponges. ISME J 15:1641–1654.

85. Pita L, Rix L, Slaby BM, Franke A, Hentschel U. 2018. The sponge holobiont in a changing ocean: from microbes to ecosystems. Microbiome 6:46.

86. Loureiro C, Galani A, Gavriilidou A, Chaib de Mares M, van der Oost J, Medema MH, Sipkema D. 2022. Comparative metagenomic analysis of biosynthetic diversity across sponge microbiomes highlights metabolic novelty, conservation, and diversification. mSystems 7 :e0035722.

87. Zhou Z, John E St., Anantharaman K, Reysenbach A-L. 2022. Global patterns of diversity and metabolism of microbial communities in deep-sea hydrothermal vent deposits. Microbiome 10:241.

88. 88. Dottorini G, Michaelsen TY, Kucheryavskiy S, Andersen KS, Kristensen JM, Peces M, Wagner DS, Nierychlo M, Nielsen PH. 2021. Mass-immigration determines the assembly of activated sludge microbial communities. Proc Natl Acad Sci 118:e2021589118.

89. Peces M, Dottorini G, Nierychlo M, Andersen KS, Dueholm MKD, Nielsen PH. 2022. Microbial communities across activated sludge plants show recurring species-level seasonal patterns. ISME Commun 2:18.

90. Shi HX, Wang J, Liu SY, Guo JS, Fang F, Chen YP, Yan P. 2022. New insight into filamentous sludge bulking: potential role of AHL-mediated quorum sensing in deteriorating sludge floc stability and structure. Water Res 212:118096.

91. Kragelund C, Levantesi C, Borger A, Thelen K, Eikelboom D, Tandoi V, Kong Y, Krooneman J, Larsen P, Thomsen TR, Nielsen PH. 2008. Identity, abundance and ecophysiology of filamentous bacteria belonging to the Bacteroidetes present in activated sludge plants. Microbiology 154:886– 894.

92. Yamada T, Sekiguchi Y, Imachi H, Kamagata Y, Ohashi A, Harada H. 2005. Diversity, localization, and physiological properties of filamentous microbes belonging to Chloroflexi subphylum I in mesophilic and thermophilic methanogenic sludge granules. Appl Environ Microbiol 71:7493–7503.

93. Yamada T, Sekiguchi Y, Hanada S, Imachi H, Ohashi A, Harada H, Kamagata Y. 2006. *Anaerolinea thermolimosa* sp. nov., *Levilinea saccharolytica* gen. nov., sp. nov. and *Leptolinea tardivitalis* gen. nov., sp. nov., novel filamentous anaerobes, and description of the new classes Anaerolineae classis nov. and Caldilineae classis nov. in the . Int J Syst Evol Microbiol 56:1331–1340.

94. Bovio-Winkler P, Cabezas A, Etchebehere C. 2021. Database mining to unravel the ecology of the phylum Chloroflexi in methanogenic full scale bioreactors. Front Microbiol 11:2020.

95. Petriglieri F, Nierychlo M, Nielsen PH, Jon S, Id M. 2018. *In situ* visualisation of the abundant Chloroflexi populations in full-scale anaerobic digesters and the fate of immigrating species. PLoS ONE 13: e0206255 .

96. McIlroy SJ, Kirkegaard RH, Dueholm MS, Fernando E, Karst SM, Albertsen M, Nielsen PH. 2017. Culture-independent analyses reveal novel Anaerolineaceae as abundant primary fermenters in anaerobic digesters treating waste activated sludge. Front Microbiol 8:1134.

