## Additional File 1 - Supplementary phylogenetic analysis for "A comprehensive overview of the Chloroflexota community in wastewater treatment plants worldwide"

Of the 53 MAGs analyzed, three MAGs were classified into an undescribed family (UCB3) and order (UCB3) within the class Anaerolineae by the Genome Taxonomy Database (GTDB) GTDB-Tk tool, with order o__UCB3 and family f__UCB3 (**Table 1**). Phylogenomic analysis, supported also by our additional ANI analysis (**Figure 1)** and 16S rRNA gene- based classification, suggested the division in two novel genera and species, here called *Ca*. Epilinea brevis (2 MAGs) and *Ca*. Avedoeria danica (1 MAG). The names of *Ca*. Epilineaceae and Epiliniales are proposed for this family and order, as we were able to visualize this genus in situ. One MAG was distantly related to the isolate genome *Thermoflexus hugenholtzii*,and belongs to the order *Thermoflexales* and undescribed family f__J036 (**Table 1**). As the classification in a new lineage was also confirmed by 16S rRNA gene-based phylogeny, we propose to rename it as *Ca*. Brachythrix odensensis. The name of *Ca*. Brachytrichaceae is proposed for the family.

GTDB-Tk classified 16 MAGs as OLB14 (14 MAGs) and g_UBA12294 (2 MAGs), both genera belonging to the class Anaerolineae and the family f__envOPS12, originally identified in anammox granules (1, 2). This classification was also confirmed by 16S rRNA gene-based analysis, with 6 species within the *Ca*. Villigracilis genus (*Ca*. Villigracilis vicinus, *Ca*. Villigracilis adiacens, *Ca*. Villigracilis propinquus, *Ca.* Villigracilis affinis, *Ca*. Villigracilis proximus and *Ca*. Villigracilis saccharophilus) and 2 species representing the novel genus *Ca*. Defluviilinea (for which we propose the names *Ca*. Defluviilinea gracilis and *Ca*. Defluviilinea proxima) (**Table 1**). Supplementary ANI analysis (**Figure 2**), suggests the possibility of four genera, though the recovery of more HQ MAGs will add resolution to this section of the tree. The name of *Ca*. Villigracilaceae is proposed for the family. An in-depth analysis of the recently published *Ca*. Villigracilis nielsenii MAG (3) revealed its clustering within the *Ca*. Villigracilaceae family but outside of the *Ca*. Villigracilis genus (**Figure 3**), and we therefore propose to rename it *Candidatus* Manresella nielsenii, reflecting the origin of the sludge (**Additional File 2**).

Similarly, 6 MAGs were classified as g__JADJXA01 (4) and 2 MAGs as g__GCA-2699125 (5), both within the family *Ca*. Promineofilaceae, while MiDAS4 taxonomy only recognized them as two species within the novel genus midas_g_461 (**Table 1**). Based on the GTDB taxonomy, confirmed by additional ANI analysis (**Figure 4**), we have named them *Ca.* Leptovillus (with the novel species *Ca.* Leptovillus affinis (5 MAGs) and gracilis (1 MAG) and *Ca.* Leptofilum (with the two species *Ca.* Leptofilum proximum (1 MAG) and gracile (1 MAG)). Furthermore, one MAG grouped together with *Ca*. Promineofilum breve (6) and represented a new species in the same genus (78% ANI) (**Figure 4**), and 2 MAGs, classified as g__JADJSZ01 and g__JADJUV01, represent novel genera and species, for which we propose the names *Ca*. Hadersleviella danica and *Ca*. Trichofilum aggregatum (**Table 1)**.

Five MAGs represented novel genera and species in the undescribed family f__A4b, within the order Aggregatineales. GTDB-Tk classified 2 MAGs as g__OLB13 and 2 MAGs as g__OLB15, while MiDAS 4 slightly differed by separating the latter into 2 genera (**Table 1**). As our additional ANI analysis confirmed the GTDB taxonomy (**Figure 5**), we proposed the names *Ca*. Flexicrinis (with species *Ca*. Flexicrinis affinis and *Ca.* Flexicrinis proximus) and *Ca*. Flexifilum (with species *Ca*. Flexifilum breve and *Ca*. Flexifilum affine) for the novel genera, and *Ca*. Flexilaceae for the family.

Additionally, four MAGs belonged to the well-known genus *Ca*. Amarolinea and represented a new species. This was confirmed by whole genome ANI (**Figure 6**), which was between the genus and species ANI thresholds of 75% and 95%, respectively (7, 8). Similarly, the GTDB-Tk tool classified one MAG as a novel species within the genus *Caldilinea* and it clustered together with *Caldilinea aerophila* (GCF_000281175). Our ANI analysis (**Figure 7**) supported this classification, even though MiDAS4 classified it as midas_g_105, and we propose the name *Ca.* Caldilinea saccharophila for this species (**Table 1**). An additional MAG clustered within the *Caldilineaceae* family and was classified by GTDB as the undescribed genus g__JADJPH01. As 16S rRNA gene-based phylogeny supported also the classification into a new genus and species, we propose to rename it as *Ca*. Fredericiella danica (**Table 1**).

Two MAGs were classified to family-level in the *Roseiflexaceae* family by GTDB (**Table 1**). One of them represented a novel species in the *Kouleothrix* genus (79% ANI to *K. aurantiaca*, **Figure 8**), while the second represented a closely related new genus. While 16S rRNA gene-based analysis fully supported the taxonomic assignment for the novel *Kouleothrix* species, it is interesting to notice that it would classify the 4 16S rRNA gene sequences extracted from the MAG Ribe_BAT3C.183 into two separate genera (midas_g_2775 and midas_g_4945). Intragenomic variation of rRNA genes has been previously observed in several prokaryotes and it must be taken into consideration when 16S rRNA gene sequences are used for quantification purposes (9, 10). However, as this is only one of the 4 16S rRNA gene sequences and there is only one MAG for this species, difference in the 16S rRNA gene sequence could also be due to Nanopore sequencing error or misbinning. Here, we propose the names *Ca.* Kouleothrix ribensis and *Ca*. Ribeiella danica for these novel lineages.

Finally, 10 MAGs belonged to the undescribed genera g__FeB-14 (9 MAGs) and g__JACPOQ01 (1 MAG), within the recently described *Tepidiformaceae* family of the *Dehalococcoidia* class (**Table 1**). The close relationship with the isolate *Dehalococcoides mccartyi* was also supported by 16S rRNA gene-based analysis and ANI (**Figure 9**), and we therefore propose the names of *Ca.* Amarobacter glycogenicus and *Ca.* Amarobacillus elongatus for the novel species.

Despite lack of representation by a recovered MAG, we used 16S rRNA gene-based phylogeny, FISH probe design and experimental analysis (see below) to describe three more genera abundant in both global and Danish activated sludge samples. Therefore, we propose to rename the genera with placeholder names midas_g_391, midas_g_550, midas_g_9648 as *Ca*. Amarofilum, *Ca*. Pachofilum, and *Ca*. Tricholinea, respectively (**Table 1**). 16S rRNA gene-based analysis using the MiDAS 4 database showed that midas_g_169 corresponded to *Ca*. Defluviifilum, as defined by Speirs et al., (11) and we suggest to adopt this name in future studies.

**Table 1**. Different phylogenetic taxonomies of Chloroflexota members present in AS**.** New species names proposed in this study, MAG identifiers and MiDAS 4-classification with V1-V3 ASVs details are shown. The species are ordered as in the genome tree in **Figure 1**.

| **New proposed species name** | **MAG identifier** | **GTDB taxonomy at genus level** | **MiDAS 4-classification and ASV details (>98% identity)** |
| --- | --- | --- | --- |
| *Ca.* Epilinea brevis | **Hirt_BATAC.427**, EsbW_MAXAC.090 | g__JADJUE01, s__JADJUE01 sp016710785 | Midas_g_119, midas_s_119, ASV1943* |
| *Ca.* Avedoeria danica | **Aved_BATAC.767** | g__JADJCV01, s__JADJCV01 sp016703025 | Midas_g_1676, midas_s_1676, ASV12858**, ASV19407** |
| *Ca.* Brachythrix odensensis | **OdNW_BATAC.48** | g__JADKDT01, s__JADKDT01 sp016714465 | Midas_g_72, Midas_s_72, ASV240**, ASV282**, ASV753** |
| *Ca.* Defluviilinea gracilis | **Kalu_BAT3C.361** | g_UBA12294, s__UBA12294 sp016716235 | UTCFX1, midas_s_9708, ASV34* |
| *Ca.* Defluviilinea proxima | **Skiv_MAXAC.174** | g_UBA12294, s__UBA12294 sp016721115 | midas_g_9708, midas_s_16519, ASV7206* |
| *Ca.* Villigracilis vicinus | **Skiv_MAXAC.043**, EsbW_MAXAC.032, Aved_MAXAC.057_sub | g_OLB14, s__OLB14 sp016721315 | *Ca*. Villigracilis, midas_s_6664, ASV2620* |
| *Ca.* Villigracilis adiacens | **Aved_BAT3C.518** | g_OLB14, s__OLB14 sp016703605 | *Ca*. Villigracilis, midas_s_15163, ASV4573* |
| *Ca.* Villigracilis propinquus | **OdNW_BATAC.378** | g_OLB14, s__OLB14 sp016714565 | *Ca.* Villigracilis, midas_s_9223, ASV997* |
| *Ca.* Villigracilis affinis | **OdNW_MAXAC.037**, Skiv_MAXAC.035, EsbE_MAXAC.031_sub | g_OLB14, s__OLB14 sp016718275 | *Ca*. Villigracilis, midas_s_471, ASV556***, ASV42184*** |
| *Ca.* Villigracilis proximus | **Mari_MAXAC.029**, OdNE_MAXAC.047, Hirt_MAXAC.030 | g_OLB14, s__OLB14 sp016715195 | *Ca*. Villigracilis, midas_s_471, ASV556* |
| *Ca.* Villigracilis saccharophilus | **EsbW_MAXAC.021**, Hade_MAXAC.042 | g_OLB14, s__OLB14 sp016709305 | *Ca*. Villigracilis, midas_s_471, ASV4757* |
| *Ca.* Hadersleviella danica | **Hade_MAXAC.236_sub** | g__JADJSZ01, s__JADJSZ01 sp016711405 | midas_g_2111, midas_s_2642, ASV1441* |
| *Ca.* Trichofilum aggregatum | **Hirt_MAXAC.142** | g__JADJUV01 s__JADJUV01 sp016716885 | midas_g_1951, midas_s_1951, ASV3428***, ASV9417***, ASV42894***, ASV32145*** |
| *Ca.* Promineofilum glycogenico | **Ega_BAT3C.159** | g__Promineofilum, s__Promineofilum sp016707605 | *Ca*. Promineofilum, midas_s_176, ASV640**, ASV2251**, ASV11392**, ASV15239**, ASV24645**, ASV32837** |
| *Ca*. Leptofilum proximum | **Kalu_MAXAC.106v2** | g__GCA-2699125, s__GCA-2699125 sp016710325 | Midas_g_461, midas_s_1196, ASV706**, ASV3461**, ASV3687**, ASV5480**, ASV70841** |
| *Ca.* Leptofilum gracile | **Fred_BAT3C.445** | g__GCA-2699125, s__GCA-2699125 sp016713825 | Midas_g_461, midas_s_1196, ASV9669* |
| *Ca.* Leptovillus gracilis | **Kalu_BATAC.47** | g__JADJXA01, s__JADJXA01 sp016716065 | Midas_g_461, midas_s_13373, ASV3306** |
| *Ca.* Leptovillus affinis | **AalE_BATAC.251**, Aved_BATAC.192, Lyne_BAT3C.469, Fred_BAT3C.529_sub, Damh_MAXAC.008 | g__JADJXA01, s__JADJXA01 sp016705235 | Midas_g_461, midas_s_461, ASV698* |
| *Ca.* Flexicrinis affinis | **Kalu_BAT3C.186** | g__OLB13, s__OLB13 sp016716525 | OLB13, midas_s_19645, ASV7416* |
| *Ca.* Flexicrinis proximus | **Fred_MAXAC.112** | g__OLB13, s__OLB13 sp016712885 | OLB13, midas_s_1091, ASV1734* |
| *Ca.* Flexifilum breve | **Ribe_BATAC.253,** Fred_BATAC.369 | g__OLB15, s__OLB15 sp016717205 | midas_g_4871, midas_s_4871, ASV9809* |
| *Ca.* Flexifilum affine | **Fred_BATAC.421** | g__OLB15, s__OLB15 sp016713325 | midas_g_5525, midas_s_5525, ASV15537* |
| *Ca.* Amarolinea dominans | **Lyne_BATAC.272,** OdNW_BAT3C.33_sub, Damh_MAXAC.106, AalW_BAT3C.371_sub | g__JADJYJ01, s__JADJYJ01 sp016719785 | *Ca.* Amarolinea, midas_s_1, ASV692* |
| *Ca.* Fredericiella danica | **Fred_BATAC.359** | g__JADJPH01, s__JADJPH01 sp016713335 | midas_g_2265, midas_s_4097, ASV16104* |
| *Ca.* Caldilinea saccharophila | **Hjor_MAXAC.079_sub** | g__Caldilinea, s__Caldilinea sp016710365 | Midas_g_105, midas_s_105, ASV2647***, ASV17928*** |
| *Ca.* Ribeiella danica | **Ribe_BAT3C.183** | g__JADKFS01, s__JADKFS01 sp016717335 | midas_g_2775, midas_s_2775, ASV4502* |
| *Ca.* Kouleothrix ribensis | **Ribe_MAXAC.079_sub** | g__Kouleothrix, s__Kouleothrix sp016722075 | *Kouleothrix*, midas_s_2308, ASV1266* |
| *Ca.* Amarobacter glycogenicus | **Lyne_MAXAC.019**, Ejby_BAT3C.216, OdNW_MAXAC.019, Rand_BAT3C.210, EsbE_MAXAC.015, Damh_BAT3C.259, AalE_BAT3C.409, Hirt_BATAC.715, AalW_MAXAC.018 | g__FeB-14, s__FeB-14 sp016719395 | Midas_g_731, midas_s_849, ASV795** |
| *Ca.* Amarobacillus elongatus | **Aved_BAT3C.689** | g__JACPOQ01, s__JACPOQ01 sp016703545 | Midas_g_1412, midas_s_8871, ASV487**, ASV949**, ASV5646**, ASV9583** |
| *Ca.* Amarofilum | **-** | - | Midas_g_391 |
| *Ca.* Pachofilum | **-** | - | Midas_g_550 |
| *Ca.* Tricholinea | **-** | - | Midas_g_9648 |
| *Ca.* Defluviifilum | **-** | - | Midas_g_169 |

* 100% identity hit

** > 99% identity hit

*** > 98% identity hit

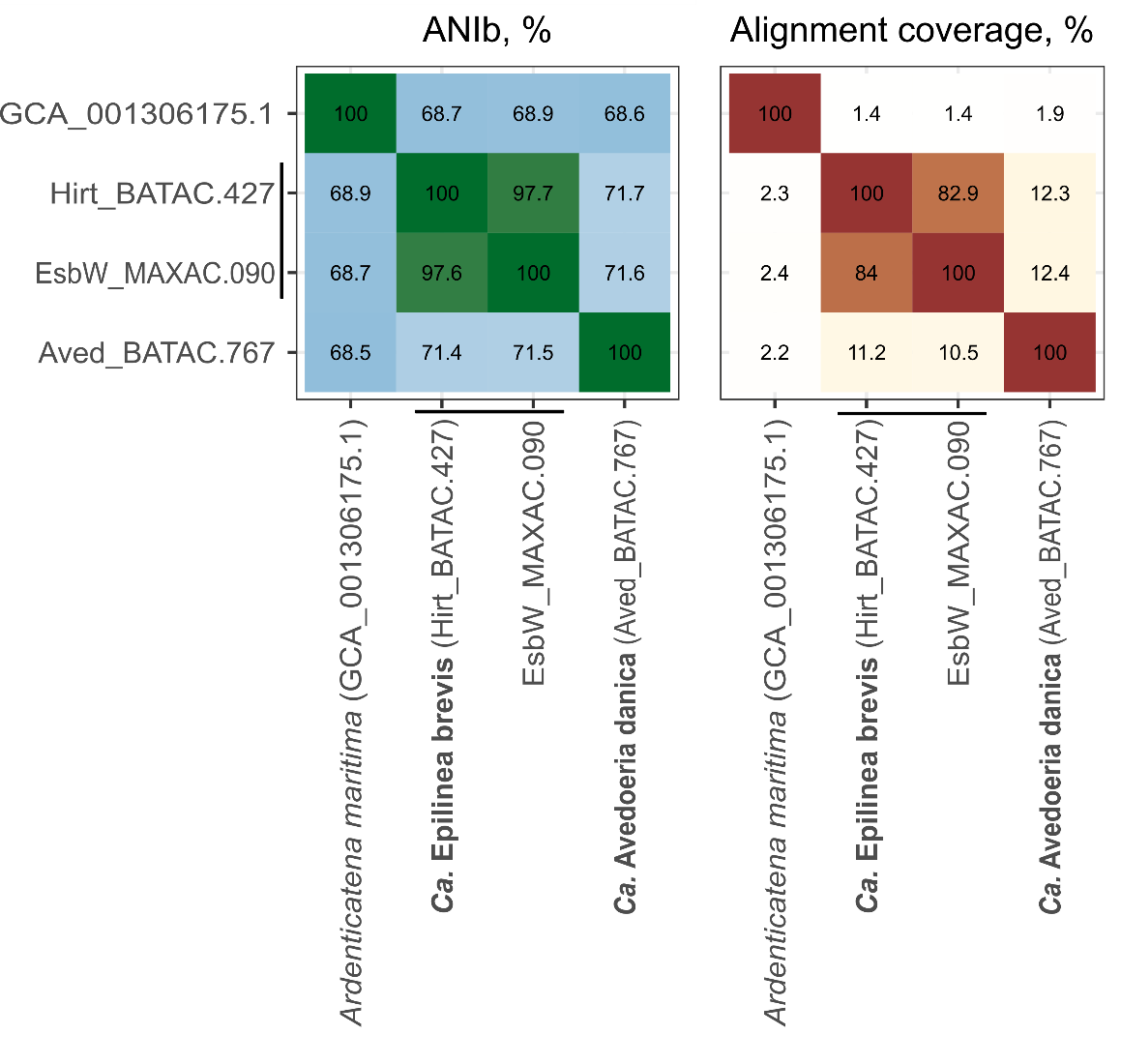

**Figure 1.** Average nucleotide identity (%) using BLASTN+ (ANIb) and alignment coverage (%) of the genomes and MAGs belonging to the *Epilineaceae* family. New species names are written in bold next to representative MAG.

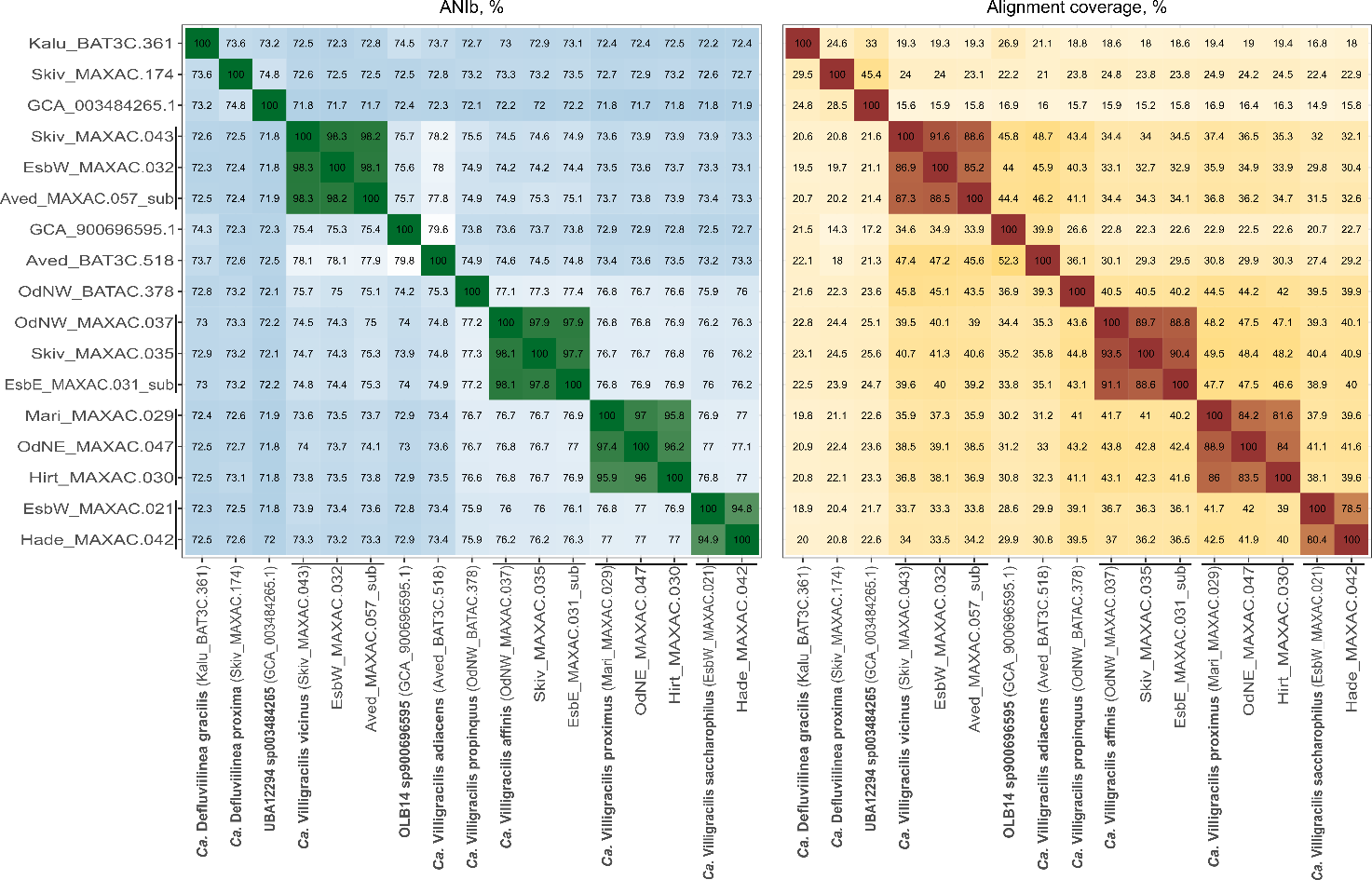

**Figure 2.** Average nucleotide identity (%) using BLASTN+ (ANIb) and alignment coverage (%) of the genomes and MAGs belonging to the *Ca.* Villigracilaceae family. New species names are written in bold next to representative MAG.

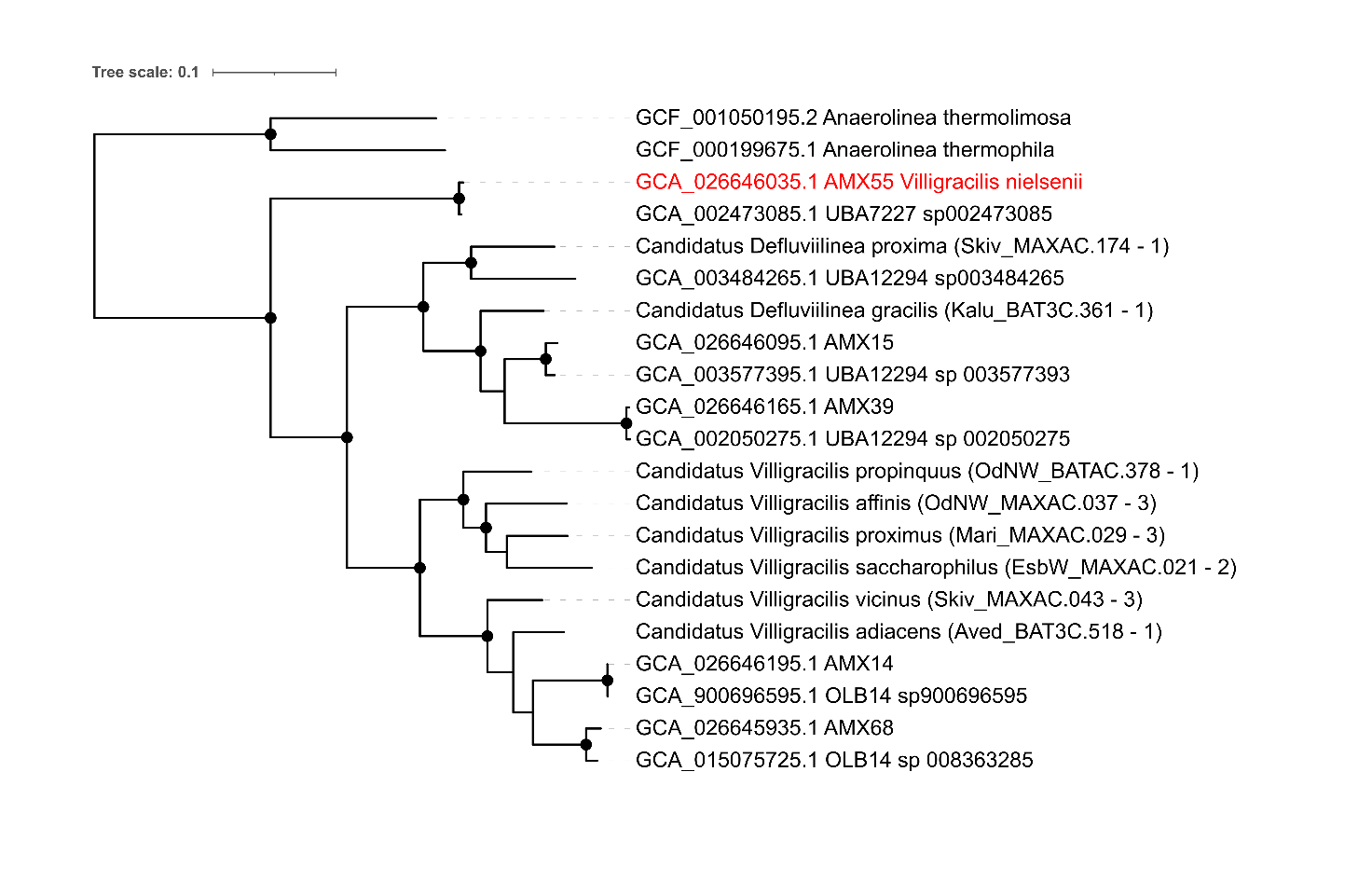

**Figure 3**. **Phylogenetic genome tree of *Ca.* Villigracilaceae representatives.** The Chloroflexota MAGs from Bovio-Winker et al. 2023 (3) most closely related to the *Ca.* Villigracilis MAGs in this study were downloaded from NCBI. The MAGs from both studies were combined and used as input for the GTDB-Tk de_novo_wf. The maxinum likelihood tree was created from the concatenated alignment of 120 single copy marker genes created by GTDB-Tk trimmed to ~5000 amino acids using the WAG+G model and 1000x UFBoot bootstrap iterations in IQ-TREE. Bootstrap support >95% is indicated by the black circles. The red MAG indicates *Ca*. Villigracilis nielsenii.

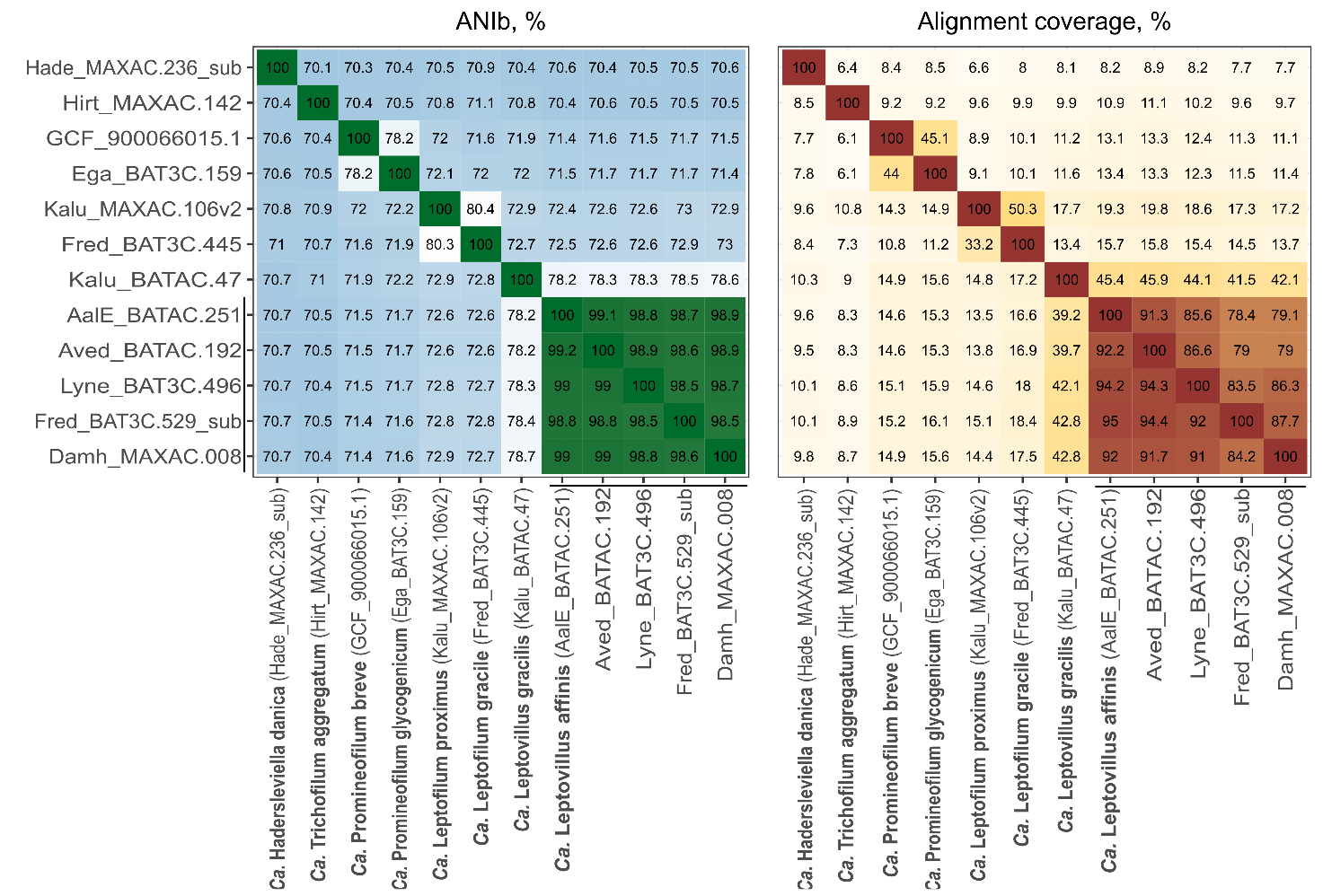

**Figure 4.** Average nucleotide identity (%) using BLASTN+ (ANIb) and alignment coverage (%) of the genomes and MAGs belonging to the *Ca*. Promineofilaceae family. New species names are written in bold next to representative MAG.

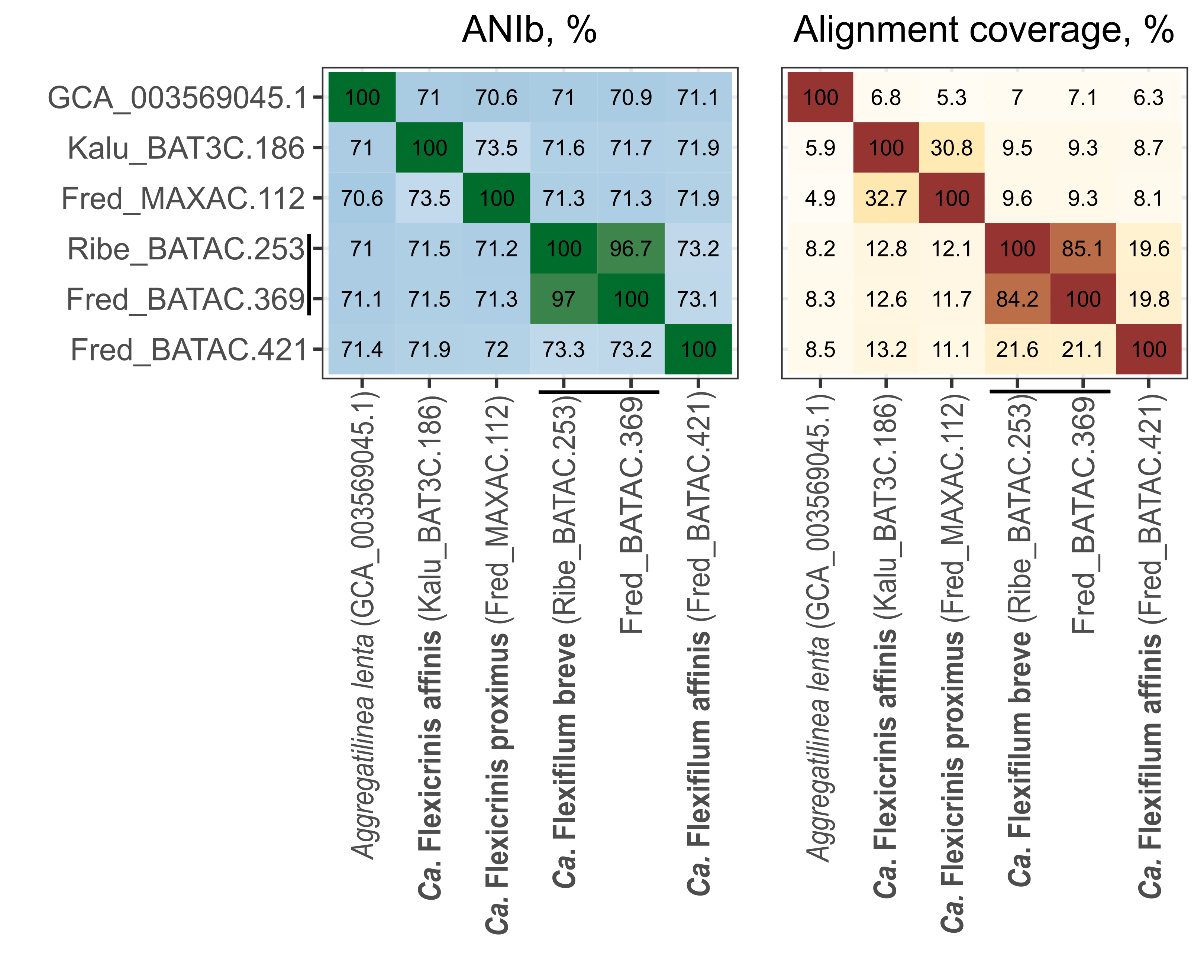

**Figure 5.** Average nucleotide identity (%) using BLASTN+ (ANIb) and alignment coverage (%) of the genomes and MAGs belonging to the *Flexifilaceae* family. New species names are written in bold next to representative MAG.

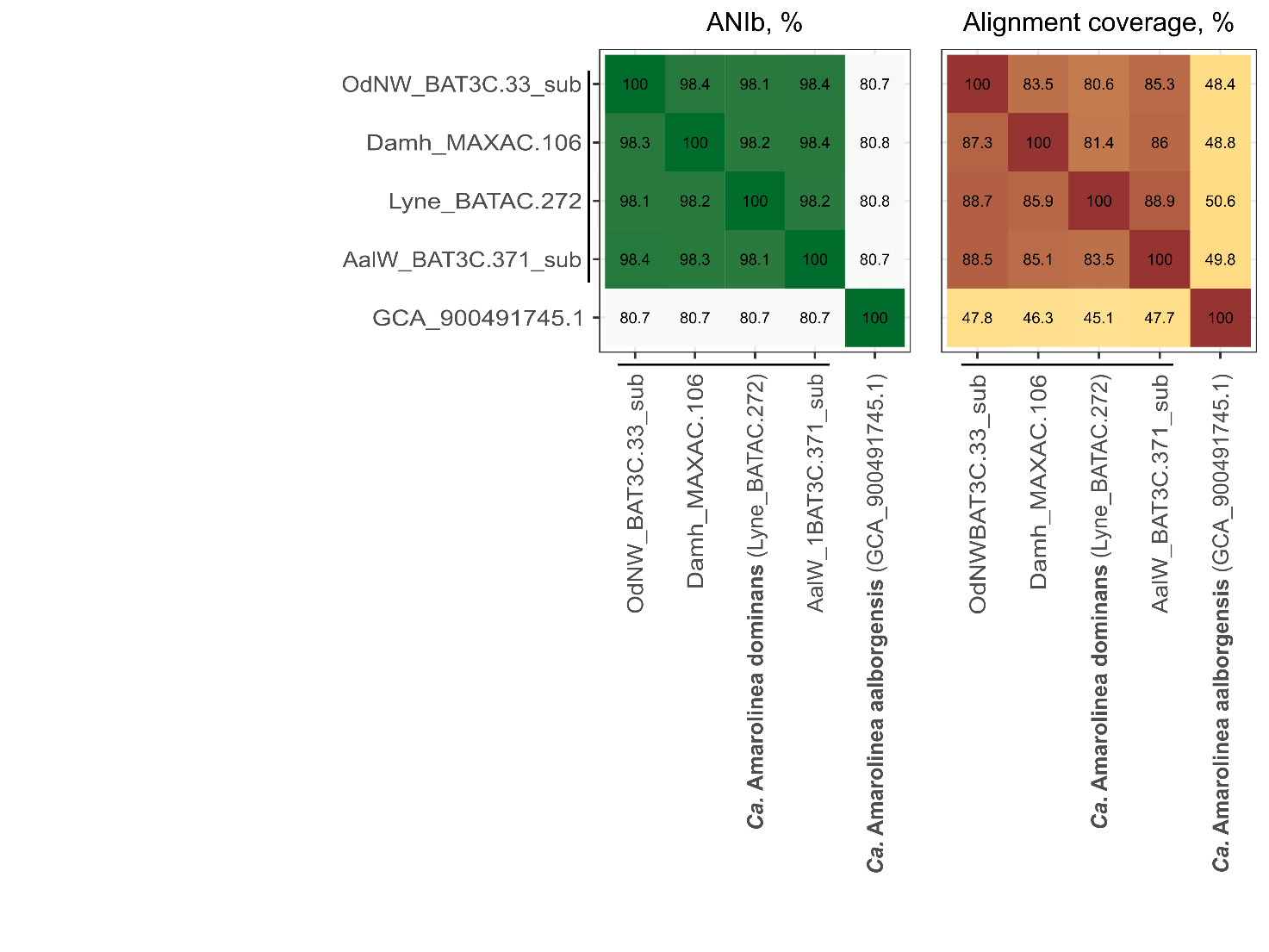

**Figure 6.** Average nucleotide identity (%) using BLASTN+ (ANIb) and alignment coverage (%) of the genomes and MAGs belonging to the genus *Ca*. Amarolinea. New species names are written in bold next to representative MAG.

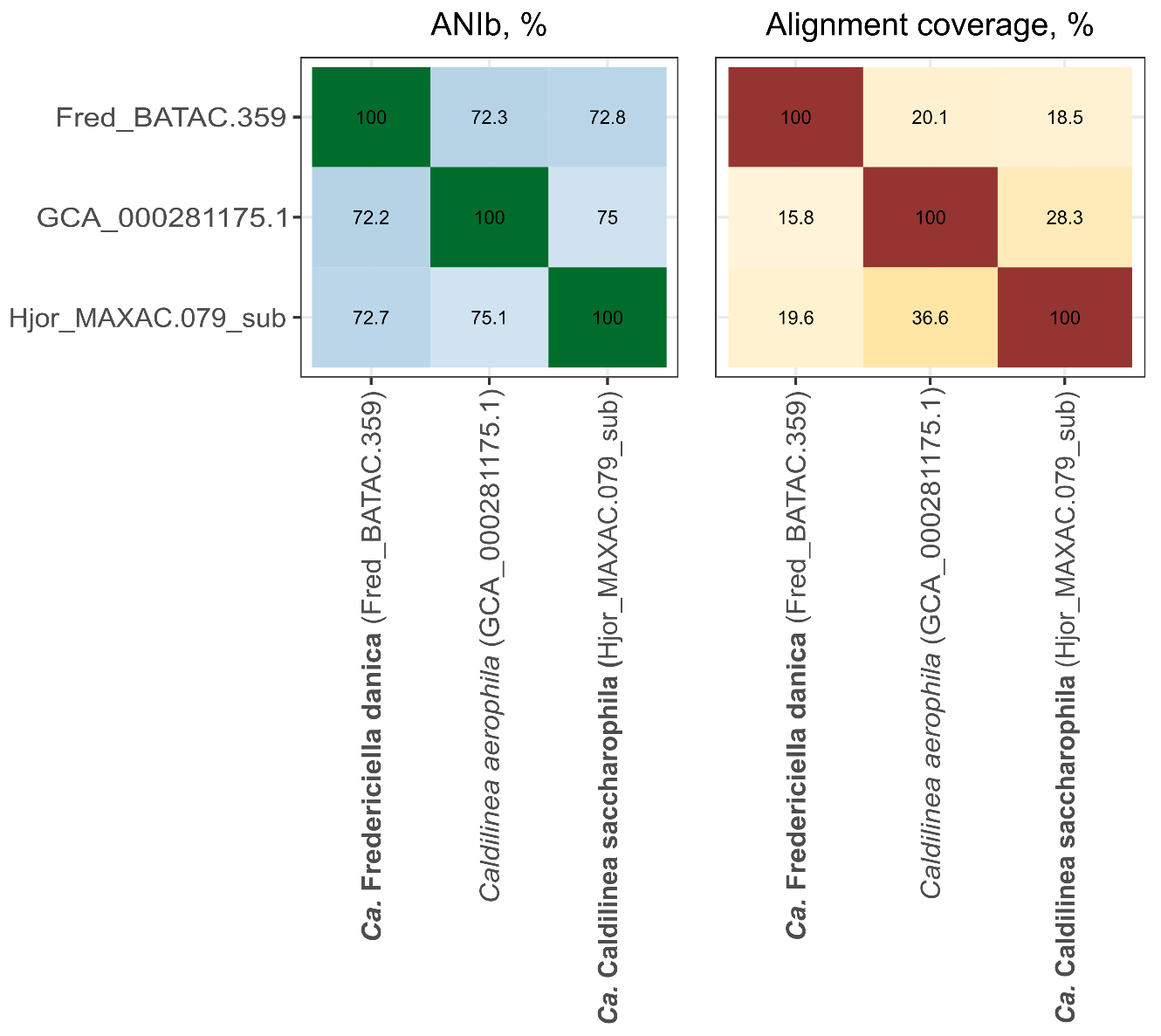

**Figure 7**. Average nucleotide identity (%) using BLASTN+ (ANIb) and alignment coverage (%) of *Caldilinea aerophila* and the MAG Hjor_MAXAC.079_sub. New species names are written in bold next to representative MAG.

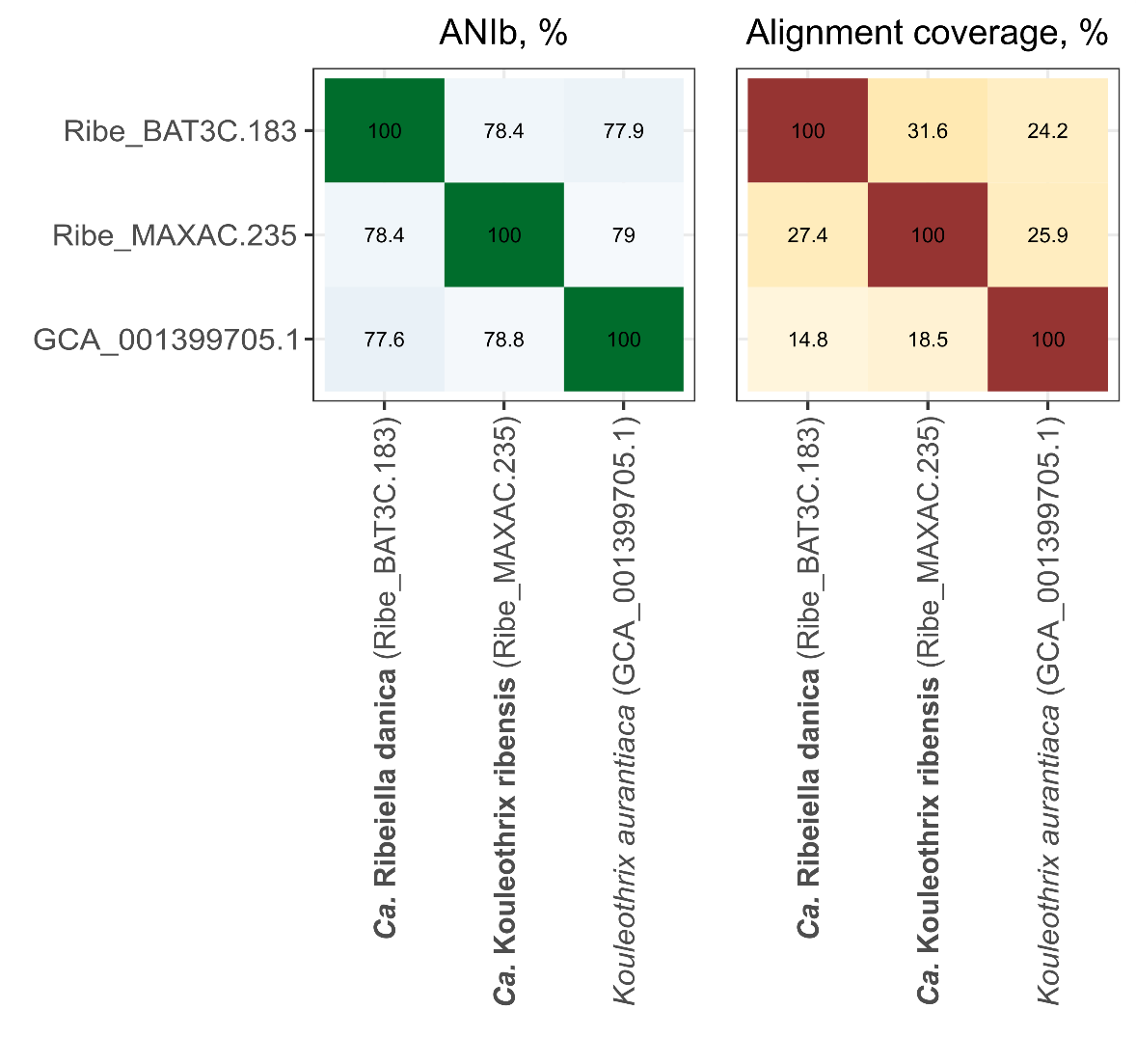

**Figure 8.** Average nucleotide identity (%) using BLASTN+ (ANIb) and alignment coverage (%) of the genomes and MAGs belonging to the genus *Kouleothrix*. New species names are written in bold next to representative MAG.

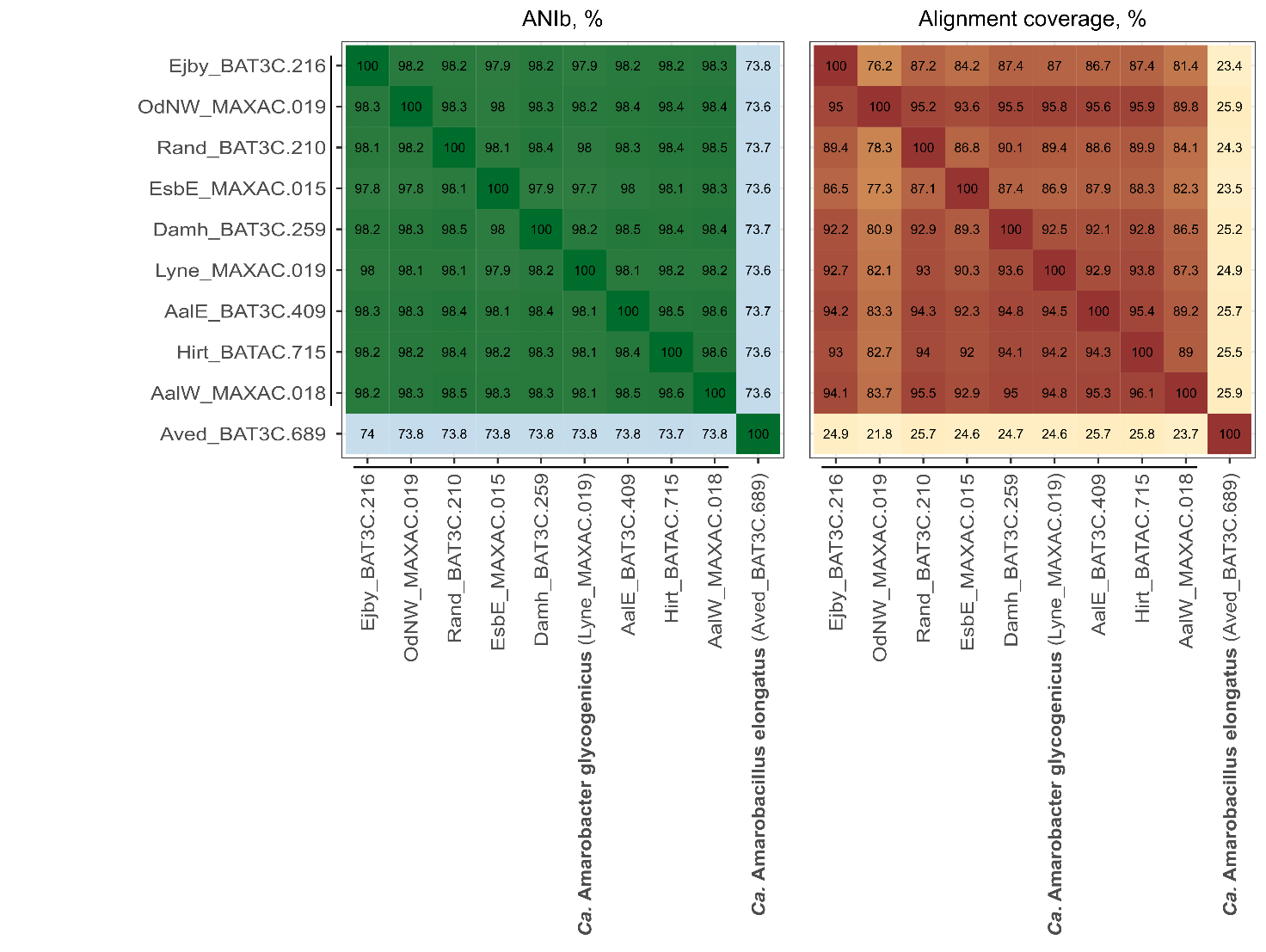

**Figure 9.** Average nucleotide identity (%) using BLASTN+ (ANIb) and alignment coverage (%) of the genomes and MAGs belonging to the *Ca*. Amarobacterceae family. New species names are written in bold next to representative MAG.

**References**

1.	Lawson CE, Wu S, Bhattacharjee AS, Hamilton JJ, Mcmahon KD, Goel R, Noguera DR. 2017. Metabolic network analysis reveals microbial community interactions in anammox granules. Nat Commun 8:1–12.

2.	Speth DR, In’T Zandt MH, Guerrero-Cruz S, Dutilh BE, Jetten MSM. 2016. Genome-based microbial ecology of anammox granules in a full-scale wastewater treatment system. Nat Commun 7:11172.

3.	Bovio-Winkler P, Guerrero LD, Erijman L, Oyarzúa P, Suárez-Ojeda ME, Cabezas A, Etchebehere C. 2023. Genome-centric metagenomic insights into the role of Chloroflexi in anammox, activated sludge and methanogenic reactors. BMC Microbiol 23:45.

4.	Bik EM, Costello EK, Switzer AD, Callahan BJ, Holmes SP, Wells RS, Carlin KP, Jensen ED, Venn-Watson S, Relman DA. 2016. Marine mammals harbor unique microbiotas shaped by and yet distinct from the sea. Nat Commun 7:1–13.

5.	Tully BJ, Graham ED, Heidelberg JF. 2018. The reconstruction of 2,631 draft metagenome-assembled genomes from the global oceans. Sci Data 5:1–8.

6.	McIlroy SJ, Karst SM, Nierychlo M, Dueholm MS, Albertsen M, Kirkegaard RH, Seviour RJ, Nielsen PH. 2016. Genomic and *in situ* investigations of the novel uncultured Chloroflexi associated with 0092 morphotype filamentous bulking in activated sludge. ISME J 10: 2223–2234.

7.	Parks DH, Chuvochina M, Chaumeil P-A, Rinke C, Mussig AJ, Hugenholtz P. 2020. A complete domain-to-species taxonomy for Bacteria and Archaea. Nat Biotechnol 38:1079–1086.

8.	Barco RA, Garrity GM, Scott JJ, Amend JP, Nealson KH, Emerson D, J. GS. 2020. A genus definition for Bacteria and Archaea based on a standard genome relatedness index. MBio 11:e02475-19.

9.	Wagner J, Coupland P, Browne HP, Lawley TD, Francis SC, Parkhill J. 2016. Evaluation of PacBio sequencing for full-length bacterial 16S rRNA gene classification. BMC Microbiol 16:1–17.

10.	Sun DL, Jiang X, Wu QL, Zhou NY. 2013. Intragenomic heterogeneity of 16S rRNA genes causes overestimation of prokaryotic diversity. Appl Environ Microbiol 79:5962–5969.

11.	Speirs LBM, Rice DTF, Petrovski S, Seviour RJ, Mcilroy SJ. 2019. The phylogeny, biodiversity, and ecology of the Chloroflexi in activated sludge. Front Microbiol 10:10:2015.
