## Additional File 2 - Taxonomic proposal and protologue tables for "A comprehensive overview of the Chloroflexota community in wastewater treatment plants worldwide"

**Additional File 2 – Taxonomic proposal and protologue tables for *Candidatus* species**

Description of ‘*Candidatus* Epilinea brevis’ gen. nov. sp. nov.: “*Candidatus* Epilinea brevis”, (E.pi.li´ne.a, G. prep. *epi*, on; L. fem. n. *linea*, line, filament; N.L. fem. n. Epilinea, indicating filamentous bacteria attached to other filaments; bre´vis. L. fem. adj. *brevis*, short, indicating the short length of the filaments). This taxon is represented by the MAG Hirt_BATAC.427. The complete protologue can be found in Table 1.

Description of Epilineaceae fam. nov.: *Epilineaceae* (E.pi.li.ne.a.ce´ae. from N.L. fem. n. Epilinea type genus of the family; suff. -aceae ending to denote a family; N.L. fem. pl. n. Epilineaceae, the Epilinea family).

Description of Epilineales ord. nov.: Epilineales (E.pi.li.ne.a´les. from N.L. fem. n. Epilinea type genus of the family; suff. -ales ending to denote an order; N.L. fem. pl. n. Epilineales, the Epilinea order).

**Table 1. Protologue Table for *Candidatus* Epilinea brevis**

| Species name | *Candidatus* Epilinea brevis |
| --- | --- |
| Genus name | *Candidatus* Epilinea |
| Specific epithet | brevis |
| Type species of the genus | *Candidatus* Epilinea brevis |
| Genus status | Candidatus |
| Species etymology | “*Candidatus* Epilinea brevis”, (E.pi.li´ne.a, G. prep. epi, on; L. fem. n. linea, line, filament; N.L. fem. n. Epilinea, indicating filamentous bacteria attached to other filaments; bre´vis. L. fem. adj. brevis, short, indicating the short length of the filaments. |
| Species status | sp. nov. |
| Designation of the type MAG | GCA_016710785.1 |
| MAG/SAG accession number | GCA_016710785.1 |
| Genome status | High-quality draft |
| Genome size | 4924095 |
| GC mol % | 69.08 |
| Country of origin | Denmark |
| Region of origin | Hirtshals |
| Source of sample | Activated sludge |
| Geographical location | Hirtshals |
| Latitude | 57.577275 |
| Longitude | 9.992971 |
| Depth | N/A |
| Altitude | N/A |
| Temperature of the sample | Mesophilic |
| pH of the sample | N/A |
| Relationship to oxygen | Facultative anaerobe |
| Energy metabolism | Potentially utilizing carbohydrates, fatty acids and amino acids |
| Assembly | 1 sample |
| Sequencing technology | Oxford Nanopore PromethION |
| Binning software used | MetaBAT2 |
| Assembly software used | CANU v.1.8 |
| Habitat | Full-scale enriched biological phosphorus removal wastewater treatment plant |
| Miscellaneous, extraordinary features relevant for the description | Filamentous morphology (4-57 × 0.4-0.7 µm) |

Description of “*Candidatus* Avedoeria danica” gen. nov. sp. nov.: “*Candidatus* Avedoeria danica”, (A.ve.do.e´ri.a., N.L. fem. n. *Avedoeria,* named after the city Avedoere where the MAG had been retrieved; da´ni.ca. M.L. fem. adj. *danica*, danish; indicating the country of origin). This taxon is represented by the MAG Aved_BATAC.767. The complete protologue can be found in Table 2.

**Table 2. Protologue Table for *Candidatus* Avedoeria danica**

| Species name | *Candidatus* Avedoeria danica |
| --- | --- |
| Genus name | *Candidatus* Avedoeria |
| Specific epithet | danica |
| Type species of the genus | *Candidatus* Avedoeria danica |
| Genus status | Candidatus |
| Species etymology | “*Candidatus* Avedoeria danica”, (A.ve.do.e´ri.a., N.L. fem. n. *Avedoeria,* named after the city Avedoere where the MAG had been retrieved; da´ni.ca. M.L. fem. adj. *danica*, danish; indicating the country of origin). |
| Species status | sp. nov. |
| Designation of the type MAG | GCA_016703025.1 |
| MAG/SAG accession number | GCA_016703025.1 |
| Genome status | High-quality draft |
| Genome size | 4347427 |
| GC mol % | 70.87 |
| Country of origin | Denmark |
| Region of origin | Avedøre |
| Source of sample | Activated sludge |
| Geographical location | Avedøre |
| Latitude | 55.608613 |
| Longitude | 12.450537 |
| Depth | N/A |
| Altitude | N/A |
| Temperature of the sample | Mesophilic |
| pH of the sample | N/A |
| Relationship to oxygen |  |
| Energy metabolism |  |
| Assembly | 1 sample |
| Sequencing technology | Oxford Nanopore PromethION |
| Binning software used | MetaBAT2 |
| Assembly software used | CANU v.1.8 |
| Habitat | Full-scale enriched biological phosphorus removal wastewater treatment plant |
| Miscellaneous, extraordinary features relevant for the description |  |

Description of ‘*Candidatus* Brachythrix odensensis’ gen. nov. sp. nov.: “*Candidatus* Brachythrix”, (Bra´chy.thrix, G. fem. adj. *brachys*, short; G. fem. n. *thrix*, hair, filament; N.L. fem. n. Brachythrix, short filamentous bacterium; o.den.sen’sis. (N.L. fem. adj. *odensensis*, pertinent to the city of Odense, from where the MAG was obtained). This taxon is represented by the MAG OdNW_BATAC.48. The complete protologue can be found in Table 3.

Description of Brachytrichaceae fam. nov.: *Brachytrichaceae* (Bra.chy.tri.cha.ce´ae, from N.L. fem. n. Brachythrix type genus of the family; suff. -aceae ending to denote a family; N.L. fem. pl. n. Brachytrichaceae, the Brachythrix family).

**Table 3. Protologue Table for *Candidatus* Brachythrix odensensis**

| Species name | *Candidatus* Brachythrix odensensis |
| --- | --- |
| Genus name | *Candidatus* Brachythrix |
| Specific epithet | odensensis |
| Type species of the genus | *Candidatus* Brachythrix odensensis |
| Genus status | Candidatus |
| Species etymology | “*Candidatus* Brachythrix”, (Bra´chy.thrix, G. fem. adj. *brachys*, short; G. fem. n. *thrix*, hair, filament; N.L. fem. n. *Brachythrix*, short filamentous bacterium; o.den.sen’sis. (N.L. fem. adj. *odensensis*, pertinent to the city of Odense, from where the MAG was obtained). |
| Species status | Candidatus |
| Designation of the type MAG | GCA_016714465.1 |
| MAG/SAG accession number | GCA_016714465.1 |
| Genome status | High-quality draft |
| Genome size | 4628329 |
| GC mol % | 64.67 |
| Country of origin | Denmark |
| Region of origin | Odense |
| Source of sample | Activated sludge |
| Geographical location | Odense |
| Latitude | 55.421534 |
| Longitude | 10.366234 |
| Depth | N/A |
| Altitude | N/A |
| Temperature of the sample | Mesophilic |
| pH of the sample | N/A |
| Relationship to oxygen | Facultative anaerobe |
| Energy metabolism | Potentially utilizing carbohydrates, amino acids, and fatty acids |
| Assembly | 1 sample |
| Sequencing technology | Oxford Nanopore PromethION |
| Binning software used | MetaBAT2 |
| Assembly software used | CANU v.1.8 |
| Habitat | Full-scale enriched biological phosphorus removal wastewater treatment plant |
| Miscellaneous, extraordinary features relevant for the description | Filamentous morphology (2.5-18 × 0.5-0.9 µm) |

Description of ‘*Candidatus* Defluviilinea’ gen. nov.: “*Candidatus* Defluviilinea”, (De.flu.vi.i.li´ne.a. L. neut. n. *defluvium*, sewage; L. fem. n. *linea*, line, filament; N.L. fem. n. *Defluviilinea*, filamentous bacteria found in sewage).

Description of ‘*Candidatus* Defluviilinea gracilis’ sp. nov.: “*Candidatus* Defluviilinea gracilis”, (gra´ci.lis. L. fem. adj. *gracilis*, thin, indicating the thin trichome). This taxon is represented by the MAG Kalu_BAT3C.361. The complete protologue can be found in Table 4.

Description of ‘*Candidatus* Defluviilinea proxima’ sp. nov.: “*Candidatus* Defluviilinea proxima”, (pro´xi.ma. L. fem. adj. *proxima*, next of kin, indicating the close phylogenetic relationship with *Ca.* Defluviilinea gracilis). This taxon is represented by the MAG Skiv_MAXAC.174. The complete protologue can be found in Table 5.

**Table 4. Protologue Table for *Candidatus* Defluviilinea gracilis**

| Species name | *Candidatus* Defluviilinea gracilis |
| --- | --- |
| Genus name | *Candidatus* Defluviilinea |
| Specific epithet | gracilis |
| Type species of the genus | *Candidatus* Defluviilinea gracilis |
| Genus status | Candidatus |
| Species etymology | “*Candidatus* Defluviilinea gracilis”, (De.flu.vi.i.li´ne.a. L. neut. n. *defluvium*, sewage; L. fem. n. *linea*, line, filament; N.L. fem. n. *Defluviilinea*, filamentous bacteria found in sewage; gra´ci.lis. L. fem. adj. *gracilis*, thin, indicating the thin trichome). |
| Species status | Candidatus |
| Designation of the type MAG | GCA_016716235.1 |
| MAG/SAG accession number | GCA_016716235.1 |
| Genome status | High-quality draft |
| Genome size | 4139007 |
| GC mol % | 53.56 |
| Country of origin | Denmark |
| Region of origin | Kalundborg |
| Source of sample | Activated sludge |
| Geographical location | Kalundborg |
| Latitude | 55.668006 |
| Longitude | 11.107247 |
| Depth | N/A |
| Altitude | N/A |
| Temperature of the sample | Mesophilic |
| pH of the sample | N/A |
| Relationship to oxygen | Facultative anaerobe |
| Energy metabolism | Potentially utilizing carbohydrates, amino acids, and fatty acids |
| Assembly | 1 sample |
| Sequencing technology | Oxford Nanopore PromethION |
| Binning software used | MetaBAT2 |
| Assembly software used | CANU v.1.8 |
| Habitat | Full-scale enriched biological phosphorus removal wastewater treatment plant |
| Miscellaneous, extraordinary features relevant for the description | Filamentous morphology (12-50 × 0.3-0.4 µm) |

**Table 5. Protologue Table for *Candidatus* Defluviilinea proxima**

| Species name | *Candidatus* Defluviilinea proxima |
| --- | --- |
| Genus name | *Candidatus* Defluviilinea |
| Specific epithet | proxima |
| Type species of the genus | *Candidatus* Defluviilinea gracilis |
| Genus status | Candidatus |
| Species etymology | “*Candidatus* Defluviilinea gracilis”, (De.flu.vi.i.li´ne.a. L. neut. n. *defluvium*, sewage; L. fem. n. *linea*, line, filament; N.L. fem. n. *Defluviilinea*, filamentous bacteria found in sewage; (pro´xi.ma. L. fem. adj. proxima, next of kin, indicating the close phylogenetic relationship with *Ca*. Defluviilinea gracilis) |
| Species status | Candidatus |
| Designation of the type MAG | GCA_016716235.1 |
| MAG/SAG accession number | GCA_016721115.1 |
| Genome status | High-quality draft |
| Genome size | 5022719 |
| GC mol % | 48.58 |
| Country of origin | Denmark |
| Region of origin | Skive |
| Source of sample | Activated sludge |
| Geographical location | Skive |
| Latitude | 56.565132 |
| Longitude | 9.042158 |
| Depth | N/A |
| Altitude | N/A |
| Temperature of the sample | Mesophilic |
| pH of the sample | N/A |
| Relationship to oxygen | Facultative anaerobe |
| Energy metabolism | Potentially utilizing carbohydrates, amino acids, and fatty acids |
| Assembly | 1 sample |
| Sequencing technology | Oxford Nanopore PromethION |
| Binning software used | MaxBin2 |
| Assembly software used | CANU v.1.8 |
| Habitat | Full-scale enriched biological phosphorus removal wastewater treatment plant |
| Miscellaneous, extraordinary features relevant for the description | Filamentous morphology (12-50 × 0.3-0.4 µm) |

Description of “*Candidatus* Villigracilis vicinus” sp. nov.: “*Candidatus* Villigracilis vicinus” (vi.ci´nus. L. masc. adj. *vicinus*, close; indicating the close phylogenetic relationship with *Ca*. Villigracilis saccharophilus). This taxon is represented by the MAG Skiv_MAXAC.043. The complete protologue can be found in Table 6.

Description of “*Candidatus* Villigracilis adiacens” sp. nov.: “*Candidatus* Villigracilis adiacens” (ad’ia.cens, L. part. adj. *adiacens*, close; indicating the close phylogenetic relationship with *Ca*. Villigracilis saccharophilus). This taxon is represented by the MAG Aved_BAT3C.518. The complete protologue can be found in Table 7.

Description of “*Candidatus* Villigracilis propinquus” sp. nov.: “*Candidatus* Villigracilis propinquus,” (pro.pin´qu.us. L. masc. adj. *propinquus*, next of kin; indicating the close phylogenetic relationship with *Ca*. Villigracilis saccharophilus). This taxon is represented by the MAG OdNW_BATAC.378. The complete protologue can be found in Table 8.

Description of “*Candidatus* Villigracilis affinis” sp. nov.: “*Candidatus* Villigracilis affinis,” (af.fi´nis. L. masc. adj. *affinis*, next of kin; indicating the close phylogenetic relationship with *Ca*. Villigracilis saccharophilus). This taxon is represented by the MAG OdNW_MAXAC.037. The complete protologue can be found in Table 9.

Description of “*Candidatus* Villigracilis proximus” sp. nov.: “*Candidatus* Villigracilis proximus,” (pro´xi.mus. L. masc. adj. *proximus*, next of kin; indicating the close phylogenetic with *Ca*. Villigracilis saccharophilus). This taxon is represented by the MAG OdNE_MAXAC.047. The complete protologue can be found in Table 10.

Description of ‘*Candidatus* Villigracilis saccharophilus’ sp. nov.: “*Candidatus* Villigracilis saccharophilus”, (sac.cha.ro´phi.lus. G. neut. n. *saccharon*, sugar; G. masc. n. *philos*, lover; N.L. masc. adj. saccharophilus, indicating a preference for sugars as carbon sources). This taxon is represented by the MAG EsbW_MAXAC.021. The complete protologue can be found in Table 11.

Description of Villigracilaceae fam. nov.: *Villigracilaceae* (Vil.li.gra.ci.la.ce´ae, from N.L. masc. n. Villigracilis type genus of the family; suff. -aceae ending to denote a family; N.L. fem. pl. n. Villigracilaceae, the Villigracilis family).

**Table 6. Protologue Table for *Candidatus* Villigracilis vicinus**

| Species name | *Candidatus* Villigracilis vicinus |
| --- | --- |
| Genus name | *Candidatus* Villigracilis |
| Specific epithet | vicinus |
| Type species of the genus | *Candidatus* Villigracilis vicinus |
| Genus status | Candidatus |
| Species etymology | Description of “*Candidatus* Villigracilis vicinus” sp. nov. “*Candidatus* Villigracilis vicinus” (vi.ci´nus. L. masc. adj. *vicinus*, close; indicating the close phylogenetic relationship with *Ca*. Villigracilis saccharophilus). |
| Species status | Candidatus |
| Designation of the type MAG | GCA_016709305.1 |
| MAG/SAG accession number | GCA_016721315.1 |
| Genome status | High-quality draft |
| Genome size | 4491201 |
| GC mol % | 50.26 |
| Country of origin | Denmark |
| Region of origin | Skive |
| Source of sample | Activated sludge |
| Geographical location | Skive |
| Latitude | 56.565132 |
| Longitude | 9.042158 |
| Depth | N/A |
| Altitude | N/A |
| Temperature of the sample | Mesophilic |
| pH of the sample | N/A |
| Relationship to oxygen | Facultative anaerobe |
| Energy metabolism | Potentially utilizing carbohydrates, amino acids, and fatty acids |
| Assembly | 1 sample |
| Sequencing technology | Oxford Nanopore PromethION |
| Binning software used | MaxBin2 |
| Assembly software used | CANU v.1.8 |
| Habitat | Full-scale enriched biological phosphorus removal wastewater treatment plant |
| Miscellaneous, extraordinary features relevant for the description | Filamentous morphology (12-50 × 0.3-0.4 µm) |

**Table 7. Protologue Table for *Candidatus* Villigracilis adiacens**

| Species name | *Candidatus* Villigracilis adiacens |
| --- | --- |
| Genus name | *Candidatus* Villigracilis |
| Specific epithet | adiacens |
| Type species of the genus | *Candidatus* Defluviilinea gracilis |
| Genus status | Candidatus |
| Species etymology | Description of “*Candidatus* Villigracilis adiacens” sp. nov. “*Candidatus* Villigracilis adiacens” (ad’ia.cens, L. part. adj. *adiacens*, close; indicating the close phylogenetic relationship with *Ca*. Villigracilis saccharophilus). |
| Species status | Candidatus |
| Designation of the type MAG | GCA_016709305.1 |
| MAG/SAG accession number | GCA_016703605.1 |
| Genome status | High-quality draft |
| Genome size | 4437588 |
| GC mol % | 54.97 |
| Country of origin | Denmark |
| Region of origin | Avedøre |
| Source of sample | Activated sludge |
| Geographical location | Avedøre |
| Latitude | 55.608613 |
| Longitude | 12.450537 |
| Depth | N/A |
| Altitude | N/A |
| Temperature of the sample | Mesophilic |
| pH of the sample | N/A |
| Relationship to oxygen | Facultative anaerobe |
| Energy metabolism | Potentially utilizing carbohydrates, amino acids, and fatty acids |
| Assembly | 1 sample |
| Sequencing technology | Oxford Nanopore PromethION |
| Binning software used | MetaBAT2 |
| Assembly software used | CANU v.1.8 |
| Habitat | Full-scale enriched biological phosphorus removal wastewater treatment plant |
| Miscellaneous, extraordinary features relevant for the description | Filamentous morphology (12-50 × 0.3-0.4 µm) |

**Table 8. Protologue Table for *Candidatus* Villigracilis propinquus**

| Species name | *Candidatus* Villigracilis propinquus |
| --- | --- |
| Genus name | *Candidatus* Villigracilis |
| Specific epithet | propinquus |
| Type species of the genus | *Candidatus* Villigracilis affinis |
| Genus status | Candidatus |
| Species etymology | “*Candidatus* Villigracilis propinquus”, (pro.pin´qu.us. L. masc. adj. propinquus, next of kin; indicating the close phylogenetic relationship with Ca. Villigracilis saccharophilus). |
| Species status | Candidatus |
| Designation of the type MAG | GCA_016709305.1 |
| MAG/SAG accession number | GCA_016714565.1 |
| Genome status | High-quality draft |
| Genome size | 4941904 |
| GC mol % | 50.88 |
| Country of origin | Denmark |
| Region of origin | Odense |
| Source of sample | Activated sludge |
| Geographical location | Odense |
| Latitude | 55.421534 |
| Longitude | 10.366234 |
| Depth | N/A |
| Altitude | N/A |
| Temperature of the sample | Mesophilic |
| pH of the sample | N/A |
| Relationship to oxygen | Facultative anaerobe |
| Energy metabolism | Potentially utilizing carbohydrates, amino acids, and fatty acids |
| Assembly | 1 sample |
| Sequencing technology | Oxford Nanopore PromethION |
| Binning software used | MetaBAT2 |
| Assembly software used | CANU v.1.8 |
| Habitat | Full-scale enriched biological phosphorus removal wastewater treatment plant |
| Miscellaneous, extraordinary features relevant for the description | Filamentous morphology (12-50 × 0.3-0.4 µm) |

**Table 9. Protologue Table for *Candidatus* Villigracilis affinis**

| Species name | *Candidatus* Villigracilis affinis |
| --- | --- |
| Genus name | *Candidatus* Villigracilis |
| Specific epithet | affinis |
| Type species of the genus | *Candidatus* Villigracilis affinis |
| Genus status | Candidatus |
| Species etymology | “*Candidatus* Villigracilis affinis,” (af.fi´nis. L. masc. adj. affinis, next of kin; indicating the close phylogenetic relationship with *Ca*. Villigracilis saccharophilus). |
| Species status | Candidatus |
| Designation of the type MAG | GCA_016709305.1 |
| MAG/SAG accession number | GCA_016718275.1 |
| Genome status | High-quality draft |
| Genome size | 5057985 |
| GC mol % | 51.39 |
| Country of origin | Denmark |
| Region of origin | Odense |
| Source of sample | Activated sludge |
| Geographical location | Odense |
| Latitude | 55.421534 |
| Longitude | 10.366234 |
| Depth | N/A |
| Altitude | N/A |
| Temperature of the sample | Mesophilic |
| pH of the sample | N/A |
| Relationship to oxygen | Facultative anaerobe |
| Energy metabolism | Potentially utilizing carbohydrates, amino acids, and fatty acids |
| Assembly | 1 sample |
| Sequencing technology | Oxford Nanopore PromethION |
| Binning software used | MaxBin2 |
| Assembly software used | CANU v.1.8 |
| Habitat | Full-scale enriched biological phosphorus removal wastewater treatment plant |
| Miscellaneous, extraordinary features relevant for the description | Filamentous morphology (12-50 × 0.3-0.4 µm) |

**Table 10. Protologue Table for *Candidatus* Villigracilis proximus**

| Species name | *Candidatus* Villigracilis proximus |
| --- | --- |
| Genus name | *Candidatus* Villigracilis |
| Specific epithet | proximus |
| Type species of the genus | *Candidatus* Villigracilis affinis |
| Genus status | Candidatus |
| Species etymology | “*Candidatus* Villigracilis proximus,” (pro´xi.mus. L. masc. adj. proximus, next of kin; indicating the close phylogenetic with *Ca*. Villigracilis saccharophilus). |
| Species status | Candidatus |
| Designation of the type MAG | GCA_016709305.1 |
| MAG/SAG accession number | GCA_016714625.1 |
| Genome status | High-quality draft |
| Genome size | 5019946 |
| GC mol % | 48.91 |
| Country of origin | Denmark |
| Region of origin | Odense |
| Source of sample | Activated sludge |
| Geographical location | Odense |
| Latitude | 55.432604 |
| Longitude | 10.458855 |
| Depth | N/A |
| Altitude | N/A |
| Temperature of the sample | Mesophilic |
| pH of the sample | N/A |
| Relationship to oxygen | Facultative anaerobe |
| Energy metabolism | Potentially utilizing carbohydrates, amino acids, and fatty acids |
| Assembly | 1 sample |
| Sequencing technology | Oxford Nanopore PromethION |
| Binning software used | MaxBin2 |
| Assembly software used | CANU v.1.8 |
| Habitat | Full-scale enriched biological phosphorus removal wastewater treatment plant |
| Miscellaneous, extraordinary features relevant for the description | Filamentous morphology (12-50 × 0.3-0.4 µm) |

**Table 11. Protologue Table for *Candidatus* Villigracilis saccharophilus**

| Species name | *Candidatus* Villigracilis saccharophilus |
| --- | --- |
| Genus name | *Candidatus* Villigracilis |
| Specific epithet | saccharophilus |
| Type species of the genus | *Candidatus* Villigracilis affinis |
| Genus status | Candidatus |
| Species etymology | “*Candidatus* Villigracilis saccharophilus”, (sac.cha.ro´phi.lus. G. neut. n. saccharon, sugar; G. masc. n. philos, lover; N.L. masc. adj. saccharophilus, indicating a preference for sugars as carbon sources). |
| Species status | Candidatus |
| Designation of the type MAG | GCA_016709305.1 |
| MAG/SAG accession number | GCA_016709305.1 |
| Genome status | High-quality draft |
| Genome size | 4851661 |
| GC mol % | 49.87 |
| Country of origin | Denmark |
| Region of origin | Esbjerg |
| Source of sample | Activated sludge |
| Geographical location | Esbjerg |
| Latitude | 55.488097 |
| Longitude | 8.430505 |
| Depth | N/A |
| Altitude | N/A |
| Temperature of the sample | Mesophilic |
| pH of the sample | N/A |
| Relationship to oxygen | Facultative anaerobe |
| Energy metabolism | Potentially utilizing carbohydrates, amino acids, and fatty acids |
| Assembly | 1 sample |
| Sequencing technology | Oxford Nanopore PromethION |
| Binning software used | MaxBin2 |
| Assembly software used | CANU v.1.8 |
| Habitat | Full-scale enriched biological phosphorus removal wastewater treatment plant |
| Miscellaneous, extraordinary features relevant for the description | Filamentous morphology (12-50 × 0.3-0.4 µm) |

Description of “*Candidatus* Hadersleviella danica” gen. nov. sp. nov.: “*Candidatus* Hadersleviella danica”, (Ha.der.sle.viel´la., N.L. fem. n. *Hadersleviella*, named after the city Haderslev where the MAG had been retrieved; da´ni.ca., M.L. fem. adj. *danica*, danish; indicating the country of origin). This taxon is represented by the MAG Hade_MAXAC.236_sub. The complete protologue can be found in Table 12.

**Table 12. Protologue Table for *Candidatus* Hadersleviella danica**

| Species name | *Candidatus* Hadersleviella danica |
| --- | --- |
| Genus name | *Candidatus* Hadersleviella |
| Specific epithet | danica |
| Type species of the genus | *Candidatus* Hadersleviella danica |
| Genus status | Candidatus |
| Species etymology | “*Candidatus* Hadersleviella danica”, (Ha.der.sle.viel´la., N.L. fem. n. *Hadersleviella*, named after the city Haderslev where the MAG had been retrieved; da´ni.ca., M.L. fem. adj. *danica*, danish; indicating the country of origin). |
| Species status | sp. nov. |
| Designation of the type MAG | GCA_016711405.1 |
| MAG/SAG accession number | GCA_016711405.1 |
| Genome status | High-quality draft |
| Genome size | 5674443 |
| GC mol % | 56.34 |
| Country of origin | Denmark |
| Region of origin | Haderslev |
| Source of sample | Activated sludge |
| Geographical location | Haderslev |
| Latitude | 55.249786 |
| Longitude | 9.508609 |
| Depth | N/A |
| Altitude | N/A |
| Temperature of the sample | Mesophilic |
| pH of the sample | N/A |
| Relationship to oxygen | Facultative anaerobe |
| Energy metabolism | Potentially utilizing carbohydrates and amino acids |
| Assembly | 1 sample |
| Sequencing technology | Oxford Nanopore PromethION |
| Binning software used | MaxBin2 |
| Assembly software used | CANU v.1.8 |
| Habitat | Full-scale enriched biological phosphorus removal wastewater treatment plant |
| Miscellaneous, extraordinary features relevant for the description | - |

Description of “*Candidatus* Trichofilum aggregatum” gen. nov. sp. nov.: “*Candidatus* Trichofilum aggregatum”, (Tri.cho.fi´lum., from G. fem. n. *thrix*, hair; L. neut. n. *filum*, thread, filament; N.L. neut. n. *Trichofilum*, indicating the filamentous morphology; ag.gre.ga´tum., L. neut. part. adj. *aggregatum*, indicating the bundles often formed with other filaments). This taxon is represented by the MAG Hirt_MAXAC.142. The complete protologue can be found in Table 13.

**Table 13. Protologue Table for *Candidatus* Trichofilum aggregatum**

| Species name | *Candidatus* Trichofilum aggregatum |
| --- | --- |
| Genus name | *Candidatus* Trichofilum |
| Specific epithet | aggregatum |
| Type species of the genus | *Candidatus* Trichofilum aggregatum |
| Genus status | Candidatus |
| Species etymology | “*Candidatus* Trichofilum aggregatum”, (Tri.cho.fi´lum., from G. fem. n. *thrix*, hair; L. neut. n. *filum*, thread, filament; N.L. neut. n. *Trichofilum*, indicating the filamentous morphology; ag.gre.ga´tum., L. neut. part. adj. *aggregatum*, indicating the bundles often formed with other filaments). |
| Species status | sp. nov. |
| Designation of the type MAG | GCA_016716885.1 |
| MAG/SAG accession number | GCA_016716885.1 |
| Genome status | High-quality draft |
| Genome size | 7738549 |
| GC mol % | 83.42 |
| Country of origin | Denmark |
| Region of origin | Hirtshals |
| Source of sample | Activated sludge |
| Geographical location | Hirtshals |
| Latitude | 57.577275 |
| Longitude | 9.9922971 |
| Depth | N/A |
| Altitude | N/A |
| Temperature of the sample | Mesophilic |
| pH of the sample | N/A |
| Relationship to oxygen | Facultative anaerobe |
| Energy metabolism | Potentially utilizing carbohydrates and amino acids |
| Assembly | 1 sample |
| Sequencing technology | Oxford Nanopore PromethION |
| Binning software used | MaxBin2 |
| Assembly software used | CANU v.1.8 |
| Habitat | Full-scale enriched biological phosphorus removal wastewater treatment plant |
| Miscellaneous, extraordinary features relevant for the description | Filamentous morphology (60-200 × 0.6-0.8 µm) |

Description of ‘*Candidatus* Promineofilum glycogenicum’ sp. nov.: “*Candidatus* Promineofilum glycogenicum”, (gly.co.ge´ni.cum. N.L. neut. adj. *glycogenicum*, indicating the presence of intracellular glycogen). This taxon was represented by the MAG Ega_BAT3C.159. The complete protologue can be found in Table 14.

**Table 14. Protologue Table for *Candidatus* Promineofilum glycogenicum**

| Species name | *Candidatus* Promineofilum glycogenicum |
| --- | --- |
| Genus name | *Candidatus* Promineofilum |
| Specific epithet | glycogenicum |
| Type species of the genus | *Candidatus* Promineofilum breve |
| Genus status | Candidatus |
| Species etymology | “*Candidatus* Promineofilum glycogenicum”, (gly.co.ge´ni.cum. N.L. neut. adj. *glycogenicum*, indicating the presence of intracellular glycogen). |
| Species status | sp. nov. |
| Designation of the type MAG | GCA_900066015.1 |
| MAG/SAG accession number | GCA_016707605.1 |
| Genome status | High-quality draft |
| Genome size | 5044159 |
| GC mol % | 66.77 |
| Country of origin | Denmark |
| Region of origin | Egaa |
| Source of sample | Activated sludge |
| Geographical location | Egaa |
| Latitude | 56.21314 |
| Longitude | 10.242467 |
| Depth | N/A |
| Altitude | N/A |
| Temperature of the sample | Mesophilic |
| pH of the sample | N/A |
| Relationship to oxygen | Facultative anaerobe |
| Energy metabolism | Potentially utilizing carbohydrates, amino acids and fatty acids |
| Assembly | 1 sample |
| Sequencing technology | Oxford Nanopore PromethION |
| Binning software used | MetaBAT2 |
| Assembly software used | CANU v.1.8 |
| Habitat | Full-scale enriched biological phosphorus removal wastewater treatment plant |
| Miscellaneous, extraordinary features relevant for the description | Filamentous morphology (20-140 × 0.8 µm) |

Description of ‘*Candidatus* Leptofilum’ gen. nov.: “*Candidatus* Leptofilum”, (Lep.to.fi´lum. Gr. masc. adj. *leptos*, thin; L. neut. n. *filum*, thread, filament; N.L. neut. n. Leptofilum, indicating a thin filamentous bacterium).

Description of ‘*Candidatus* Leptofilum gracile’ sp. nov.: “*Candidatus* Leptofilum gracile”, (gra´ci.le. L. neut. adj. *gracile*, thin, indicating the thin trichome). This taxon is represented by the MAG Fred_BAT3C.445. The complete protologue can be found in Table 15.

Description of ‘*Candidatus* Leptofilum proximum’ sp. nov.: “*Candidatus* Leptofilum proximum”, (pro´xi.mum. L. neut. adj. *proximum*, next of kin, indicating the close phylogenetic relationship with *Ca*. Leptofilum gracile). This taxon is represented by the MAG Kalu_MAXAC.106v2. The complete protologue can be found in Table 16.

**Table 15. Protologue Table for *Candidatus* Leptofilum gracile**

| Species name | *Candidatus* Leptofilum gracile |
| --- | --- |
| Genus name | *Candidatus* Leptofilum |
| Specific epithet | gracile |
| Type species of the genus | *Candidatus* Leptofilum gracile |
| Genus status | Candidatus |
| Species etymology | “*Candidatus* Leptofilum gracile”, (Lep.to.fi´lum. Gr. masc. adj. *leptos*, thin; L. neut. n. *filum*, thread, filament; N.L. neut. n. *Leptofilum*, indicating a thin filamentous bacterium; gra´ci.le. L. neut. adj. *gracile*, thin, indicating the thin trichome). |
| Species status | Candidatus |
| Designation of the type MAG | GCA_016713825.1 |
| MAG/SAG accession number | GCA_016713825.1 |
| Genome status | High-quality draft |
| Genome size | 5432132 |
| GC mol % | 56.03 |
| Country of origin | Denmark |
| Region of origin | Fredericia |
| Source of sample | Activated sludge |
| Geographical location | Fredericia |
| Latitude | 55.552368 |
| Longitude | 9.720404 |
| Depth | N/A |
| Altitude | N/A |
| Temperature of the sample | Mesophilic |
| pH of the sample | N/A |
| Relationship to oxygen | Facultative anaerobe |
| Energy metabolism | Potentially utilizing carbohydrates, amino acids, acetate |
| Assembly | 1 sample |
| Sequencing technology | Oxford Nanopore PromethION |
| Binning software used | MetaBAT2 |
| Assembly software used | CANU v.1.8 |
| Habitat | Full-scale enriched biological phosphorus removal wastewater treatment plant |
| Miscellaneous, extraordinary features relevant for the description | Filamentous morphology (10-70 × 0.7-0.9 µm) |

**Table 16. Protologue Table for *Candidatus* Leptofilum proximum**

| Species name | *Candidatus* Leptofilum proximum |
| --- | --- |
| Genus name | *Candidatus* Leptofilum |
| Specific epithet | proximum |
| Type species of the genus | *Candidatus* Leptofilum gracile |
| Genus status | Candidatus |
| Species etymology | “*Candidatus* Leptofilum gracile”, (Lep.to.fi´lum. Gr. masc. adj. *leptos*, thin; L. neut. n. *filum*, thread, filament; N.L. neut. n. *Leptofilum*, indicating a thin filamentous bacterium; pro´xi.mum. L. neut. adj. *proximum*, next of kin, indicating the close phylogenetic relationship with *Ca.* Leptofilum gracile). This taxon is represented by the MAG Kalu_MAXAC.106v2. |
| Species status | Candidatus |
| Designation of the type MAG | GCA_016713825.1 |
| MAG/SAG accession number | GCA_016710325.1 |
| Genome status | High-quality draft |
| Genome size | 8146679 |
| GC mol % | 56.31 |
| Country of origin | Denmark |
| Region of origin | Kalundborg |
| Source of sample | Activated sludge |
| Geographical location | Kalundborg |
| Latitude | 55.668006 |
| Longitude | 11.107247 |
| Depth | N/A |
| Altitude | N/A |
| Temperature of the sample | Mesophilic |
| pH of the sample | N/A |
| Relationship to oxygen | Facultative anaerobe |
| Energy metabolism | Potentially utilizing carbohydrates, amino acids, acetate |
| Assembly | 1 sample |
| Sequencing technology | Oxford Nanopore PromethION |
| Binning software used | MaxBin2 |
| Assembly software used | CANU v.1.8 |
| Habitat | Full-scale enriched biological phosphorus removal wastewater treatment plant |
| Miscellaneous, extraordinary features relevant for the description | Filamentous morphology (10-70 × 0.7-0.9 µm) |

Description of ‘*Candidatus* Leptovillus’ gen. nov.: “*Candidatus* Leptovillus”, (Lep.to.vil´lus, Gr. masc. adj. *leptos*, thin; L. masc. n. *villus*, hair, filament; N.L. masc. n. *Leptovillus*, indicating a thin filamentous bacterium).

Description of ‘*Candidatus* Leptovillus gracilis’ sp. nov.: “*Candidatus* Leptovillus gracilis (gra´ci.lis. L. masc. adj. *gracilis*, thin, indicating the thin trichome). This taxon is represented by the MAG Kalu_BATAC.47. The complete protologue can be found in Table 17.

Description of ‘*Candidatus* Leptovillus affinis’ sp. nov.: “*Candidatus* Leptovillus affinis”, (af.fi´nis. L. masc. adj. *affinis*, next of kin, indicating the close phylogenetic relationship with *Ca*. Leptovillus gracilis). This taxon is represented by the MAG AalE_BATAC.251. The complete protologue can be found in Table 18.

**Table 17. Protologue Table for *Candidatus* Leptovillus gracilis**

| Species name | *Candidatus* Leptovillus gracilis |
| --- | --- |
| Genus name | *Candidatus* Leptovillus |
| Specific epithet | gracilis |
| Type species of the genus | *Candidatus* Leptovillus affinis |
| Genus status | Candidatus |
| Species etymology | “*Candidatus* Leptovillus gracilis”, (Lep.to.vil´lus, Gr. masc. adj. *leptos*, thin; L. masc. n. *villus*, hair, filament; N.L. masc. n. *Leptovillus*, indicating a thin filamentous bacterium; gra´ci.lis. L. masc. adj. *gracilis*, thin, indicating the thin trichome). |
| Species status | Candidatus |
| Designation of the type MAG | GCA_016716065.1 |
| MAG/SAG accession number | GCA_016716065.1 |
| Genome status | High-quality draft |
| Genome size | 6811151 |
| GC mol % | 57.32 |
| Country of origin | Denmark |
| Region of origin | Kalundborg |
| Source of sample | Activated sludge |
| Geographical location | Kalundborg |
| Latitude | 55.668006 |
| Longitude | 11.107247 |
| Depth | N/A |
| Altitude | N/A |
| Temperature of the sample | Mesophilic |
| pH of the sample | N/A |
| Relationship to oxygen | Facultative anaerobe |
| Energy metabolism | Potentially utilizing carbohydrates, amino acids, acetate and fatty acids |
| Assembly | 1 sample |
| Sequencing technology | Oxford Nanopore PromethION |
| Binning software used | MetaBAT2 |
| Assembly software used | CANU v.1.8 |
| Habitat | Full-scale enriched biological phosphorus removal wastewater treatment plant |
| Miscellaneous, extraordinary features relevant for the description | Filamentous morphology (10-70 × 0.7-0.9 µm) |

**Table 18. Protologue Table for *Candidatus* Leptovillus affinis**

| Species name | *Candidatus* Leptovillus affinis |
| --- | --- |
| Genus name | *Candidatus* Leptovillus |
| Specific epithet | affinis |
| Type species of the genus | *Candidatus* Leptovillus affinis |
| Genus status | Candidatus |
| Species etymology | “Candidatus Leptovillus affinis”, (Lep.to.vil´lus, Gr. masc. adj. *leptos*, thin; L. masc. n. *villus*, hair, filament; N.L. masc. n. *Leptovillus*, indicating a thin filamentous bacterium; af.fi´nis. L. masc. adj. *affinis*, next of kin, indicating the close phylogenetic relationship with *Ca.* Leptovillus gracilis). |
| Species status | Candidatus |
| Designation of the type MAG | GCA_016716065.1 |
| MAG/SAG accession number | GCA_016705235.1 |
| Genome status | High-quality draft |
| Genome size | 5709216 |
| GC mol % | 56.75 |
| Country of origin | Denmark |
| Region of origin | Aalborg |
| Source of sample | Activated sludge |
| Geographical location | Aalborg |
| Latitude | 57.045161 |
| Longitude | 10.045761 |
| Depth | N/A |
| Altitude | N/A |
| Temperature of the sample | Mesophilic |
| pH of the sample | N/A |
| Relationship to oxygen | Facultative anaerobe |
| Energy metabolism | Potentially utilizing carbohydrates, amino acids, acetate and fatty acids |
| Assembly | 1 sample |
| Sequencing technology | Oxford Nanopore PromethION |
| Binning software used | MetaBAT2 |
| Assembly software used | CANU v.1.8 |
| Habitat | Full-scale enriched biological phosphorus removal wastewater treatment plant |
| Miscellaneous, extraordinary features relevant for the description | Filamentous morphology (10-70 × 0.7-0.9 µm) |

Description of “*Candidatus* Flexicrinis” gen. nov.: “*Candidatus* Flexincrinis”, (Fle.xi.cri´nis., from L. masc. part. *flexus*, bent; L. masc. n. *crinis,* hair, here interpreted as filaments; N.L. masc. n. *Flexicrinis*, indicating the bent shape of the filaments).

Description of “*Candidatus* Flexicrinis affinis” gen. nov. sp. nov. (af.fi´nis., L. masc. adj. *affinis*, next of kin; indicating the close phylogenetic relationship with *Ca.* Flexicrinis proximus). This taxon is represented by the MAG Kalu_BAT3C.186. The complete protologue can be found in Table 19.

Description of “*Candidatus* Flexicrinis proximus” gen. nov. sp. nov.: (pro´xi.mus., L. masc. adj. *proximus*, next of kin; indicating the close phylogenetic relationship with *Ca.* Flexicrinis affinis). This taxon is represented by the MAG Fred_MAXAC.112. The complete protologue can be found in Table 20.

**Table 19. Protologue Table for *Candidatus* Flexicrinis affinis**

| Species name | *Candidatus* Flexicrinis affinis |
| --- | --- |
| Genus name | *Candidatus* Flexicrinis |
| Specific epithet | affinis |
| Type species of the genus | *Candidatus* Flexicrinis affinis |
| Genus status | Candidatus |
| Species etymology | *Candidatus* Flexicrinis affinis” gen. nov. sp. nov.: “*Candidatus* Flexincrinis”, (Fle.xi.cri´nis., from L. masc. part. *flexus*, bent; L. masc. n. *crinis*, hair, here interpreted as filaments; N.L. masc. n. *Flexicrinis*, indicating the bent shape of the filaments; af.fi´nis., L. masc. adj. *affinis*, next of kin; indicating the close phylogenetic relationship with *Ca.* Flexicrinis proximus). |
| Species status | sp. nov. |
| Designation of the type MAG | GCA_016716525.1 |
| MAG/SAG accession number | GCA_016716525.1 |
| Genome status | High-quality draft |
| Genome size | 5503990 |
| GC mol % | 62.60 |
| Country of origin | Denmark |
| Region of origin | Kalundborg |
| Source of sample | Activated sludge |
| Geographical location | Kalundborg |
| Latitude | 55.668006 |
| Longitude | 11.107247 |
| Depth | N/A |
| Altitude | N/A |
| Temperature of the sample | Mesophilic |
| pH of the sample | N/A |
| Relationship to oxygen | Facultative anaerobe |
| Energy metabolism | Potentially utilizing carbohydrates and amino acids |
| Assembly | 1 sample |
| Sequencing technology | Oxford Nanopore PromethION |
| Binning software used | MetaBAT2 |
| Assembly software used | CANU v.1.8 |
| Habitat | Full-scale enriched biological phosphorus removal wastewater treatment plant |
| Miscellaneous, extraordinary features relevant for the description | Filamentous morphology (40-110 × 0.7-1.1µm) |

**Table 20. Protologue Table for *Candidatus* Flexicrinis proximus**

| Species name | *Candidatus* Flexicrinis proximus |
| --- | --- |
| Genus name | *Candidatus* Flexicrinis |
| Specific epithet | proximus |
| Type species of the genus | *Candidatus* Flexicrinis proximus |
| Genus status | Candidatus |
| Species etymology | *Candidatus* Flexicrinis proximus” gen. nov. sp, nov.: “*Candidatus* Flexincrinis”, (Fle.xi.cri´nis., from L. masc. part. *flexus*, bent; L. masc. n. *crinis*, hair, here interpreted as filaments; N.L. masc. n. *Flexicrinis*, indicating the bent shape of the filaments; pro´xi.mus., L. masc. adj. *proximus*, next of kin; indicating the close phylogenetic relationship with *Ca*. Flexicrinis affinis). |
| Species status | sp. nov. |
| Designation of the type MAG | GCA_016716525.1 |
| MAG/SAG accession number | GCA_016712885.1 |
| Genome status | High-quality draft |
| Genome size | 6159798 |
| GC mol % | 60.03 |
| Country of origin | Denmark |
| Region of origin | Fredericia |
| Source of sample | Activated sludge |
| Geographical location | Fredericia |
| Latitude | 55.552368 |
| Longitude | 9.720404 |
| Depth | N/A |
| Altitude | N/A |
| Temperature of the sample | Mesophilic |
| pH of the sample | N/A |
| Relationship to oxygen | Facultative anaerobe |
| Energy metabolism | Potentially utilizing carbohydrates and amino acids |
| Assembly | 1 sample |
| Sequencing technology | Oxford Nanopore PromethION |
| Binning software used | MaxBin2 |
| Assembly software used | CANU v.1.8 |
| Habitat | Full-scale enriched biological phosphorus removal wastewater treatment plant |
| Miscellaneous, extraordinary features relevant for the description | Filamentous morphology (40-110 × 0.7-1.1µm) |

Description of “*Candidatus* Flexifilum” gen. nov. sp. nov.: “*Candidatus* Flexifilum”, (Fle.xi.fi´lum., from L. masc. part. *flexus*, bent; L. neut. n. *filum*, thread, filament; N.L. neut. n. *Flexifilum*, indicating the often bent filaments)

Description of “*Candidatus* Flexifilum breve” gen. nov. sp. nov.: (bre´ve., L. neut. adj. *breve*, short, indicating the short length of the filaments). This taxon is represented by the MAG Ribe_BATAC.253. The complete protologue can be found in Table 21.

Description of “*Candidatus* Flexifilum affine” gen. nov. sp. nov.: “*Candidatus* Flexifilum”, (af.fi´ne., L. neut. adj. *affinis*, next of kin; indicating the close phylogenetic relationship with *Ca.* Flexifilum breve). This taxon is represented by the MAG Fred_BATAC.421. The complete protologue can be found in Table 22.

Description of Flexifilaceae fam. nov. (Fle.xi.fi.la.ce´ae. from N.L. fem. n. Flexifilum type genus of the family; suff. -aceae ending to denote a family; N.L. fem. pl. n. Flexifilaceae, the Flexifilum family).

**Table 21. Protologue Table for *Candidatus* Flexifilum breve**

| Species name | *Candidatus* Flexifilum breve |
| --- | --- |
| Genus name | *Candidatus* Flexifilum |
| Specific epithet | breve |
| Type species of the genus | *Candidatus* Flexifilum breve |
| Genus status | Candidatus |
| Species etymology | *Candidatus* Flexifilum breve” gen. nov. sp. nov.: “*Candidatus* Flexifilum breve”, (Fle.xi.fi´lum., from L. masc. part. *flexus*, bent; L. neut. n. *filum*, thread, filament; N.L. neut. n. *Flexifilum*, indicating the often bent filaments; bre´ve., L. neut. adj. *breve*, short, indicating the short length of the filaments). |
| Species status | sp. nov. |
| Designation of the type MAG | GCA_016717205.1 |
| MAG/SAG accession number | GCA_016717205.1 |
| Genome status | High-quality draft |
| Genome size | 7801835 |
| GC mol % | 59.58 |
| Country of origin | Denmark |
| Region of origin | Ribe |
| Source of sample | Activated sludge |
| Geographical location | Ribe |
| Latitude | 55.329053 |
| Longitude | 8.74336 |
| Depth | N/A |
| Altitude | N/A |
| Temperature of the sample | Mesophilic |
| pH of the sample | N/A |
| Relationship to oxygen | Facultative anaerobe |
| Energy metabolism | Potentially utilizing carbohydrates and amino acids |
| Assembly | 1 sample |
| Sequencing technology | Oxford Nanopore PromethION |
| Binning software used | MetaBAT2 |
| Assembly software used | CANU v.1.8 |
| Habitat | Full-scale enriched biological phosphorus removal wastewater treatment plant |
| Miscellaneous, extraordinary features relevant for the description | Filamentous morphology (>100 × 0.8-1.1 µm) |

**Table 22. Protologue Table for *Candidatus* Flexifilum affine**

| Species name | *Candidatus* Flexifilum affine |
| --- | --- |
| Genus name | *Candidatus* Flexifilum |
| Specific epithet | affinis |
| Type species of the genus | *Candidatus* Flexifilum affine |
| Genus status | Candidatus |
| Species etymology | *Candidatus* Flexifilum affine” gen. nov. sp. nov.: “*Candidatus* Flexifilum affine”, (Fle.xi.fi´lum., from L. masc. part. *flexus*, bent; L. neut. n. *filum*, thread, filament; N.L. neut. n. *Flexifilum*, indicating the often bent filaments; af.fi´ne., L. neut. adj. *affine*, next of kin; indicating the close phylogenetic relationship with *Ca*. Flexifilum breve). |
| Species status | sp. nov. |
| Designation of the type MAG | GCA_016717205.1 |
| MAG/SAG accession number | GCA_016713325.1 |
| Genome status | High-quality draft |
| Genome size | 9033822 |
| GC mol % | 62.18 |
| Country of origin | Denmark |
| Region of origin | Fredericia |
| Source of sample | Activated sludge |
| Geographical location | Fredericia |
| Latitude | 55.552368 |
| Longitude | 9.720404 |
| Depth | N/A |
| Altitude | N/A |
| Temperature of the sample | Mesophilic |
| pH of the sample | N/A |
| Relationship to oxygen | Facultative anaerobe |
| Energy metabolism | Potentially utilizing carbohydrates and amino acids |
| Assembly | 1 sample |
| Sequencing technology | Oxford Nanopore PromethION |
| Binning software used | MetaBAT2 |
| Assembly software used | CANU v.1.8 |
| Habitat | Full-scale enriched biological phosphorus removal wastewater treatment plant |
| Miscellaneous, extraordinary features relevant for the description | Filamentous morphology (>100 × 0.8-1.1 µm) |

Description of ‘*Candidatus* Amarolinea dominans’ sp. nov. “*Candidatus* Amarolinea dominans”, (do´mi.nans. L. part. adj. *dominans*, dominant, indicating the high abundance in sewage systems). This taxon is represented by the MAG Lyne_BATAC.272. The complete protologue can be found in Table 23.

**Table 23. Protologue Table for *Candidatus* Amarolinea dominans**

| Species name | *Candidatus* Amarolinea dominans |
| --- | --- |
| Genus name | *Candidatus* Amarolinea |
| Specific epithet | dominans |
| Type species of the genus | *Candidatus* Amarolinea aalborgensis |
| Genus status | Candidatus |
| Species etymology | “*Candidatus* Amarolinea dominans”, (do´mi.nans. L. part. adj. *dominans*, dominant, indicating the high abundance in sewage systems). |
| Species status | sp. nov. |
| Designation of the type MAG | GCA_900491745.1 |
| MAG/SAG accession number | GCA_016719785.1 |
| Genome status | High-quality draft |
| Genome size | 6591815 |
| GC mol % | 62.59 |
| Country of origin | Denmark |
| Region of origin | Lynetten |
| Source of sample | Activated sludge |
| Geographical location | Lyne |
| Latitude | 55.695362 |
| Longitude | 12.612702 |
| Depth | N/A |
| Altitude | N/A |
| Temperature of the sample | Mesophilic |
| pH of the sample | N/A |
| Relationship to oxygen | Facultative anaerobe |
| Energy metabolism | Potentially utilizing carbohydrates, fatty acids and amino acids |
| Assembly | 1 sample |
| Sequencing technology | Oxford Nanopore PromethION |
| Binning software used | MetaBAT2 |
| Assembly software used | CANU v.1.8 |
| Habitat | Full-scale enriched biological phosphorus removal wastewater treatment plant |
| Miscellaneous, extraordinary features relevant for the description | Filamentous morphology (20-140 × 2.2 µm) |

Description of “*Candidatus* Fredericiella danica” gen. nov. sp. nov.: “*Candidatus* Fredericiella danica”, (Fre.de.ri.ciel´la., N.L. fem. n. *Fredericiella*, named after the city Fredericia where the MAG had been retrieved; da´ni.ca. M.L. fem. adj. *danica*, danish; indicating the country of origin). This taxon is represented by the MAG Fred_BATAC.359. The complete protologue can be found in Table 24.

**Table 24. Protologue Table for *Candidatus* Fredericiella danica**

| Species name | *Candidatus* Fredericiella danica |
| --- | --- |
| Genus name | *Candidatus* Fredericiella |
| Specific epithet | danica |
| Type species of the genus | *Candidatus* Fredericiella danica |
| Genus status | Candidatus |
| Species etymology | *Candidatus* Fredericiella danica” gen. nov. sp. nov.: “*Candidatus* Fredericiella danica”, (Fre.de.ri.ciel´la., N.L. fem. n. *Fredericiella*, named after the city Fredericia where the MAG had been retrieved; da´ni.ca. M.L. fem. adj. *danica*, danish; indicating the country of origin). |
| Species status | sp. nov. |
| Designation of the type MAG | GCA_016713335.1 |
| MAG/SAG accession number | GCA_016713335.1 |
| Genome status | High-quality draft |
| Genome size | 6824975 |
| GC mol % | 59.48 |
| Country of origin | Denmark |
| Region of origin | Fredericia |
| Source of sample | Activated sludge |
| Geographical location | Fredericia |
| Latitude | 55.552368 |
| Longitude | 9.720404 |
| Depth | N/A |
| Altitude | N/A |
| Temperature of the sample | Mesophilic |
| pH of the sample | N/A |
| Relationship to oxygen | Facultative anaerobe |
| Energy metabolism | Potentially utilizing carbohydrates and amino acids |
| Assembly | 1 sample |
| Sequencing technology | Oxford Nanopore PromethION |
| Binning software used | MetaBAT2 |
| Assembly software used | CANU v.1.8 |
| Habitat | Full-scale enriched biological phosphorus removal wastewater treatment plant |
| Miscellaneous, extraordinary features relevant for the description |  |

Description of ‘*Candidatus* Caldilinea saccharophila’ sp. nov. “*Candidatus* Caldilinea saccharophila”, (sac.cha.ro´phi.la. Gr. neut. n. *saccharon*, sugar; Gr. masc. n. *philos*, lover; N.L. fem. adj. *saccharophila*, indicating a preference for sugars as carbon sources). This taxon is represented by the MAG Hjor_MAXAC.079_sub. The complete protologue can be found in Table 25.

**Table 25. Protologue Table for *Candidatus* Caldilinea saccharophila**

| Species name | *Candidatus* Caldilinea saccharophila |
| --- | --- |
| Genus name | *Caldilinea* |
| Specific epithet | saccharophila |
| Type species of the genus | *Caldilinea aerophila* |
| Genus status | Validly published (Taxonomy ID: 926550) |
| Species etymology | “*Candidatus* Caldilinea saccharophila”, (sac.cha.ro´phi.la. Gr. neut. n. *saccharon*, sugar; Gr. masc. n. *philos*, lover; N.L. fem. adj. *saccharophila*, indicating a preference for sugars as carbon sources). |
| Species status | sp. nov. |
| Designation of the type MAG | GCA_000281175.1 |
| MAG/SAG accession number | GCA_016710365.1 |
| Genome status | High-quality draft |
| Genome size | 6608906 |
| GC mol % | 60.66 |
| Country of origin | Denmark |
| Region of origin | Hjørring |
| Source of sample | Activated sludge |
| Geographical location | Hjørring |
| Latitude | 57.421265 |
| Longitude | 9.975411 |
| Depth | N/A |
| Altitude | N/A |
| Temperature of the sample | Mesophilic |
| pH of the sample | N/A |
| Relationship to oxygen | Facultative anaerobe |
| Energy metabolism | Potentially utilizing carbohydrates, amino acids and fatty acids |
| Assembly | 1 sample |
| Sequencing technology | Oxford Nanopore PromethION |
| Binning software used | MaxBin2 |
| Assembly software used | CANU v.1.8 |
| Habitat | Full-scale enriched biological phosphorus removal wastewater treatment plant |
| Miscellaneous, extraordinary features relevant for the description | Filamentous morphology (70-200 × 0.8 ± 0.2 µm) |

Description of “*Candidatus* Ribeiella danica” gen. nov. sp. nov.: “*Candidatus* Ribeiella”, (Ri.be.i.el’la., N.L. fem. n. *Ribeiella*, named after the city Ribe where the MAG had been retrieved; da´ni.ca., M.L. fem. adj. *danica*, danish; indicating the country of origin). This taxon is represented by the MAG Ribe_BAT3C.183. The complete protologue can be found in Table 26.

**Table 26. Protologue Table for *Candidatus* Ribeiella danica**

| Species name | *Candidatus* Ribeiella danica |
| --- | --- |
| Genus name | *Candidatus* Ribeiella |
| Specific epithet | danica |
| Type species of the genus | *Candidatus* Ribeiella danica |
| Genus status | Candidatus |
| Species etymology | *Candidatus* Ribeiella danica” gen. nov. sp. nov.: “*Candidatus* Ribeiella danica”, (Ri.be.i.el’la., N.L. fem. n. Ribeiella, named after the city Ribe where the MAG had been retrieved; da´ni.ca. M.L. fem. adj. *danica*, danish; indicating the country of origin). |
| Species status | sp. nov. |
| Designation of the type MAG | GCA_016717335.1 |
| MAG/SAG accession number | GCA_016717335.1 |
| Genome status | High-quality draft |
| Genome size | 8174342 |
| GC mol % | 67.95 |
| Country of origin | Denmark |
| Region of origin | Ribe |
| Source of sample | Activated sludge |
| Geographical location | Ribe |
| Latitude | 55.329053 |
| Longitude | 8.74336 |
| Depth | N/A |
| Altitude | N/A |
| Temperature of the sample | Mesophilic |
| pH of the sample | N/A |
| Relationship to oxygen | Facultative anaerobe |
| Energy metabolism | Potentially utilizing carbohydrates and amino acids |
| Assembly | 1 sample |
| Sequencing technology | Oxford Nanopore PromethION |
| Binning software used | MetaBAT2 |
| Assembly software used | CANU v.1.8 |
| Habitat | Full-scale enriched biological phosphorus removal wastewater treatment plant |
| Miscellaneous, extraordinary features relevant for the description |  |

Description of ‘*Candidatus* Kouleothrix ribensis’ sp. nov. “*Candidatus* Kouleothrix ribensis”, (ri.ben´sis. N.L. fem. adj. *ribensis*, pertinent to the city of Ribe, from where the MAG was obtained). This taxon is represented by the MAG Ribe_MAXAC.235. The complete protologue can be found in Table 27.

**Table 27. Protologue Table for *Candidatus* Kouleothrix ribensis**

| Species name | *Candidatus* Kouleothrix ribensis |
| --- | --- |
| Genus name | *Kouleothrix* |
| Specific epithet | ribensis |
| Type species of the genus | *Kouleothrix auriantiaca* |
| Genus status | Validly published (Taxonomy ID: 186475) |
| Species etymology | “*Candidatus* Kouleothrix ribensis”, (ri.ben´sis. N.L. fem. adj. *ribensis*, pertinent to the city of Ribe, from where the MAG was obtained). |
| Species status | sp. nov. |
| Designation of the type MAG | GCA_001399705.1 |
| MAG/SAG accession number | GCA_016722075.1 |
| Genome status | High-quality draft |
| Genome size | 7093236 |
| GC mol % | 64.99 |
| Country of origin | Denmark |
| Region of origin | Ribe |
| Source of sample | Activated sludge |
| Geographical location | Ribe |
| Latitude | 55.329053 |
| Longitude | 8.74336 |
| Depth | N/A |
| Altitude | N/A |
| Temperature of the sample | Mesophilic |
| pH of the sample | N/A |
| Relationship to oxygen | Facultative anaerobe |
| Energy metabolism | Potentially utilizing carbohydrates, amino acids, acetate and fatty acids |
| Assembly | 1 sample |
| Sequencing technology | Oxford Nanopore PromethION |
| Binning software used | MaxBin2 |
| Assembly software used | CANU v.1.8 |
| Habitat | Full-scale enriched biological phosphorus removal wastewater treatment plant |
| Miscellaneous, extraordinary features relevant for the description | Filamentous morphology (>200 × 0.5-0.7 µm) |

Description of ‘*Candidatus* Amarobacter glycogenicus’ gen. nov. sp. nov. “*Candidatus* Amarobacter glycogenicus”, (A.ma.ro.bac´ter. Gr. fem. n*. amara*, trench, conduit, channel, here, a sewage duct; N.L. masc. n. *bacter*, rod-shaped bacterium; N.L. masc. n. *Amarobacter*, indicating a rod-shaped bacterium found in sewage; gly.co.ge´ni.cus. N.L. masc. adj. *glycogenicus*, indicating the presence of intracellular glycogen). This taxon is represented by the MAG Lyne_MAXAC.019. The complete protologue can be found in Table 28.

**Table 28. Protologue Table for *Candidatus* Amarobacter glycogenicus**

| Species name | *Candidatus* Amarobacter glycogenicus |
| --- | --- |
| Genus name | *Candidatus* Amarobacter |
| Specific epithet | glycogenicus |
| Type species of the genus | *Candidatus* Amarobacter glycogenicus |
| Genus status | Candidatus |
| Species etymology | “*Candidatus* Amarobacter glycogenicus”, (A.ma.ro.bac´ter. Gr. fem. n. *amara*, trench, conduit, channel, here, a sewage duct; N.L. masc. n. *bacter*, rod-shaped bacterium; N.L. masc. n. Amarobacter, indicating a rod-shaped bacterium found in sewage; gly.co.ge´ni.cus. N.L. masc. adj. *glycogenicus*, indicating the presence of intracellular glycogen). |
| Species status | sp. nov. |
| Designation of the type MAG | GCA_016719395.1 |
| MAG/SAG accession number | GCA_016719395.1 |
| Genome status | High-quality draft |
| Genome size | 3446936 |
| GC mol % | 65.53 |
| Country of origin | Denmark |
| Region of origin | Lynetten |
| Source of sample | Activated sludge |
| Geographical location | Lynetten |
| Latitude | 55.695362 |
| Longitude | 12.612702 |
| Depth | N/A |
| Altitude | N/A |
| Temperature of the sample | Mesophilic |
| pH of the sample | N/A |
| Relationship to oxygen | Facultative anaerobe |
| Energy metabolism | Potentially utilizing long chain fatty acids, carbohydrates and amino acids |
| Assembly | 1 sample |
| Sequencing technology | Oxford Nanopore PromethION |
| Binning software used | MaxBin2 |
| Assembly software used | Canu v.1.8 |
| Habitat | Full-scale enriched biological phosphorus removal wastewater treatment plant |
| Miscellaneous, extraordinary features relevant for the description | Rod-shaped morphology (1-2 × 0.3-0.5 µm) |

Description of ‘*Candidatus* Amarobacillus elongatus gen. nov. sp. nov. “*Candidatus* Amarobacillus elongatus”, (A.ma.ro.ba.cil´lus. G. fem. n. *amara*, trench, conduit, channel, here, a sewage duct; L. masc. n. *bacillus*, rod-shaped bacterium; N.L. masc. n. Amarobacillus, indicating a rod-shaped bacterium found in sewage; e.lon.ga´tus. L. masc. adj. *elongatus*, elongated, indicating the elongated shape). This taxon is represented by the MAG Aved_BAT3C.689. The complete protologue can be found in Table 29.

**Table 29. Protologue Table for *Candidatus* Amarobacillus elongatus**

| Species name | *Candidatus* Amarobacillus elongatus |
| --- | --- |
| Genus name | *Candidatus* Amarobacillus |
| Specific epithet | elongatus |
| Type species of the genus | *Candidatus* Amarobacillus elongatus |
| Genus status | Candidatus |
| Species etymology | “*Candidatus* Amarobacillus elongatus”, (A.ma.ro.ba.cil´lus. G. fem. n. *amara*, trench, conduit, channel, here, a sewage duct; L. masc. n. *bacillus*, rod-shaped bacterium; N.L. masc. n. *Amarobacillus*, indicating a rod-shaped bacterium found in sewage; e.lon.ga´tus. L. masc. adj. *elongatus*, elongated, indicating the elongated shape). |
| Species status | sp. nov. |
| Designation of the type MAG | GCA_016703545.1 |
| MAG/SAG accession number | GCA_016703545.1 |
| Genome status | High-quality draft |
| Genome size | 3490229 |
| GC mol % | 69.25 |
| Country of origin | Denmark |
| Region of origin | Avedøre |
| Source of sample | Activated sludge |
| Geographical location | Avedøre |
| Latitude | 55.608613 |
| Longitude | 12.450537 |
| Depth | N/A |
| Altitude | N/A |
| Temperature of the sample | Mesophilic |
| pH of the sample | N/A |
| Relationship to oxygen | Facultative anaerobe |
| Energy metabolism | Potentially utilizing long chain fatty acids, carbohydrates and amino acids |
| Assembly | 1 sample |
| Sequencing technology | Oxford Nanopore PromethION |
| Binning software used | MetaBAT2 |
| Assembly software used | CANU v.1.8 |
| Habitat | Full-scale enriched biological phosphorus removal wastewater treatment plant |
| Miscellaneous, extraordinary features relevant for the description | Rod-shaped morphology (1-2 × 0.3-0.5 µm) |

Description of “*Candidatus* Amarofilum” gen. nov. “*Candidatus* Amarofilum” (midas_g_391), (A.ma.ro.fi´lum., from Gr. fem. n*. amara*, trench, conduit, channel, here, a sewage duct; L. neut. n. *filum*, thread, filament; N.L. neut. n. Amarofilum, indicating a filamentous bacterium found in sewage).

**Table 30. Items for inclusion in the codified record of provisional taxon for “*Candidatus* Amarofilum”.**

| **Attribute** | **Specifications** |
| --- | --- |
| Status | *Candidatus* |
| Vernacular epithet | “another” |
| Phylogenetic lineage or possible genus | Genus Amarofilum in the family *Roseiflexaceae* |
| Cultivation | Not cultivated |
| Gram reaction | Unknown |
| Morphology | Filamentous |
| Basis of assignment | FL-ASV391 in MiDAS4 database (1) |
| Specified identification of morphotype | Oligonucleotide sequences complementary to unique region of 16S rRNA:  CFX122: 5´- CTT GGG CAC ATT CCC ACG T-3´  used with competitor  CFX122­_C1: 5´-TTT GGG CAC ATT CCC ACG C-3´ |
| Habitat, association, or host | Activated sludge |
| Metabolism and unusual features | Unknown |
| Growth temperature | Mesophilic |
| Source | Full-scale biological nutrient removal wastewater treatment plant |
| Author(s) | This study |

Description of “*Candidatus* Pachofilum” gen. nov. “*Candidatus* Pachofilum” (midas_g_550), (Pa.cho.fi´lum., from Gr. masc. adj. *pachys*, thick; L. neut. n. *filum*, thread, filament; N.L. neut. n. *Pachofilum*, indicating a thick filament).

**Table 33. Items for inclusion in the codified record of provisional taxon for “*Candidatus* Pachofilum”.**

| **Attribute** | **Specifications** |
| --- | --- |
| Status | *Candidatus* |
| Vernacular epithet | “another” |
| Phylogenetic lineage or possible genus | Genus Pachofilum in the family midas_f_105, order 1-20, class Anaerolineae |
| Cultivation | Not cultivated |
| Gram reaction | Unknown |
| Morphology | Filamentous |
| Basis of assignment | FL-ASV550 in MiDAS 4 database (1) |
| Specified identification of morphotype | Oligonucleotide sequences complementary to unique region of 16S rRNA:  CFX682: 5´- ATC TAC ATA TTC CAC CAT TAC ACC-3´  used with helper probes:  CFX682­_H1: 5´- TCC GCA TTC CTC TCA TYG CC -3´  CFX682­_H2: 5´- CTT TCG CAC ATG AGC GTC AGG C-3´ |
| Habitat, association, or host | Activated sludge |
| Metabolism and unusual features | Unknown |
| Growth temperature | Mesophilic |
| Source | Full-scale biological nutrient removal wastewater treatment plant |
| Author(s) | This study |

Description of “*Candidatus* Tricholinea” gen. nov. “*Candidatus* Tricholinea” (midas_g_9648), (Tri.cho.li´nea., from G. fem. n. *thrix*, hair; L. fem. n. *linea,* line, filament; N.L. neut. n. *Tricholinea*, indicating a bacterium with filamentous morphology).

**Table 33. Items for inclusion in the codified record of provisional taxon for “*Candidatus* Tricholinea”.**

| **Attribute** | **Specifications** |
| --- | --- |
| Status | *Candidatus* |
| Vernacular epithet | “another” |
| Phylogenetic lineage or possible genus | Genus Tricholinea in the family *Anaerolineaceae* |
| Cultivation | Not cultivated |
| Gram reaction | Unknown |
| Morphology | Filamentous |
| Basis of assignment | FL-ASV9648 in MiDAS 4 database (1) |
| Specified identification of morphotype | Oligonucleotide sequences complementary to unique region of 16S rRNA:  CFX166: 5´- GTA ACY TCA TGC GGT ATT AGC AG -3´  used with helper probes:  CFX166­_H1: 5´- GCA GGT CAC CAA CGC GTT ACT C-3´  CFX166­_H2: 5´- GCT GAT GGG ACG CAG GCC CCT CC -3´ |
| Habitat, association, or host | Activated sludge |
| Metabolism and unusual features | Unknown |
| Growth temperature | Mesophilic |
| Source | Full-scale biological nutrient removal wastewater treatment plant |
| Author(s) | This study |

Rename of *Ca*. Villigracilis nielsenii (2) to “Candidatus Manresella nielsenii” gen. nov. sp. nov.: “Candidatus Manresella”, (Man.re.sel’la., N.L. fem. n. Manresella, named after the city Manresa, origin of the sludge from where the MAG had been retrieved; niel.se’ni.i. N.L. gen. n. nielsenii, of Nielsen, named in honor of Per Halkjær Nielsen, the Danish environmental microbiologist who made important contributions to research and practices in the field of Microbial Ecology and Water Engineering, as suggested by Bovio-Winkler et al., (2023).

**References**

1. Dueholm MKD, Nierychlo M, Andersen KS, Rudkjøbing V, Knutsson S, the MiDAS Global Consortium, Albertsen M, Nielsen PH, 2022. MiDAS 4: A global catalogue of full-length 16S rRNA gene sequences and taxonomy for studies of bacterial communities in wastewater treatment plants. Nat Commun 13:1908.

2. Bovio-Winkler P, Guerrero LD, Erijman L, Oyarzúa P, Suárez-Ojeda ME, Cabezas A, Etchebehere C. 2023. Genome-centric metagenomic insights into the role of Chloroflexi in anammox, activated sludge and methanogenic reactors. BMC Microbiol 23:45.
