## Supplementary Material - images and tables for "A comprehensive overview of the Chloroflexota community in wastewater treatment plants worldwide"

List of content:

**Figure S1.** Average mean abundance of the most abundant Chloroflexota genera in Danish WWTPs. Data is retrieved from the Danish MiDAS 3 survey (Nierychlo et al., 2020).

**Figure S2.** V1-V3 and V4 amplicon read abundance comparison for Chloroflexota phylum (first graph) and selected abundant genera. Data is retrieved from the global MiDAS survey (Dueholm et al., 2022) with total of 929 activated sludge plants with different process design. Grey diagonal line denotes equal V1-V3 and V4 abundances. For visualization purposes, samples with abundances higher than 10% are not shown in this figure.

**Figure S3.** Example of a Raman spectrum from the species *Ca*. Promineofilum glycogenicum, showing the presence of intracellular glycogen.

**Table S1**. Summary information about the climate zone division.

**Table S2**. Detailed summary of the probes designed and optimized in this study.

**Table S3.** Abundance estimation (percentage of total) performed by 16S rRNA amplicon sequencing (V1-V3 regions primer set) and qFISH.

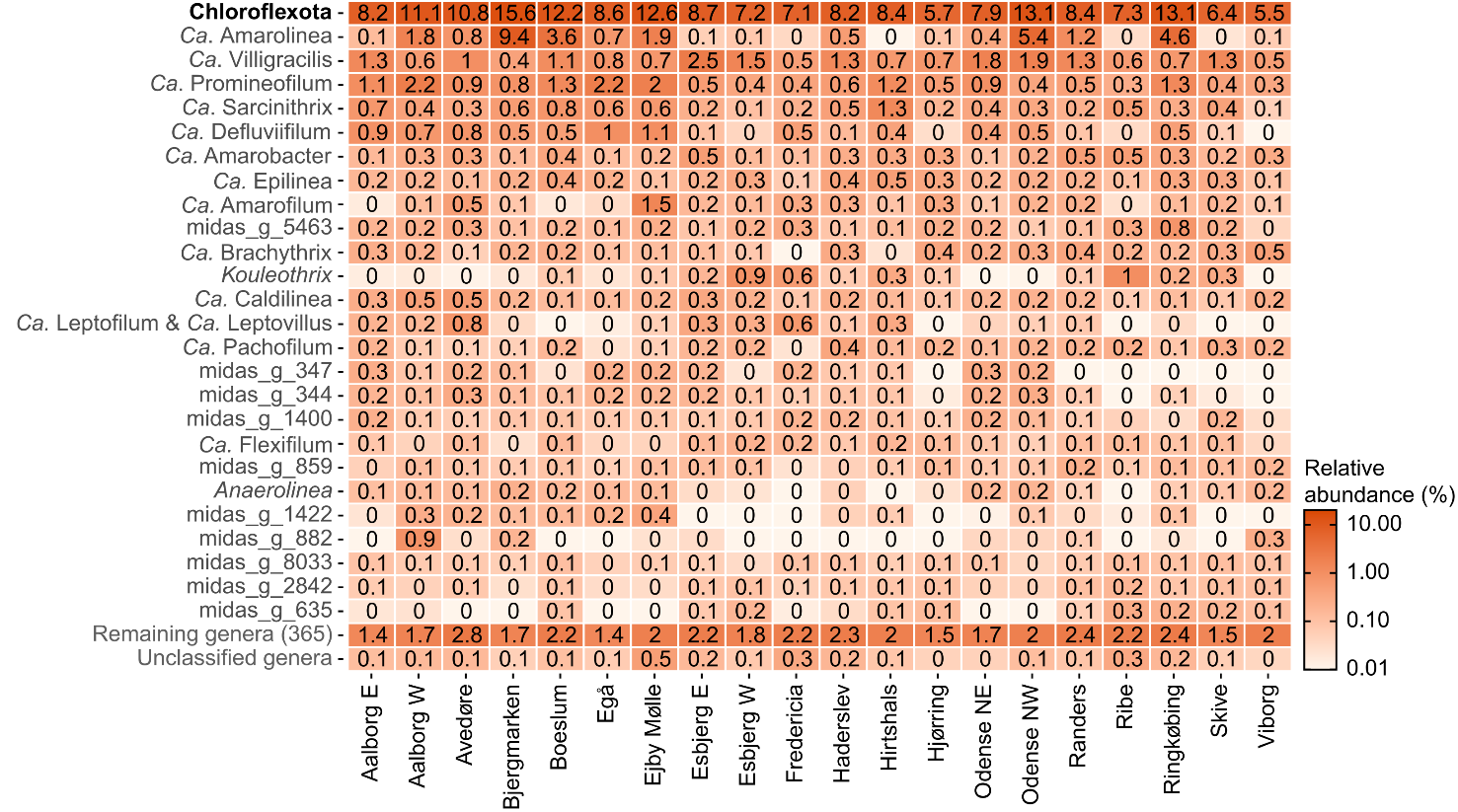

**Figure S1.** Average mean abundance of the most abundant Chloroflexota genera in Danish WWTPs. Data is retrieved from the Danish MiDAS 3 survey (Nierychlo et al., 2020).

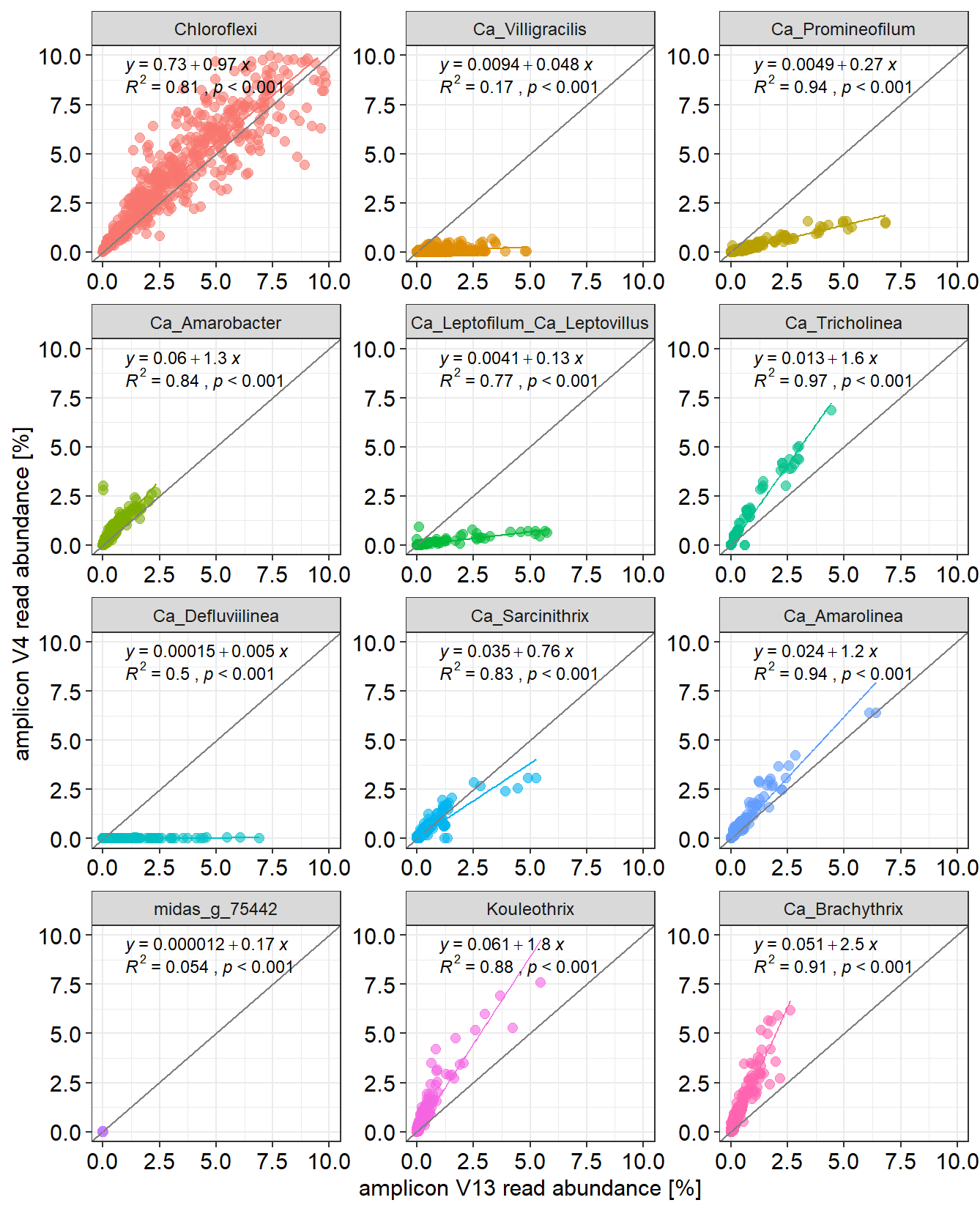

**Figure S2**. V1-V3 and V4 amplicon read abundance comparison for Chloroflexota phylum (first graph) and selected abundant genera. Data is retrieved from the global MiDAS survey (Dueholm et al., 2022) with total of 929 activated sludge plants with different process design. Grey diagonal line denotes equal V1-V3 and V4 abundances. For visualization purposes, samples with abundances higher than 10% are not shown in this figure.

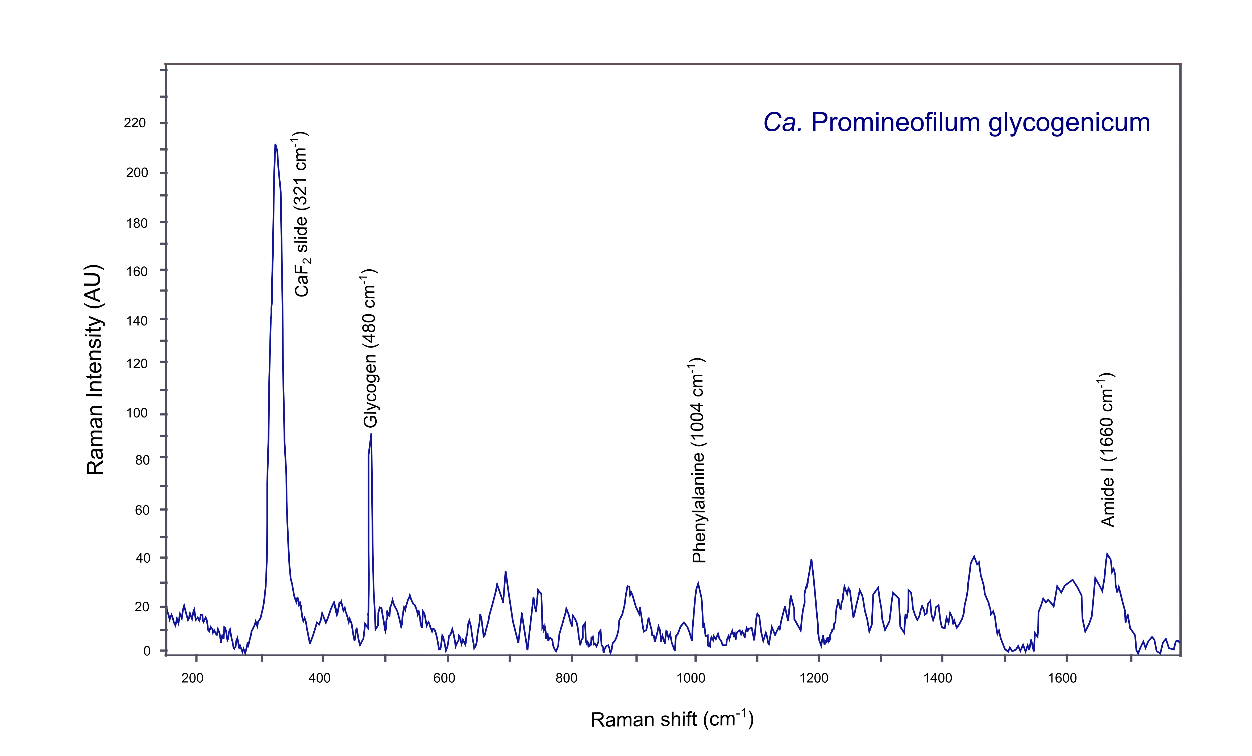

**Figure S3.** Example of a Raman spectrum from the species *Ca*. Promineofilum glycogenicum, showing the presence of intracellular glycogen.

**Table S1**. Summary information about the climate zone division.

| **Abbreviation** | **Climate** | **Countries** |
| --- | --- | --- |
| **A** | Tropical/mesothermal | India, Malaysia, Philippines, Singapore |
| **B** | Dry (desert and semi-arid) | Argentina, Australia, Canada, China, Israel, India, Mexico, Saudi Arabia, Spain, USA |
| **C** | Temperate/mesothermal | Argentina, Australia, Austria, Belgium, China, Cyprus, Czech Republic, Denmark, Germany, Hong Kong, Israel, Italy, Netherlands, Norway, Portugal, Poland, Spain, South Africa, Switzerland, Sweden, UK, USA, Uruguay |
| **D** | Continental/microthermal | Canada, China, Finland, Norway, Poland, South Africa, Sweden, USA |
| **E** | Polar | Switzerland |

**Table S2.** Detailed summary of the probes designed and optimized in this study.

| **Probe** | ***E. coli* pos.** | **Target group** | **Coverage**  **MiDAS4.8** | **Non-target hits** | **Sequence (5’-3’)** | **[FA]%** |
| --- | --- | --- | --- | --- | --- | --- |
| **CFX1111** | **1111-1132** | ***Ca.* Epilinea breve** | **17/20** | **0** | **CAC GTG AAA CAT ACG CCA AGG GT** | **40** |
| CFX1111_H1 | 1085-1106 | Helper for CFX1111 probe | N/A | N/A | GCG CTC GTT GCG GGA CTT AAC | N/A |
| CFX1111_H2 | 1135-1154 | Helper for CFX1111 probe | N/A | N/A | CGC CGG CAG TYG CGC ATG A | N/A |
| **CFX325** | **325-346** | ***Brachythrichaceae*** | **124/136** | **0** | **GTA GGC GTC TGG ACC GTG TTT** | **30** |
| CFX325_C1 | 325-346 | Competitor for CFX325 probe | N/A | N/A | GTA GGW GTC TGG ACC GTG TTT | N/A |
| CFX325_H1 | 299-320 | Helper for CFX325 probe | N/A | N/A | TCC TCT CAG AYC CCC TAC CCG | N/A |
| CFX325_H2 | 350-371 | Helper for CFX325 probe | N/A | N/A | TAT TCC TCM CTG CTG CCA CCC | N/A |
| **CFX198** | **198-219** | ***Ca.* Brachythrix** | **18/18** | **0** | **CCT CTC CTC ACG CCT TTC GAC** | **50** |
| CFX198_H1 | 170-194 | Helper for CFX198 probe | N/A | N/A | GAC CCT TTT GGG TAT TAG CCT CTC | N/A |
| CFX198_H2 | 230-249 | Helper for CFX198 probe | N/A | N/A | CTA GCT GAT GGG CCG CGG GCT | N/A |
| **CFX841_2** | **841-865** | ***Ca.* Trichofilum** | **33/33** | **0** | **AGC TAC AGC ACA GAG GGA TTG GAT** | **30** |
| CFX841_2_C1 | 841-865 | Competitor for CFX841_2 probe | N/A | N/A | AGC TAC AGC ACA GAG GGG TTG GAT | N/A |
| CFX841_2_C2 | 841-865 | Competitor for CFX841_2 probe | N/A | N/A | AGC TAC AGC ACA GAG GGA TTG GCT | N/A |
| CFX841_2_C3 | 841-865 | Competitor for CFX841_2 probe | N/A | N/A | AGC TAC AGC ACA GGG GGG TTG GAT | N/A |
| **CFX1086** | **1086-1110** | ***Flexifilaceae*** | **429/733** | **0** | **GCG CTC GTT TTC GGA CTT AAC CGA** | **30** |
| **CFX643** | **643-662** | ***Ca.* Flexifilum breve** | **116/119** | **0** | **TCC CAC TCT AGT CCC ACA G** | **30** |
| CFX643_C1 | 643-662 | Competitor for CFX643 probe | N/A | N/A | TCC CAC TCT AGT CCC GCA G | N/A |
| **CFX748** | **748-769** | ***Ca.* Leptofilum & *Ca.* Leptovillus** | **57/62** | **0** | **TTT CGC ATC TGA GCG TCA GGA** | **35** |
| CFX748_C1 | 748-769 | Competitor for CFX748 probe | N/A | N/A | TTT CGC ATC TGA GCG TCA GGT | N/A |
| CFX748_H1 | 720-738 | Helper for CFX748 probe | N/A | N/A | TGG CCC AGA GAG CCG CCT | N/A |
| CFX748_H2 | 740-760 | Helper for CFX748 probe | N/A | N/A | ATC CYG TTC TCT CCC CTA GC | N/A |
| **CFX1194** | **1194-1216** | ***Tepidiformales*** | **132/159** | **0** | **CGT AAG GGC CAC GCT GAC CTG A** | **50** |
| CFX1194_H1 | 1170-1190 | Helper for CFX1194 probe | N/A | N/A | TCG TCC CCT CCT TCC TCC GA | N/A |
| CFX1194_H2 | 1220-1241 | Helper for CFX1194 probe | N/A | N/A | GTA GCG TGT GTG TAG CCC CAG G | N/A |
| **CFX193** | **193-215** | ***Ca.* Amarobacter** | **33/104** | **0** | **TAG CGC CGG AGC TTT TAC CAC C** | **35** |
| CFX193_H1 | 168-192 | Helper for CFX193 probe | N/A | N/A | GGG TGT TAT GCG GTA TTA GCT CGC | N/A |
| CFX193_H2 | 220-240 | Helper for CFX193 probe | N/A | N/A | AGC TAA TCG GCC GCG GGC CC | N/A |
| **CFX122** | **122-141** | ***Ca.* Amarofilum** | **5/6** | **0** | **CTT GGG CAC ATT CCC ACG T** | **35** |
| CFX122_C1 | 122-141 | Competitor for CFX122 probe | N/A | N/A | TTT GGG CAC ATT CCC ACG C | N/A |
| **CFX682** | **682-706** | ***Ca.* Pachofilum** | **55/55** | **0** | **ATC TAC ATA TTC CAC CAT TAC ACC** | **35** |
| CFX682_H1 | 652-672 | Helper for CFX682 probe | N/A | N/A | TCC GCA TTC CTC TCA TYG CC | N/A |
| CFX682_H2 | 710-732 | Helper for CFX682 probe | N/A | N/A | CTT TCG CAC ATG AGC GTC AGG C | N/A |
| **CFX166** | **166-189** | ***Ca.* Tricholinea** | **14/17** | **0** | **GTA ACY TCA TGC GGT ATT AGC AG** | **35** |
| CFX166_H1 | 138-160 | Helper for CFX166 probe | N/A | N/A | GCA GGT CAC CAA CGC GTT ACT C | N/A |
| CFX166_H2 | 195-218 | Helper for CFX166 probe | N/A | N/A | GCT GAT GGG ACG CAG GCC CCT CC | N/A |
| **CFX1423** | **1423-1446** | ***Ca.* Defluviifilum** | **31/116** | **0** | **GAG TCA CCG ACT TCA GGT GTT CC** | **50** |
| CFX1423_C1 | 1423-1446 | Competitor for CFX1423 probe | N/A | N/A | AAG TCA CCG ACT TCA GGT GTT CC | N/A |
| CFX1423_C2 | 1423-1446 | Competitor for CFX1423 probe | N/A | N/A | AAG TCA CCG ACT TCA GGT GTT TC | N/A |

**Table S3.** Abundance estimation (percentage of total) performed by 16S rRNA amplicon sequencing (V1-V3 regions primer set) and qFISH.

| **WWTP** | **Sample date** | **Abundance (%)** | |
| --- | --- | --- | --- |
|  |  | **Sequencing** | **qFISH** |
| ***Ca.* Epilinea** | | | |
| Aars | August 2008 | 2.4 | 1.3 ± 0.3 |
| Boeslum | August 2007 | 1.4 | 1.6 ± 0.4 |
| ***Brachytrichaceae*** | | | |
| Oud-Turnhout (Belgium) | March 2018 | 2.2 | 6.6 ± 3.0 |
| Merelbeke (Belgium) | March 2018 | 2.1 | 2.2 ± 1.4 |
| ***Ca.* Brachythrix** | | | |
| Oud-Turnhout (Belgium) | March 2018 | 1.7 | 5 ± 1.2 |
| Merelbeke (Belgium) | March 2018 | 1.8 | 1.8 ± 0.5 |
| ***Ca.* Trichofilum**  Randers  Aars | August 2012  August 2008 | 0.6  0.5 | 0.8 ± 0.3  2.0 ± 0.8 |
| ***Flexifilaceae***  Fredericia  Kalundborg | August 2016  September 2017 | 2.6  2.8 | 1.1 ±0.5   < 0.5 |
| ***Ca.* Flexifilum breve**  Esbjerg W  Hirtshals | August 2016  Hirtshals 2010 | 0.5  0.4 | < 0.5  < 0.5 |
| ***Ca.* Leptofilum & *Ca.* Leptovillus** | | | |
| Avedøre | August 2010 | 1.6 | 5.1 ± 1.5 |
| Aars | August 2007 | 0.8 | 1.6 ± 1.3 |
| ***Tepidiformales*** | | | |
| Randers | August 2010 | 1.2 | < 0.5 |
| Marselisborg | August 2006 | 1.0 | < 0.5 |
| ***Ca.* Amarobacter** | | | |
| Randers | August 2007 | 1.0 | < 0.5 |
| JeddahA (Saudi Arabia) | May 2018 | 3.0 | 1.1 ± 0.5 |
| ***Ca.* Amarofilum** | | | |
| Ejby Mølle | August 2008 | 8.3 | 3.8 ± 2.7 |
| Aars | August 2006 | 1.0 | 1.4 ± 1.3 |
| ***Ca.* Pachofilum** | | | |
| Aars | August 2009 | 2.8 | 2.4 ± 1.1 |
| Aars | August 2006 | 1.9 | 1.6 ± 0.8 |
| ***Ca.* Tricholinea** |  |  |  |
| Dahej (India) | July 2018 | 9.0 | 6.1 ± 1.6 |
| ***Ca.* Defluviifilum** |  |  |  |
| Åby | August 2007 | 1.5 | 9.9 ± 2.5 |
| Ejby Mølle | August 2018 | 1.2 | 5.2 ± 1.5 |
